# Heterologous Reconstitution of 4-Amidopentadienoate Biosynthesis Identifies the PKS Machinery Responsible for Warhead Formation

**DOI:** 10.64898/2026.09.17.752105

**Authors:** Marie S. Tvilum, Jonas L. Veiler, Thomas T. Paulsen, Carsten Kegler, Camilla B. Nielsen, Mogens Johannsen, Thomas B. Poulsen, Helge B. Bode, Thomas Tørring

## Abstract

The 4-amidopentadienoate (APD) motif is a rare and bioactive pharmacophore found in the cyclic lipodepsipeptide natural products BE-43547, rakicidins, microtermolides, and related compounds. Although several biosynthetic gene clusters associated with this family have been identified, the enzymatic origin of the APD warhead has remained unknown. Here, we investigated APD formation using NRPS engineering to install the unusual APD-associated polyketide synthase (PKS) module from the BE-43547 biosynthetic pathway into heterologous nonribosomal peptide synthetase/polyketide synthase (NRPS/PKS) assembly lines expressed in *Escherichia coli*. The engineered hybrids produced the designed lipopeptide containing the APD moiety, demonstrating that the core biosynthetic machinery is sufficient for APD installation and that no trans-acting enzymes are required. Total synthesis of the target compound enabled structural validation by co-elution, MS/MS analysis, and NMR spectroscopy. Production titers of 59 ± 13 μg L^−1^ were achieved in the heterologous host. Systematic mutagenesis further revealed that the conserved dehydratase His-Asp catalytic dyad is essential for APD formation, whereas an additional highly conserved histidine is dispensable. These findings establish the biosynthetic origin of the APD warhead and provide a foundation for incorporating this privileged pharmacophore into engineered peptide scaffolds.

## Introduction

Natural products continue to inspire the development of new leads in the pharmaceutical industry. They provide an extensive source of structural diversity, as well as new modes of action. One such example is the 4-amidopentadienoate cyclic lipodepsipeptide (APD-CLD) natural product family, including microtermolides, rakicidins, boholamide, and many others (Figure S1).^[1–4]^ Arguably, the most prominent member is BE-43547, which consists of seven congeners that have demonstrated potent hypoxia-selective cytotoxicity across multiple cell lines.^[5–7]^ We have recently shown that this selectivity is also reflected in their antimicrobial activity.^[8]^ Under anaerobic cultivation conditions, these compounds act as highly potent antibiotics against the Gram-positive pathogen *Staphylococcus aureus*, with a minimal inhibitory concentration (MIC) of 0.06 µg/mL as well as a modest bioactivity against biofilms and persister cells. Previous studies have established that the APD moiety functions as the primary biochemical warhead and is therefore essential for bioactivity.^[6,9–12]^

While we and others have connected biosynthetic gene clusters (BGCs) to several members of the APD-CLDs, the installation of the APD warhead has never been investigated. The predominant hypothesis is that the core biosynthetic machinery within the polyketide synthase or non-ribosomal synthetase is somehow involved, since no gene for a free-standing enzyme appears to be conserved across all BGCs associated with the APD-CLD family. All APD-CLD BGCs are hybrid systems composed of non-ribosomal peptide synthetases (NRPSs) and type I polyketide synthases (PKSs). Because these BGCs are modular, the biosynthesis and assembly of the corresponding natural products can be predicted assuming a co-linearity principle, where each module is responsible for elongating the growing product one building block at a time and in the order dictated by the arrangement of the genes in the genome. The two types of biosynthetic modules, NRPS and PKS, encode different domains with predictable and specific enzymatic activity. Within the NRPS, an adenylation (A) domain selects and activates the extender unit, which is then covalently transferred onto the phosphopantetheine arm, attached to a thiolation (T) domain. The condensation (C) domain then facilitates the linking between the growing peptide chain and the extender unit. Mirroring domains and mechanisms are found within the PKS, having the acetyltransferase (AT) and ketosynthase (KS) domains as equivalents to the A and C domains, respectively. Additionally, multiple tailoring domains can be present, such as the N-methyltransferase (MT) in the NRPS, and the ketoreductase (KR), dehydratase (DH), and enoylreductase (ER) in the PKS. The final hybrid product can be released by either a terminal C domain or a thioesterase (TE) domain via hydrolysis or macrocyclization. The T domain is a general term encompassing both the acyl carrier protein (ACP) from the PKS and the peptidyl carrier protein (PCP) from the NRPS. Since the discovery of these modular assembly lines more than 30 years ago, researchers have envisioned the design of rationally designed molecules by employing the modules in a LEGO-like fashion. Recently, large strides towards this vision have been demonstrated using the “exchange unit between T domains” (XUT) approach.^[13]^ This has also allowed for the generation of peptide-polyketide hybrids.^[14]^ XUT facilitates the assembly of NRPS or NRPS and PKS units at conserved fusion points in the middle of or directly before the thiolation (T) domain.^[15]^ Inspired by this, we hypothesized that the biosynthesis of the APD warhead could be investigated by transplanting modules from the BE-43547 BGCs into simpler constructs and paving the way for incorporating potent warheads into rationally designed molecules.

## Results and Discussion

### PKS Module from the APD-CLD BGC Installs the APD Warhead

Based on bioinformatic predictions on the genomes of producing microorganisms, we have previously proposed the BGC shown in Figure 1a as responsible for the BE-43547 congeners.^[5,8]^ Assembly of the non-ribosomal peptide/polyketide (NRP/PK) hybrid is initiated by loading an acyl chain, a step that determines which BE-43547 congener is produced depending on the length and branching of the starter unit. Notably, all seven congeners are produced by both *Salinispora arenicola* CNR107 and *Micromonospora* sp. RV43, and we have therefore illustrated the assembly line using BE-43547A_1_ as a representative. It has previously been proposed that the tailoring domains within PKS module six, which contains an unusual order of the KR-DH didomain, are responsible for installing both the canonical α,β- and the non-canonical γ,δ-double bonds.^[5,16]^ This requires serine to be installed by the preceding NRPS module five, which is well in line with the prediction made by antiSMASH^[17]^ and earlier feeding experiments using [1-^13^C]-L-serine in the rakicidin D producer *Streptomyces* sp. MWW064.^[16]^

**Figure 1.**
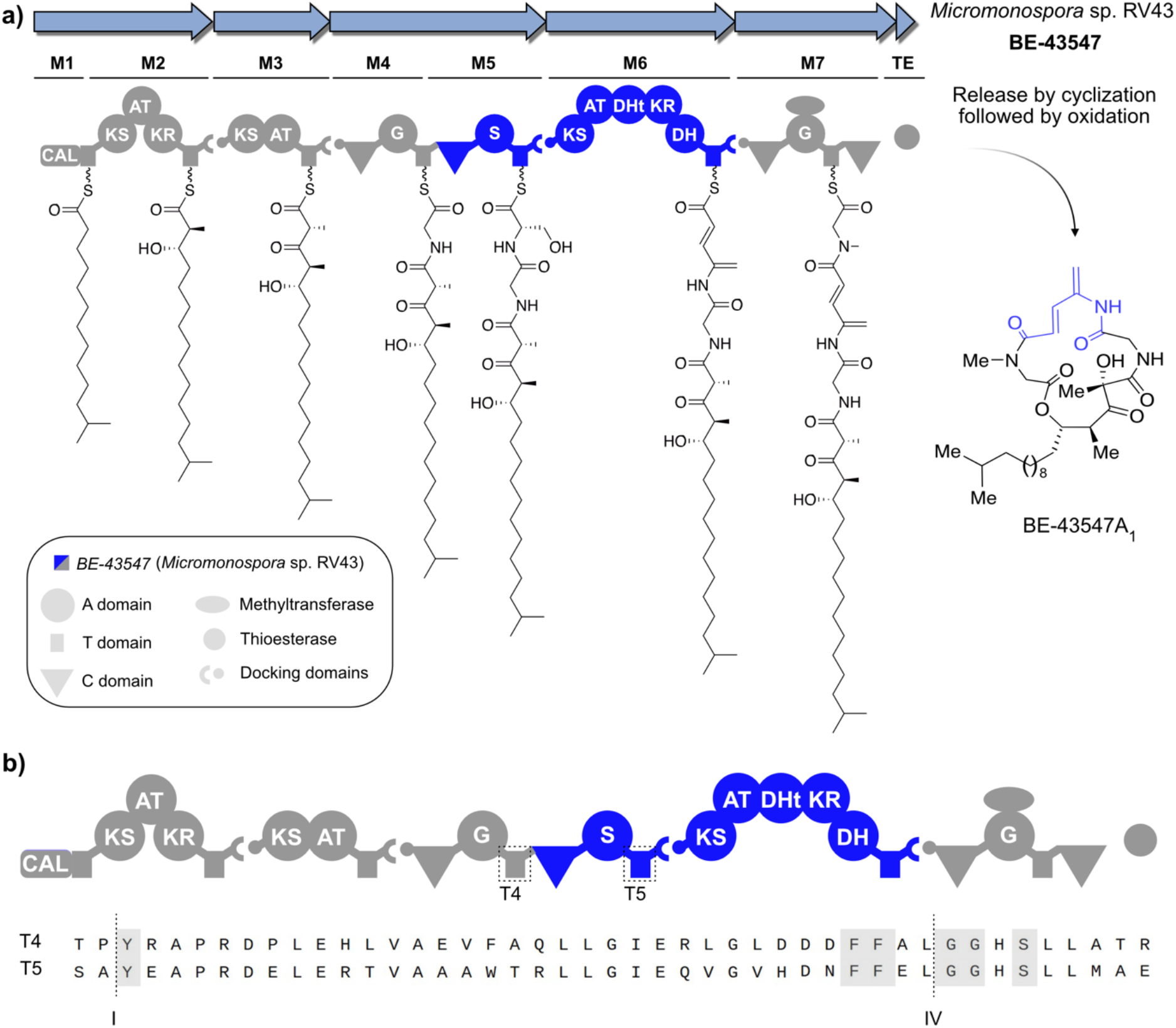
Biosynthetic gene cluster and biosynthesis of the BE-43547 from Micromonospora sp. RV43. a) The biosynthesis of BE-43547 is illustrated using the A_1_ congener, where blue indicates the modules implementing the APD moiety as well as the APD moiety on the final structure. The BGC encodes the biosynthesis of all seven congeners, as they only differ in length and branching of the lipophilic tail implemented through the CAL domain. b) Amino acid sequence of thiolation domains 4 and 5 from modules 4 and 5. The fusion sites are indicated for both XUT^I^ and XUT^IV^ along with important conserved amino acids as indicated in grey. CAL: Coenzyme-A ligase, KS: Ketosynthase, AT: Acyltransferase, KR: Ketoreductase, DHt: Dehydratase mainly found in trans-AT PKS, DH: Dehydratase.

After several failed attempts to genetically manipulate the BGC in *Micromonospora* sp. RV43, we sought to express the APD warhead in a heterologous expression system to investigate the unusual PKS module. The XUT approach has proven to be quite versatile with respect to the assembly of NRPS and even PKS parts from different organisms, including *Streptomyces*, myxobacteria, and even fungi.^[13–15]^ Since PKS module six from the BE-43547 BGC is flanked by two NPRS modules (module five incorporating L-Ser and module seven incorporating N-Me glycine, Figure 1a), we were quite optimistic that XUT could also be applied here to generate simplified peptide-polyketide hybrids. An important design parameter was to use a suitable starter module that is compatible with expression in the heterologous host.^[13]^ Accordingly, we decided to fuse an alternative starter module to the NRPS/PKS hybrid system.

Using antiSMASH, we identified all thiolation domains upstream of module six as either PCPs or ACPs. Here, we identified two domains as PCPs containing suitable fusion sites for the XUT approach (Figure 1b). Both domains contain the conserved tyrosine residue and the characteristic core motif (FFxxGGxS), enabling the use of fusion sites I and IV.

After identifying fusion sites in the BE-43547 BGC, we screened different starter modules to assess their suitability for heterologous expression. We used *Escherichia coli* DH10B::*mtaA* as the heterologous host because it expresses the phosphopantetheinyl transferase MtaA, which is required for the formation of functional thiolation domains.^[18– 20]^ Due to the high GC content (~74%) found in *Micromonospora* sp. RV43 and a codon usage that differs substantially from that of *E. coli*, we included codon-optimized constructs as well. The β-hydroxy starter unit from the xefoampeptide-producing NRPS XfpS has been used previously,^[13]^ and we therefore decided to fuse this to our serine module 5. However, incorporation of the serine module was unsuccessful, regardless of codon optimization or fusion site selection, as only compound 1 was detected upon expression of NRPS-2, -3a, and -3b (Figure 2a). In contrast, successful expression of NRPS-4a and 4b, which contain an alternative serine module derived from a xenolindicin-related starting module XldAB with Ser A1 domain from *Xenorhabdus stockiae* DSM 17904, resulted in the production of compound **2**.

**Figure 2.**
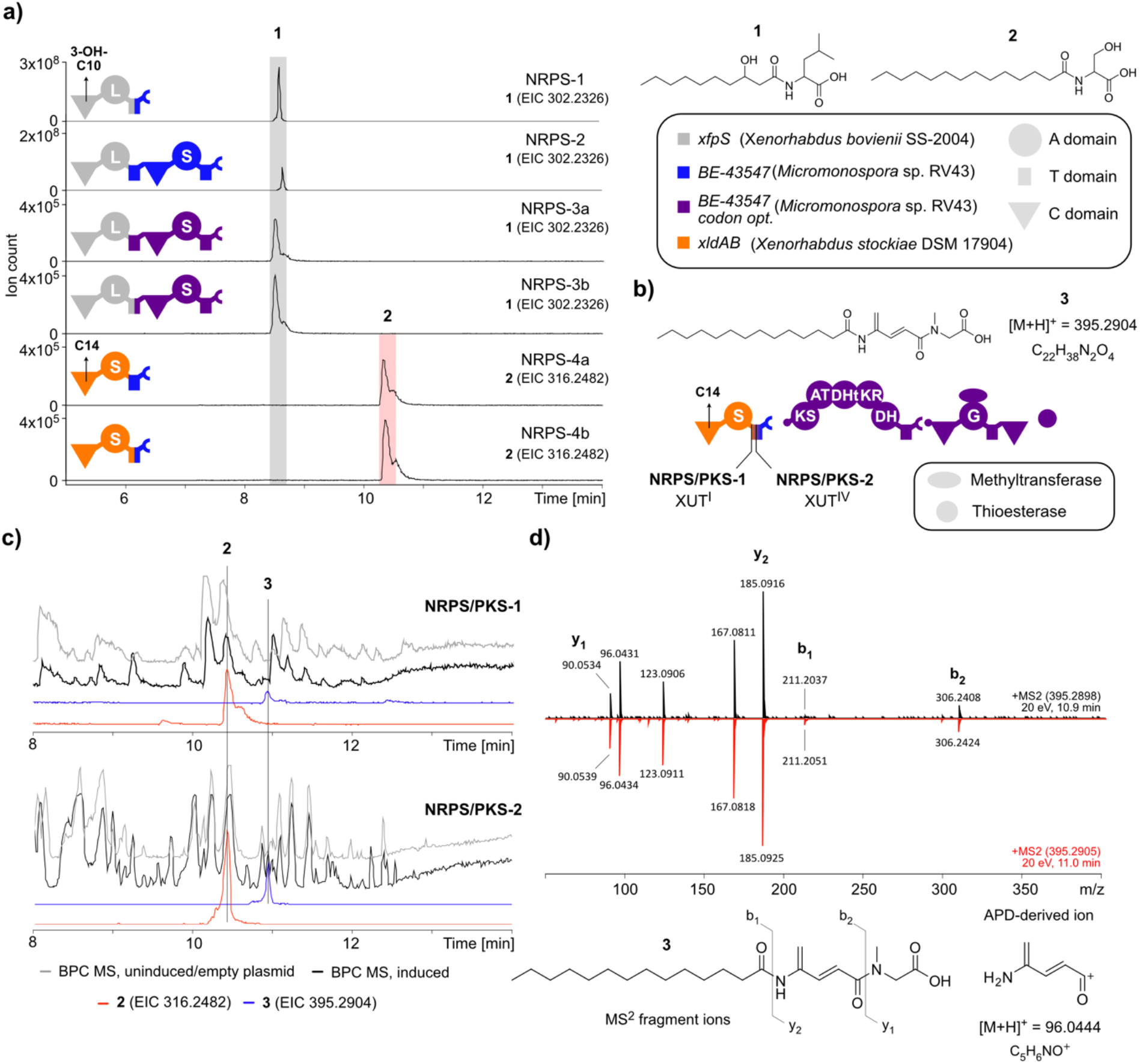
Heterologous expression of NRPS/PKS-hybrids. a) Different starter unit constructs and their resulting chemical structures observed. The initial C domains implement either a 3-OH-C10 or a C14 initially, as indicated. b) NRPS/PKS-1 and -2 hybrid systems, altering depending on whether NRPS-4a (−1) or NRPS-4b (−2) is used as the starter unit. c) HPLC-MS chromatograms from NRPS-PKS-1 and -2, using either an uninduced culture (NRPS/PKS-1) or a starter unit + empty plasmid (NRPS/PKS-2) as a reference of background signals (grey). The base peak chromatograms (BPC) are shown (black) along with extracted ion chromatograms (EICs) of **2** (*m/z* = 316.2482, red) and **3** (*m/z* = 395.2904, blue). d) MS^2^ fragmentation pattern of **3** extracted from NRPS/PKS-1 (black, top) and - 2 (red, bottom) at retention time 10.9-11.0 minutes.

To simplify the final constructs, we kept the downstream N-methylglycine-incorporating module seven, including its terminal condensation domain and thioesterase. This design yielded two final constructs, NRPS/PKS-1 and NRPS/PKS-2, both of which produced the desired product **3** (Figure 2b,c). Notably, compound **2** was also detected in both cases. Analysis of the MS^2^ fragmentation patterns revealed the presence of an APD-derived ion (*m/z* 96.04, consistent with C_5_H_6_NO^+^), indicating that the APD warhead was successfully installed by the PKS tailoring enzymes (Figure 2d). When coupling NRPS-1 directly to the downstream modules, we did not find production of any peptides other than **1**, indicating substrate selectivity of the PKS module (Figure S2).

### Synthetic Standard Confirms the APD Warhead-containing Product

To validate the proposed product, estimate production titers, and enable bioactivity testing, we designed a synthetic route inspired by a previous total synthesis of the rakicidin macrocyclic core.^[21]^ The route, shown in Figure 3, utilizes a pivotal Horner-Wadsworth-Emmons (HWE) reaction to assemble the APD moiety from two building blocks, phosphonate ester **7** and a,b-unsaturated aldehyde **8**. To prepare **7**, Boc-protected *N*-methylglycine was first esterified followed by Boc-deprotection and a-bromoacetylation with acylbromide **5**. Arbuzov reaction then delivered **7** in high yield. Aldehyde **8** was generated in a two-step sequence by initial EDCI-mediated coupling of serinol to myristic acid to give **9** followed by a cascade oxidation-dehydration reaction under Swern-conditions. Despite of the seemingly sensitive nature of **8**, the compound could be purified by a quick silica-plug. Optimized conditions for HWE-coupling of **7** and **8** were identified using stoichiometric potassium *tert*-butoxide as the base at low temperature (−78 °C) with drop-wise addition of **8**. This procedure cleanly delivered **10** in 75% yield (4:1 mixture of rotameric isomers) following silica-column purification. Finally, hydrolysis of the methyl ester using lithium hydroxide in a mixed solvent system (THF/H_2_O) delivered **3**.

**Figure 3.**
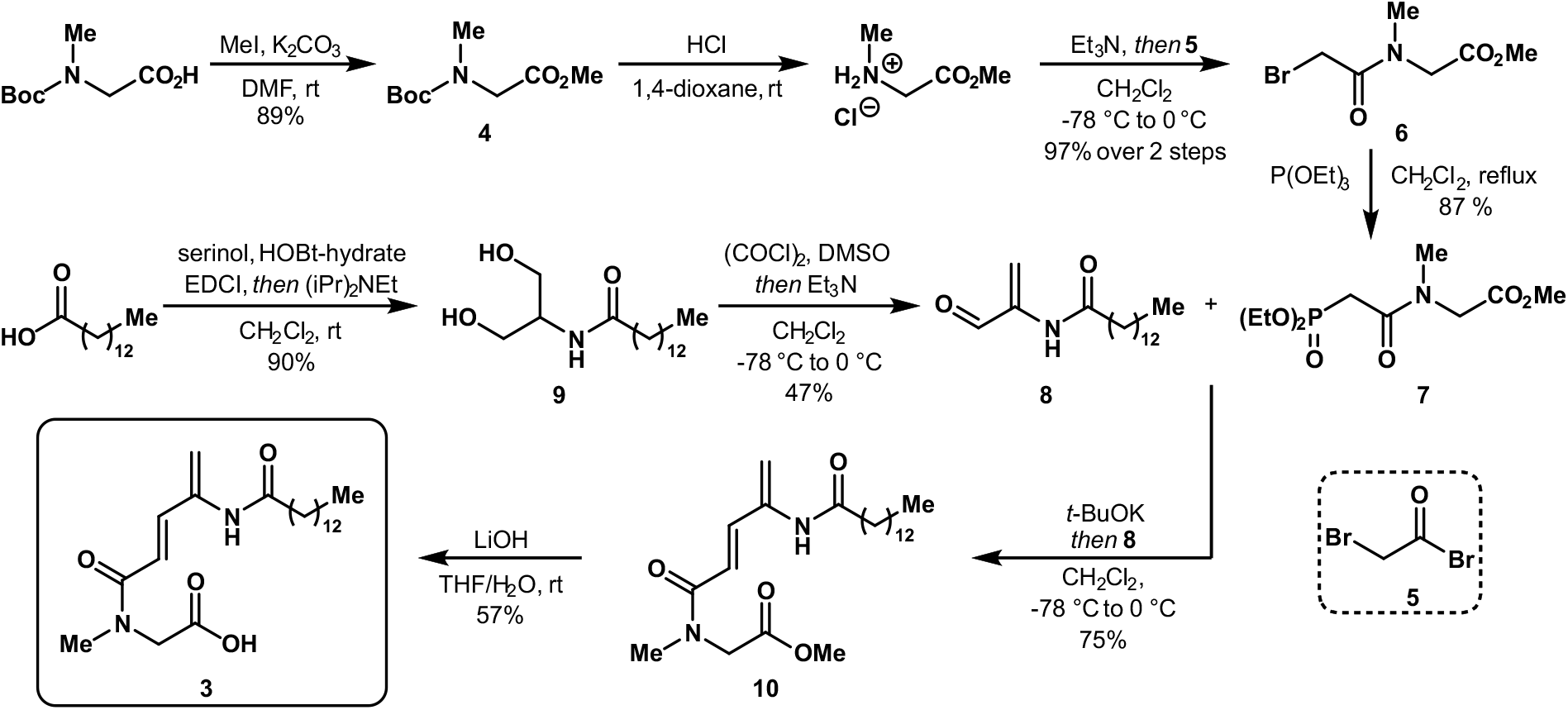
Synthetic route to **3.** Abbreviations: DMF = N,N-dimethylformamide; DMSO = Dimethylsulfoxide; HOBt = 1-hydroxybenzotriazole; EDCI = 1-Ethyl-3-(3-dimethylaminopropyl)carbodiimide; THF = Tetrahydrofuran. For full characterization data and copies of NMR spectra, see Supporting Information.

When comparing the biological extract containing the expressed NRPS/PKS-2 hybrid with the synthetic standard, we observed an identical retention time of the *m/z* corresponding to **3**. Further, we found co-elution when spiking the synthetic compound in the prepared extracts (Figure 4a). Comparison of the MS^2^ fragmentation pattern also revealed similar fragments. In Figure 4b, two ionization energies are applied and summarized (20 eV and 40 eV). This, in combination with comprehensive NMR characterization of both **10** and **3**, leads us to conclude that the designed hybrid assembly line works as intended. Furthermore, it highlights that module six from the BE-43547 BGC is responsible for the installation of the APD warhead and that no other enzymes from *Micromonospora* sp. RV43 work *in trans* on the assembly line to install it.

**Figure 4.**
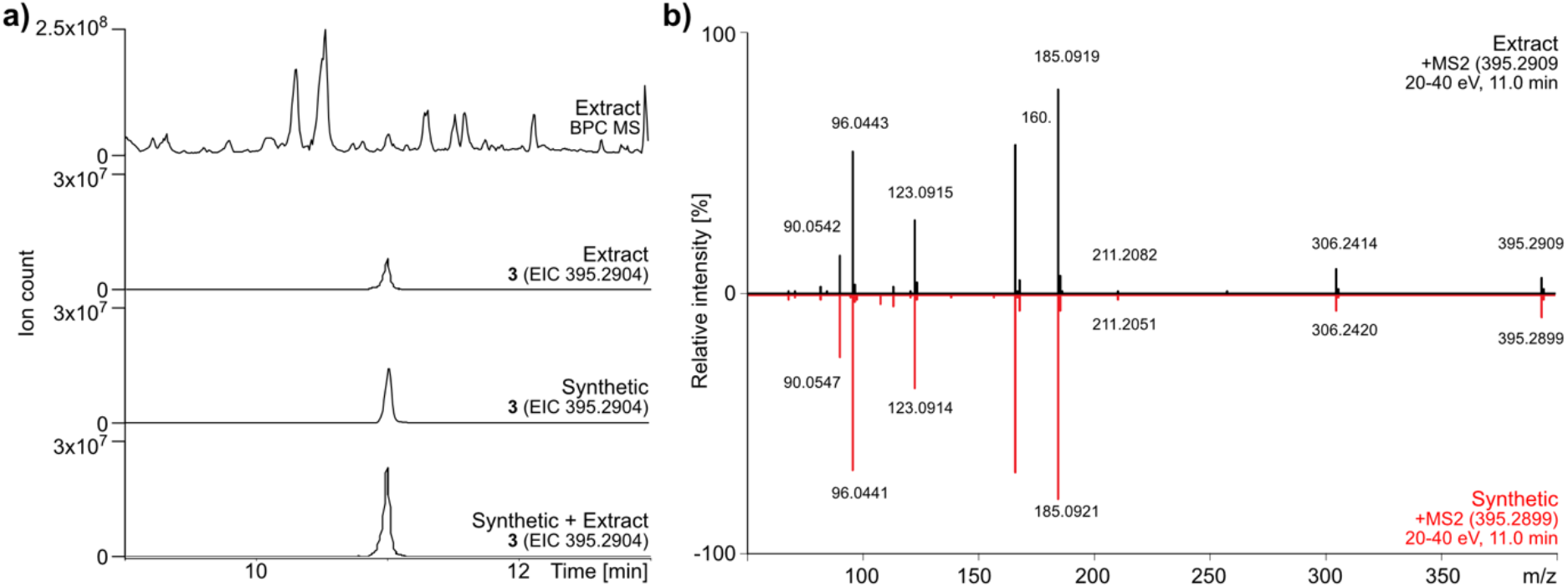
Comparison of synthetic **3** with a biological extract from the NRPS/PKS-2 heterologous expression. a) HPLC-MS of the biological extract (top) with its EIC (*m/z* = 395.2904) below. The EICs are shown for the synthetic compound **3** and the spiked sample (Synthetic + Extract) in the two bottom figures. The spiked sample contains both the synthetic compound and the biological extract. b) MS^2^ fragmentation patterns of **3** (*m/z* = 395.2904) from the biological extract (top, black) and the synthetic compound (bottom, red).

Using the synthetic standard, we could quantify the amounts of **3** expressed by the *E. coli* containing the NRPS/PKS-2 hybrid. From UHPLC-UV signals, we found that the *E. coli* produced 59 ± 13 µg/L. In previous work, titers between 0.05 and 75.88 mg/L have been observed for NRPS constructs via XUT IV fusion site, showing a tendency of a lowering in yield when fusing a high-GC NRPS into a low-GC NRPS system. Although this method has not proven to yield high productivity, this first step of moving a natural product warhead from its native setting and into a heterologous system could provide multiple new opportunities for designer peptides in a cost- and time-efficient approach. Further evaluations of the warhead were done by bioactivity assays of the synthetic standards **3** and **10**. To this, we included bacterial extracts from the NRPS/PKS-2 culture broth and BE-43547 as a positive control. All were tested against *Staphylococcus aureus* DSM 20231 by spotting extracts and isolated compounds onto a lawn of the pathogen, which was grown either aerobically or anaerobically (Figure S3). We found no bioactivity of **3** and **10**, implying that the additional strain imposed in the original cyclic molecule is important for the biological function of the warhead. Having established a minimal biosynthetic construct for installing the APD warhead, we sought to investigate the enzymatic mechanisms responsible.

### Catalytic Dyad is Essential for APD Formation

Inspection of the BGC assembly line revealed a non-canonical ordering of the KR–DH didomain, suggesting an atypical mode of interaction between the KR and DH domains compared with canonical modular PKS systems. This is rare among other PKS BGCs, but conserved across all known producers of APD-CLDs. Dehydratase domains have in several other assembly lines been found to catalyze reactions diverting from hydration leading to the canonical α,β-double bond formation. In curacin biosynthesis, two DH domains (CurJ-DH and CurH-DH) were shown to catalyze both installation of α,β- and γ,δ-double bonds using the same active site.^[22]^ Similarly, it has been demonstrated that a single DH domain from the pure polyketide gladiolin BGC in *Burkholderia gladioli* BC0238 is capable of catalyzing a double dehydration of a (3*R*, 5*S*)-3,5-dihydroxyacyl thioester bound to an ACP and yielding a (2*E*, 4*Z*)-2,4-dienoyl thioester.^[23]^ The researchers also found that it went through a (2*E*, 5*S*) intermediate, demonstrating that the α,β-elimination precedes the γ,δ-elimination. A comprehensive bioinformatic analysis showed that an additional histidine (H158) was found in the active site and universally conserved in PKS modules associated with diene compounds.^[23]^

We therefore focused on elucidating the function of the DH domain. By aligning sequences from all microorganisms harboring APD–CLD BGCs that we have shown to be producing, we identified the conserved His-Asp catalytic dyad responsible for α,β-double bond formation (Figure S4). This dyad consists of a histidine residue (H10, orange) within the HxxxGHGxxP motif and an aspartic acid residue (D186, red) within the HPALLD motif (Figure 5).^[24,25]^ In addition, we identified a second histidine (H15, grey), conserved among all producers and located five residues downstream of the catalytic histidine. Based on the work on the gladiolin BGC,^[23]^ we hypothesized that a similar mechanism may occur. However, AlphaFold-based structure prediction^[26]^ did not support this hypothesis, as the second histidine (H15) was positioned on the surface of the dehydratase and distant from the predicted active site (Figure S5).

**Figure 5.**
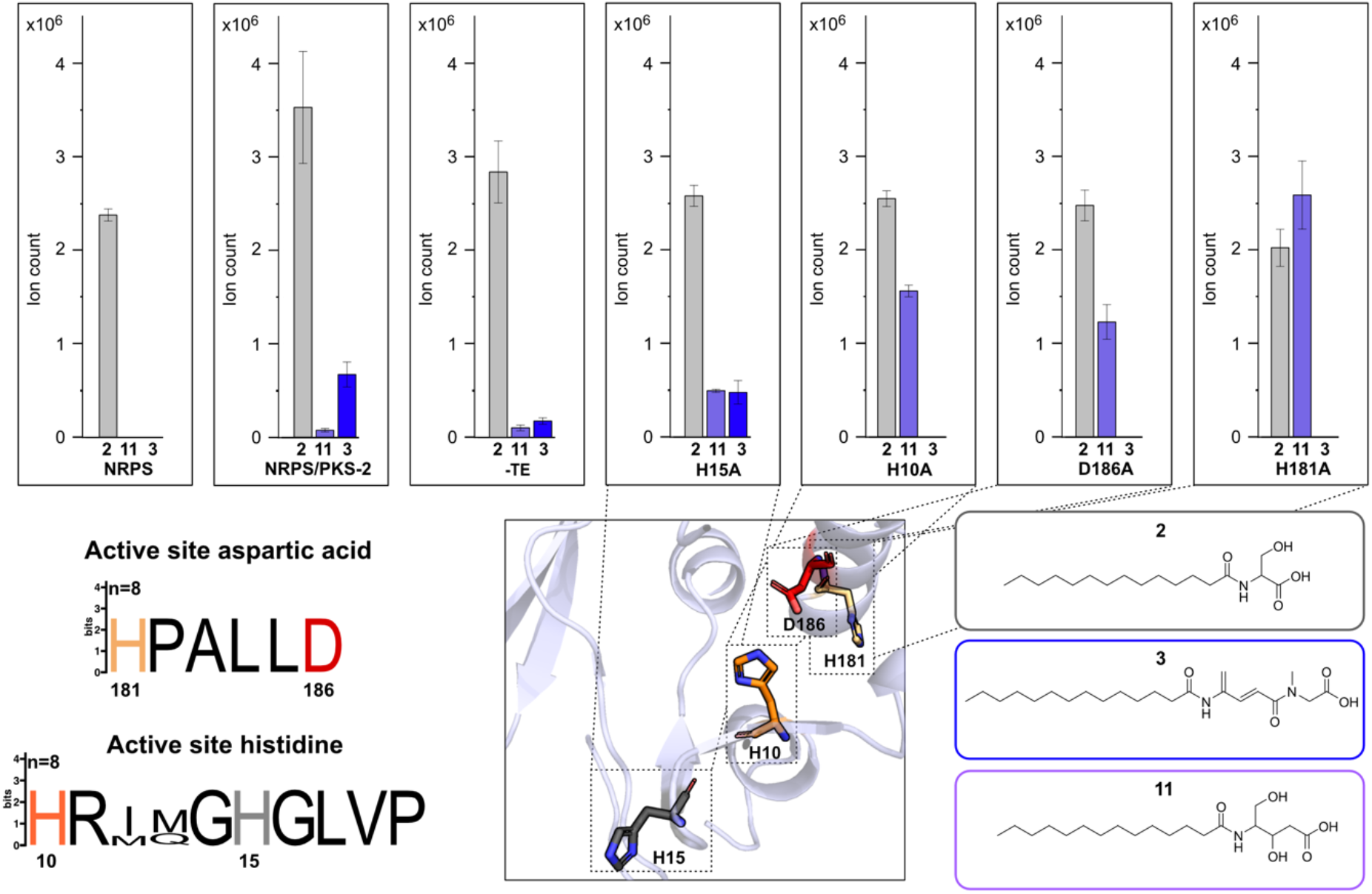
Heterologous expression of NRPS/PKS-2 mutants. Production titers are based on the ion count of the MS area using HPLC-MS (n=3 ±SD). NRPS: NRPS-4b + empty plasmid. -TE: NRPS/PKS-2 without the thioesterase. Compound **2** is from the starter unit (grey bar), **3** is the NRPS/PKS-2 product containing the APD moiety (blue bar), and **11** is created after the implementation of the malonyl building block, with the ketone from the serine reduced, followed by release from the assembly line (purple bar). The active site motif of the non-canonical DH of all APD-CLD producers we have validated is shown with numbering based on *Micromonospora* sp. RV43. The AlphaFold model of the active site of the dehydratase from *Micromonospora* sp. RV43 is shown, with the altered amino acid residues shown in sticks (H10: orange, H15: grey, H181: light orange, D186: red).

We next set out to experimentally validate the function of the identified amino acid residues through multiple point mutations within the DH domain (Figure 5). Here, the NRPS-4b + empty plasmid (NRPS) only produces the starter unit **2** (grey) compared to the NRPS/PKS-2 production, which is able to produce both the starter unit **2** and the APD-containing product **3** (blue). Similar productions are observed for the construct in which the thioesterase has been removed, indicating that the C domain is the one functioning during the release of the NRP/PK hybrid. However, the thioesterase could function as a proofreading mechanism, as the titers of **3** are not equally high. As we altered the residues H10, D186, and H181 to alanine separately, the production of **3** vanished in all mutants (H10A, D186A, H181A) (Figure 5). Additionally, we identified a mass feature matching the expected mass of the reduced form of the fatty acid-serine-malonyl peptide **11** (*m/z* = 360.2744). This was further validated with the MS^2^ fragmentation pattern (Figure S6-7). The reduced form would occur after the ketoreductase reduced the carbonyl incorporated with the serine residue, followed by the release through hydrolysis of the peptide from the assembly line. We see an increase in a mass feature matching the expected mass of compound **11** when performing point mutations, indicating that the ketoreductase functions first, prior to any elimination of the hydroxy groups. This is also the canonical order of ketoreductase and dehydratase functions. When changing the non-canonically, but highly conserved, histidine residue H15 to an alanine residue, we retain production of **3** (H15A). Thus, as suspected after inspecting the AlphaFold structure, H15 is not responsible for the second elimination of the serine hydroxy group (Figure S4-5).

Based on these results, we expanded the search for amino acid residues involved in dehydration reactions within the DH domain and identified five additional histidine residues, along with six other residues of interest (Figure S4). The latter were either found in the vicinity of the active site or could somehow help facilitate a second elimination by affecting the confirmed catalytic active His-Asp dyad. Common for all alanine-mutants was that we never detected the single-dehydrated product, suggesting that dehydration must occur fully or not at all. We did, however, find increased levels of **11** when altering amino acid residues near the active site, similar to the ones obtained when altering the catalytically active residues (Figure S8-10). Finally, we also considered whether the DHt domain found directly upstream of the ketoreductase was catalytically active. From point mutations in all histidine residues present, we did not find any indications of the DHt domain being catalytically involved in the formation of **3** (Figure S11-14).

Based on the results obtained, we conclude that the dehydratase plays a crucial part during the formation of the APD moiety. We hypothesize a reaction mechanism in which the first step is the reduction of the ketone group from the serine residue facilitated by the KR domain, followed by the canonical function of the DH domain, which eliminates the hydroxy-group at the β-position, creating a C-C double bond at the α,β-position. We further hypothesize this position to be eliminated first, due to the relative acidity of the protons on the α carbon compared to the γ carbon. From this, a secondary elimination of the δ-hydroxy-group takes place, although the specific details remain unknown. Work on the curacin biosynthesis has established that the canonical catalytic dyad, His-Asp, found in all DH domains can also catalyze a subsequent vinylogous dehydration consistent with our observations.^[22]^ Similarly, vinylogous amino acids are very rare in nature and usually endowed with biological activity, and we have only identified the motif predicted in the intermediate containing a vinylogous serine in the natural product, thalassospiramide, where it is trapped in a macrolactone.^[27,28]^ While we hypothesize a key role for the dehydratase, we cannot rule out the involvement of the KR domain or the influence of the unusual domain order. In a recent study on the otherwise unrelated PKS-NRPS hybrid BGC associated with the natural product, rhabdobranin, the usual KR-DH didomain arrangement was associated with an epimerization of L- to D-arginine.^[29]^

## Conclusion

Natural products provide an extensive source of structural diversity, but also new mode-of-actions. One interesting natural product family is the APD-CLDs, in which we and others have connected several members to their corresponding biosynthetic gene clusters. However, the full understanding of how the APD warhead is installed remains unknown. In this study, we have used the XUT approach in NRPS engineering to confirm that it is the core PKS machinery that is responsible for the implementation of the APD warhead. By developing a synthetic standard, we have confirmed the APD warhead-containing structure. Furthermore, we have shown that the dehydratase within the PKS module is involved in the elimination reactions required for the diene construction. By disturbing the conserved catalytically active residues, the machinery is subjected to module stuttering, and the growing NRP/PK chain is released from the assembly line. We therefore propose that the two elimination reactions involved in the APD warhead assembly are carried out sequentially, and that, upon disturbing the first, the second cannot be carried out. After multiple point mutations, we have yet to identify any amino acid residues responsible for the second elimination reaction. Whether it is the dehydratase that facilitates both elimination reactions, or another enzyme, such as the ketoreductase, that is involved, remains unknown and needs to be further investigated.

## Supporting Information

The authors have cited additional references within the Supporting Information.^[30-36]^

## Acknowledgements

The work was supported by a grant from AUFF (AUFF-E-2022-9-42) to T.T. Funding from Independent research fund Denmark (grant 2032-00219B to T.B.P.) and Novo Nordisk Foundation (grant NNF25OC0102763 to T.B.P.) is acknowledged.

