## Supporting Information for "Heterologous Reconstitution of 4-Amidopentadienoate Biosynthesis Identifies the PKS Machinery Responsible for Warhead Formation"

#### Figures

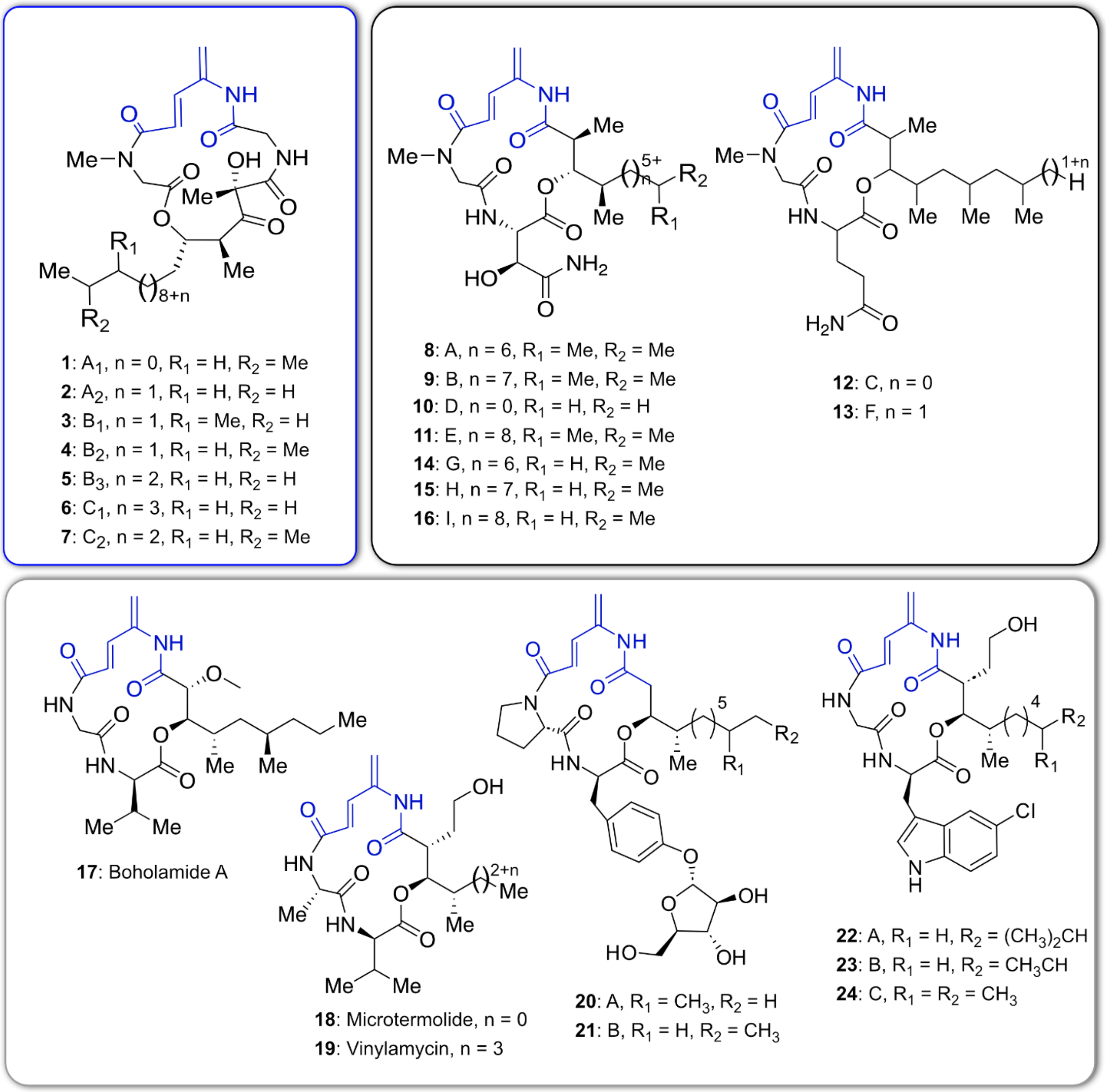


**Figure S1: All known members of the APD-CLD family.** BE-43547 (**1-7**), rakicidin (**8-16**), boholamide A (**17**), microtermolide (**18**), vinylamicin (**19**), arabimalamide (**20-21**), and chloromalamide (**22-24**). The APD moiety is highlighted in blue.

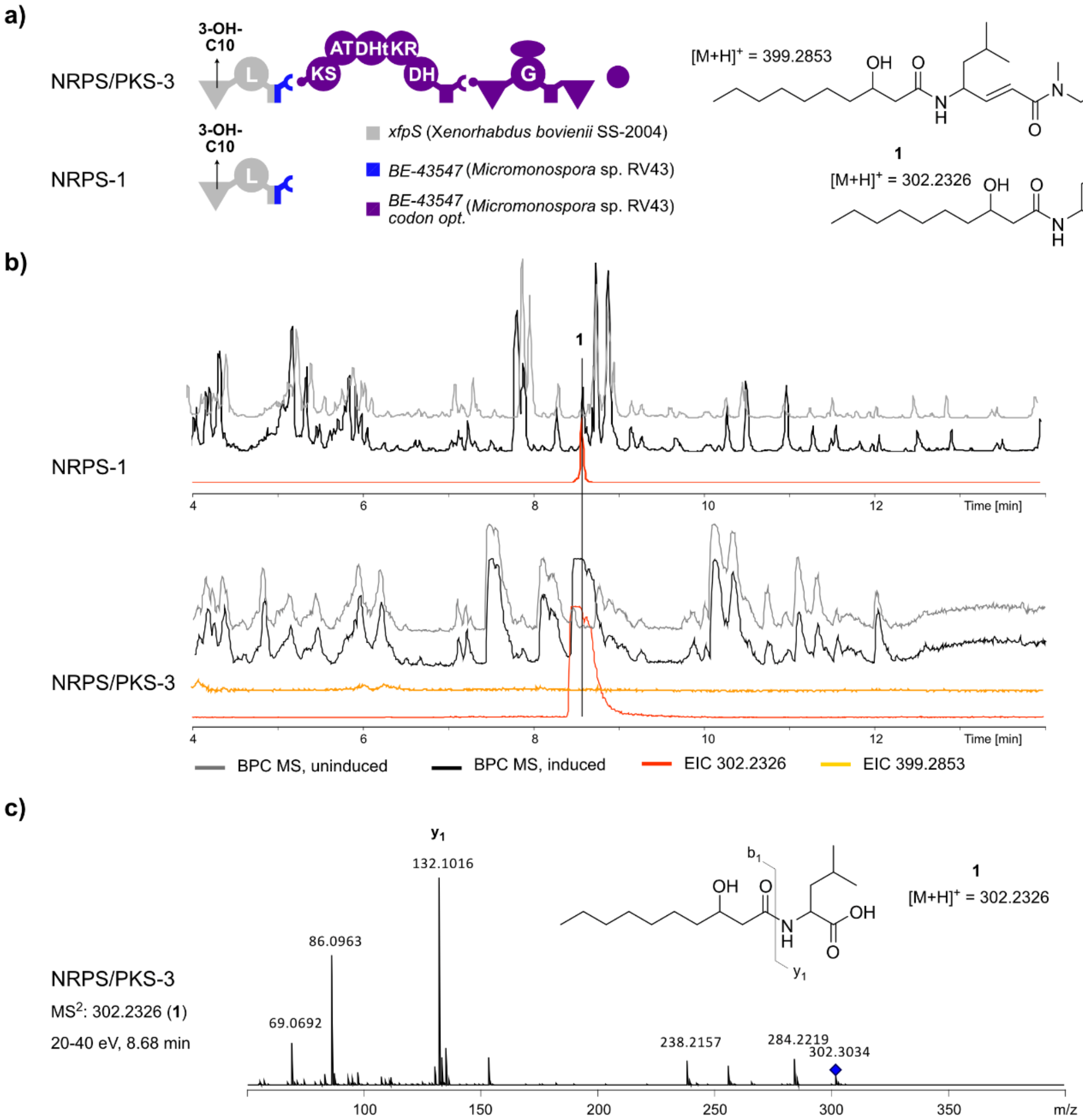


**Figure S2: Heterologous expression of NRPS/PKS-3, replacing the NRPS-4b starter unit with NRPS-1, results in production of the starter unit.** a) Description of the constructs and the expected compounds produced. b) The full NRPS/PKS-3 product is not detected, although the starter unit **1** is. The base peak chromatogram (BPC) and the extracted ion chromatograms (EIC) are shown for the two structures shown above (**1**: *m/z* 302.2306) and (*m/z* 399.2853). We have also searched for structures containing the reduced ketone or with no modification at all. All show no peaks. All productions are done in XXP3 media with 2% XAD beads. c) MS$^2$ fragmentation of **1** from NRPS/PKS-3.

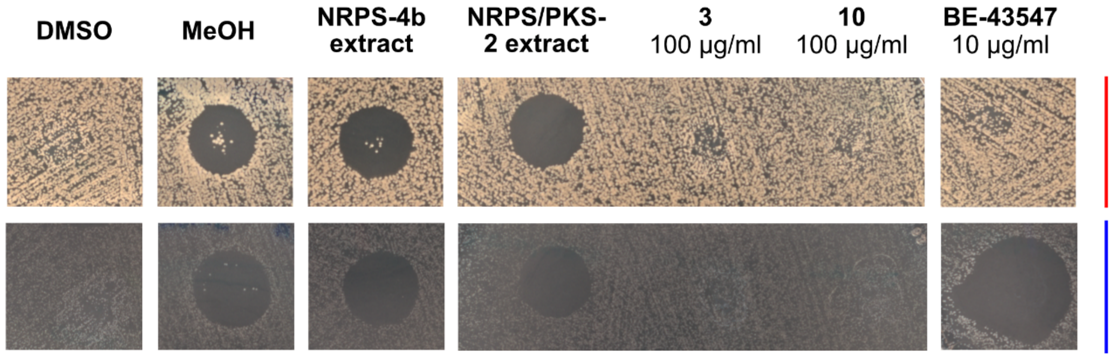


**Figure S3: Bioactivity assays on agar plates with S. aureus as the indicator strain.** No bioactivity was detected for the synthesized compounds **3** and **10**. The plates were incubated either aerobically (red) or anaerobically (blue). The tested compounds were tested at 100 µg/ml (**3** and **10**) or 10 µg/ml (BE-43547), and all were dissolved in DMSO. NRPS-4b and NRPS/PKS-2 extracts were also spotted. These were dissolved in methanol (MeOH). All tested compounds and extracts were spotted onto the plates in the same volume (10 µl).

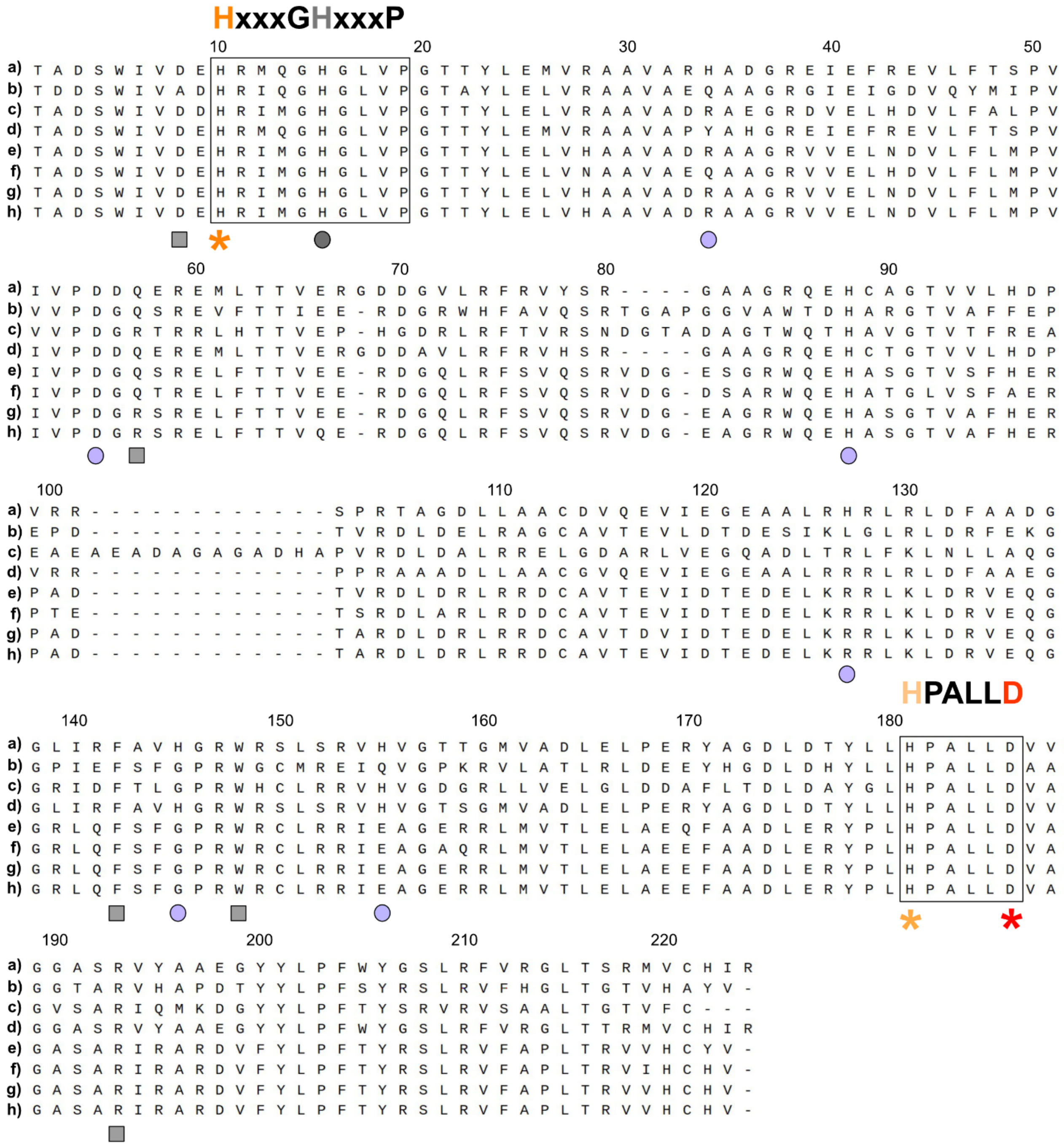


**Figure S4: Protein sequence alignment of dehydratases in APD-CLD producers, whose production has been confirmed during this study or previously in the laboratory.** *: Amino acid residues important for catalytic activity. Circle: Histidine residues and a conserved aspartic acid in *Micromonospora* sp. RV43 with the H15 indicated in grey. Square: Amino acids located near the active site. The numbering is based on the dehydratase from *Micromonospora* sp. RV43. a) *Micromonospora* sp. RV43, b) *Streptomyces* sp. MspMP-M5, c) *Streptomyces lilacinus* NRRL B-1968, d) *Salinispora arenicola* CNR 107, e) *Micromonospora purpueogenes* NRRL B-2672, f) *Micromonospora eburnea* DSM 44814, g) *Micromonospora chalcea* DSM 43026, h) *Micromonospora* sp. M42.

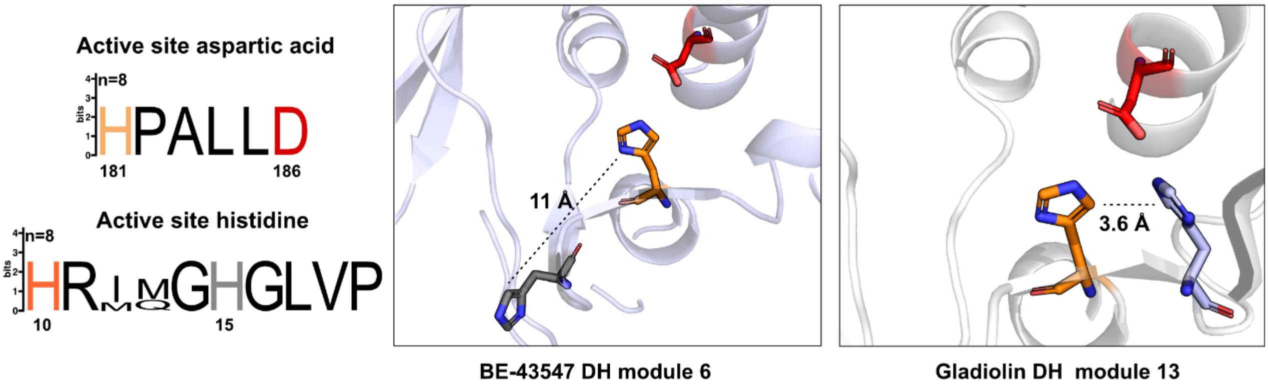


**Figure S5: Dehydratase active site motifs and structure.** The active site motif of the non-canonical DH based on all APD-CLD producers we have validated is shown in combination with the AlphaFold model of *Micromonospora* sp. RV43 dehydratase from module 6 (blue, left). The AlphaFold[1] model of the DH-like domain from module 13 of the gladiolin PKS is shown for comparison (grey, right).[2] Amino acids are colored according to what they correspond to: Active site aspartic acid (red), canonical active site histidine (orange), non-canonical active site histidine (grey for BE-43547, light-blue for gladiolin).

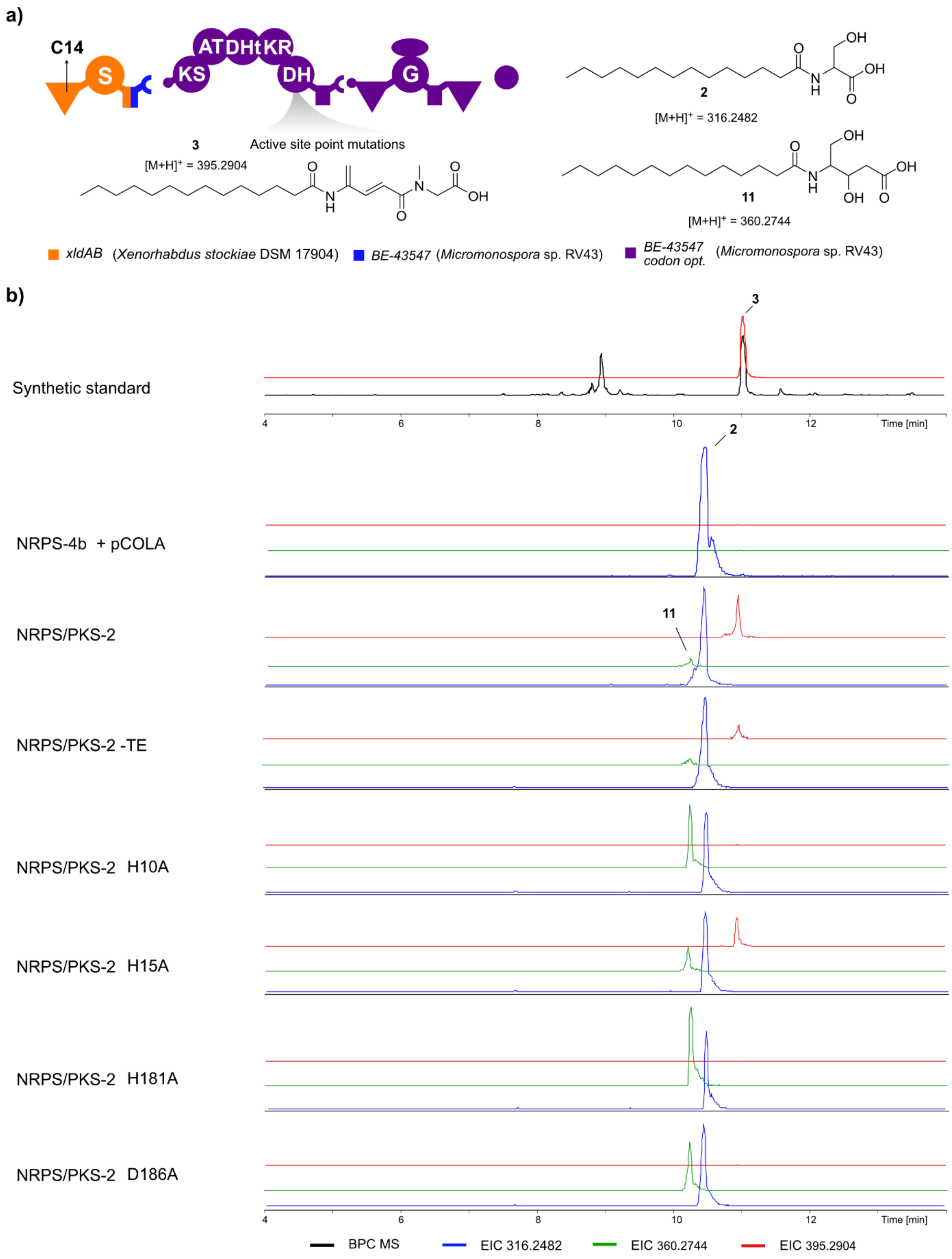


**Figure S6: Chromatograms and structures of 2, 3, and 11 derived from point mutations in the active site of the NRPS/PKS-2 DH domain.** a) Description of NRPS/PKS-2 construct with indications of point mutations in the DH active site, along with expected products **2**, **3,** and **11** indicated. b) Base peak chromatogram (BPC) is shown for the synthetic standard along with the corresponding extracted ion chromatograms (EIC). The EICs are shown for **3** (*m/z* 395.2904, red), **11** (*m/z* 360.2744, green), and **2** (*m/z* 316.2904, blue) in all productions. pCOLA is an empty vector. All productions are done in XXP3 media with 2% XAD beads. HPLC-MS data refers to **Figure 5.**

**a)**

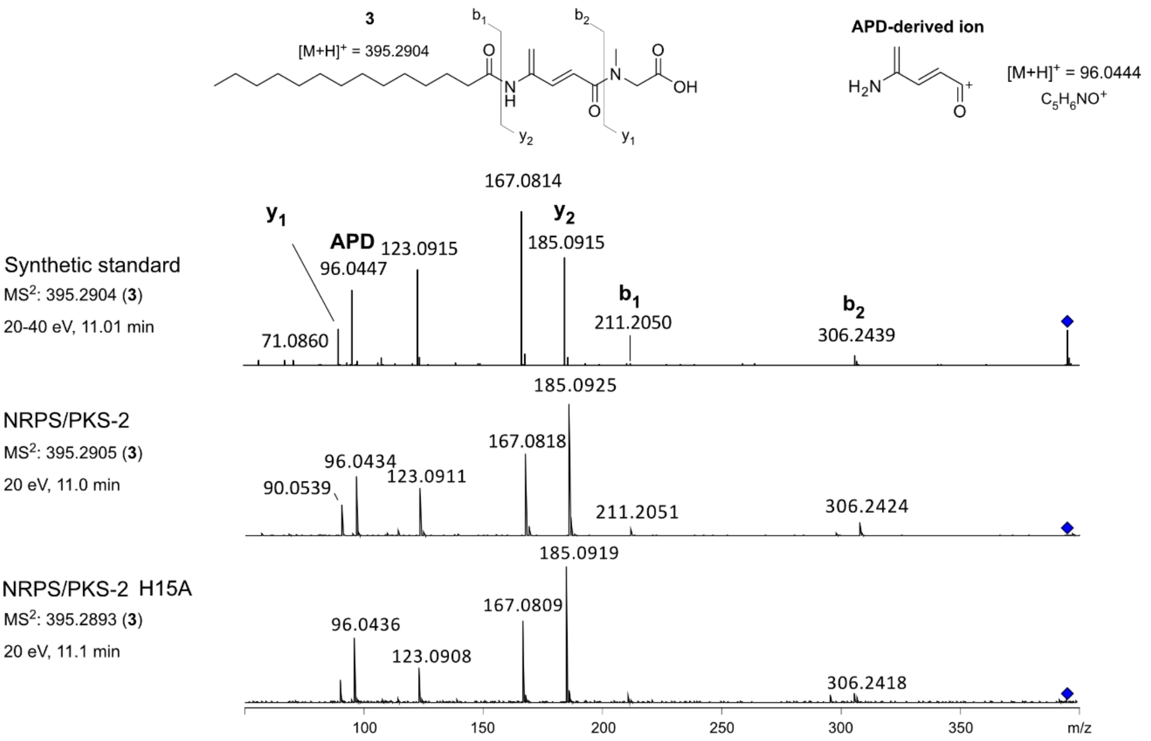
3
[M+H]+ = 395.2904
b1
b2
y1
y2
O
N
H
OH
APD-derived ion
H2N
[M+H]+ = 96.0444
C5H6NO+
Synthetic standard
MS2: 395.2904 (3)
20-40 eV, 11.01 min
y1
APD
96.0447
123.0915
167.0814
y2
185.0915
b1
211.2050
b2
306.2439
71.0860
NRPS/PKS-2
MS2: 395.2905 (3)
20 eV, 11.0 min
185.0925
96.0434
167.0818
123.0911
90.0539
211.2051
306.2424
NRPS/PKS-2 H15A
MS2: 395.2893 (3)
20 eV, 11.1 min
185.0919
96.0436
167.0809
123.0908
306.2418
100
150
200
250
300
350
m/z

**b)**

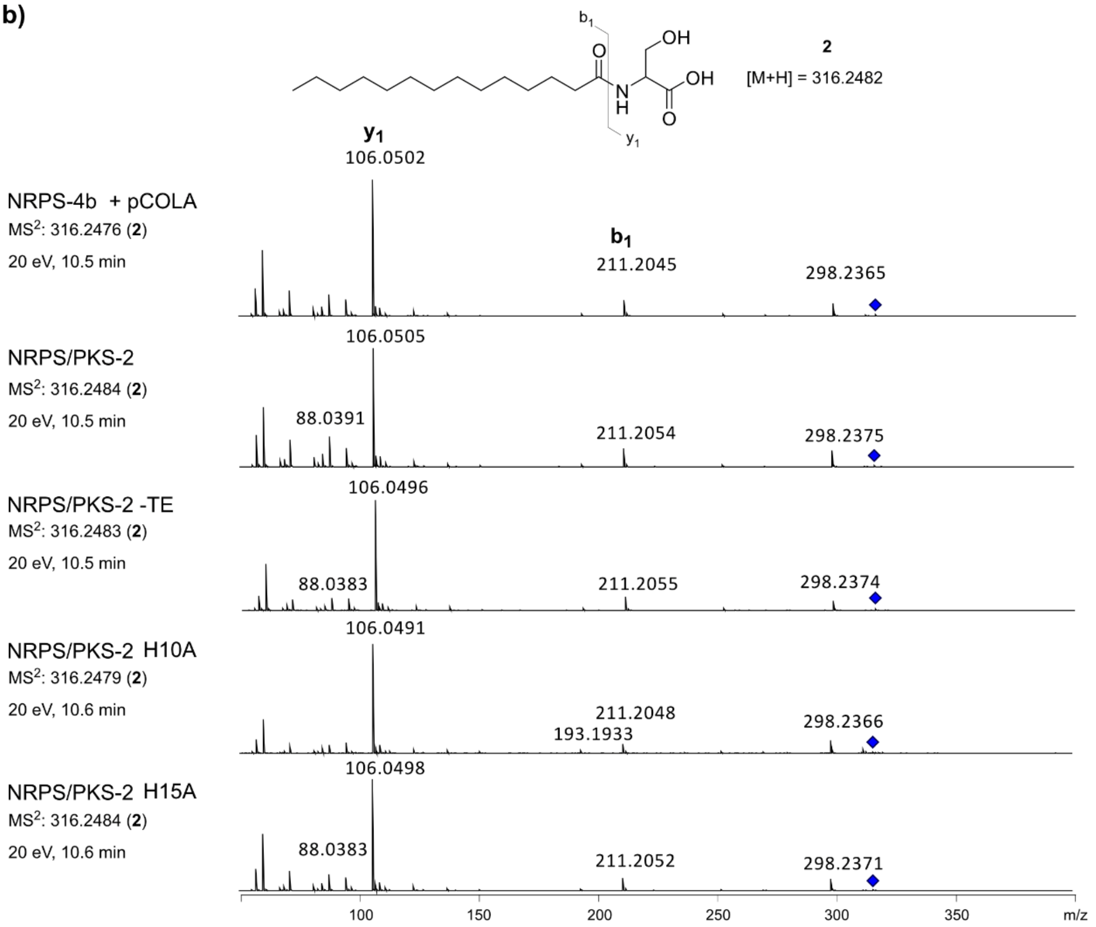
b1
y1
OH
O
N
H
OH
O
2
[M+H] = 316.2482
y1
106.0502
NRPS-4b + pCOLA
MS2: 316.2476 (2)
20 eV, 10.5 min
b1
211.2045
298.2365
NRPS/PKS-2
MS2: 316.2484 (2)
20 eV, 10.5 min
106.0505
88.0391
211.2054
298.2375
NRPS/PKS-2 -TE
MS2: 316.2483 (2)
20 eV, 10.5 min
106.0496
88.0383
211.2055
298.2374
NRPS/PKS-2 H10A
MS2: 316.2479 (2)
20 eV, 10.6 min
106.0491
211.2048
193.1933
298.2366
NRPS/PKS-2 H15A
MS2: 316.2484 (2)
20 eV, 10.6 min
106.0498
88.0383
211.2052
298.2371
100
150
200
250
300
350
m/z

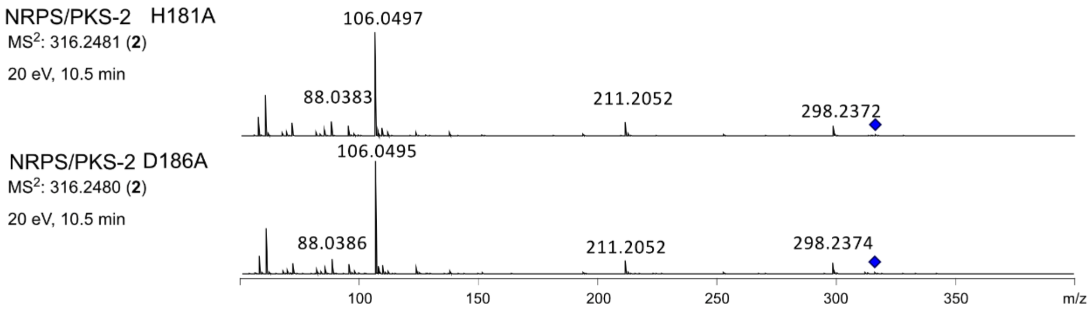


c)

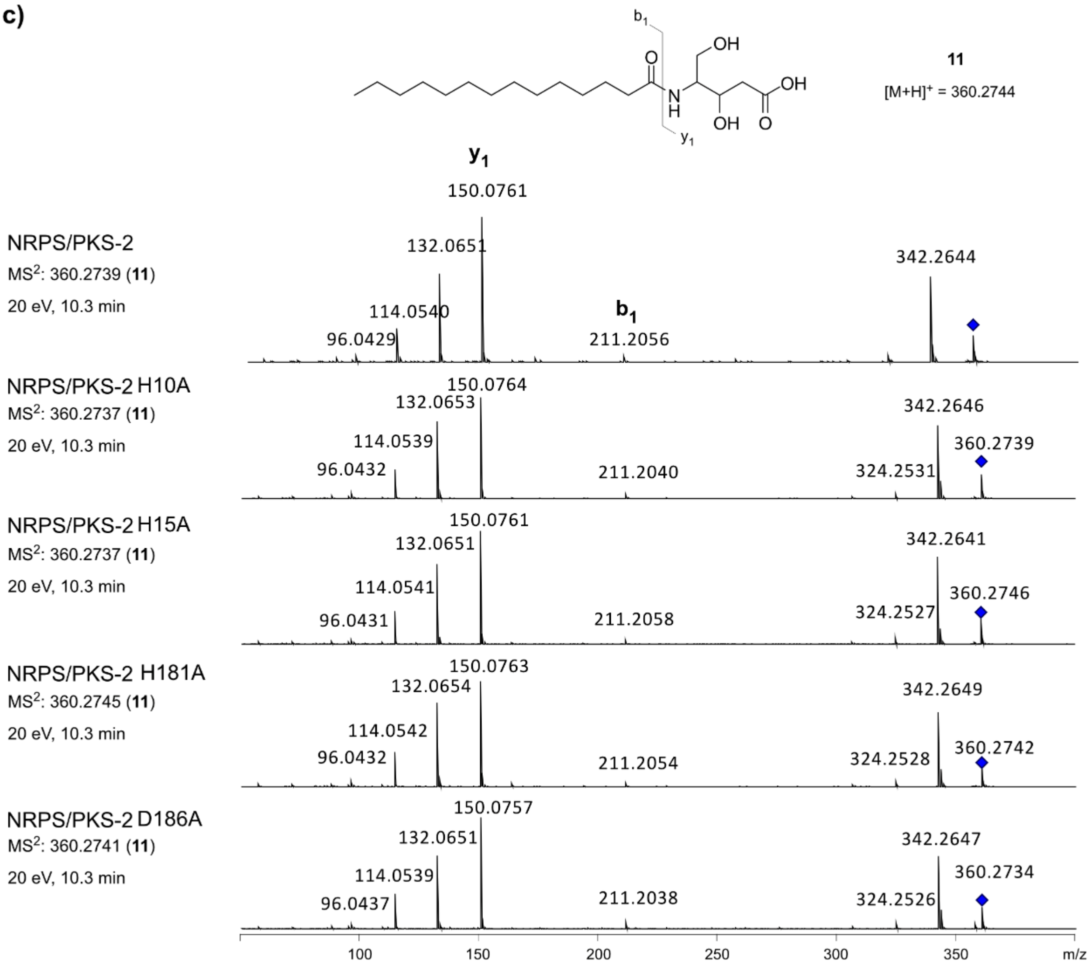


**Figure S7: MS$^2$ fragmentation spectra of 2, 3, and 11 derived from point mutations in the active site of the NRPS/PKS-2 DH domain.** a) MS$^2$ fragmentation of **3** for productions with the compound present, except for NRPS/PKS-2 -TE, as the signal was too low to be selected for fragmentation. b) MS$^2$ fragmentation of **2** for productions with the compound present. c) MS$^2$ fragmentation of **11** for productions with the compound present, except for NRPS/PKS-2 -TE, as the signal was too low to be selected for fragmentation. HPLC-MS/MS data refers to **Figure 5**.

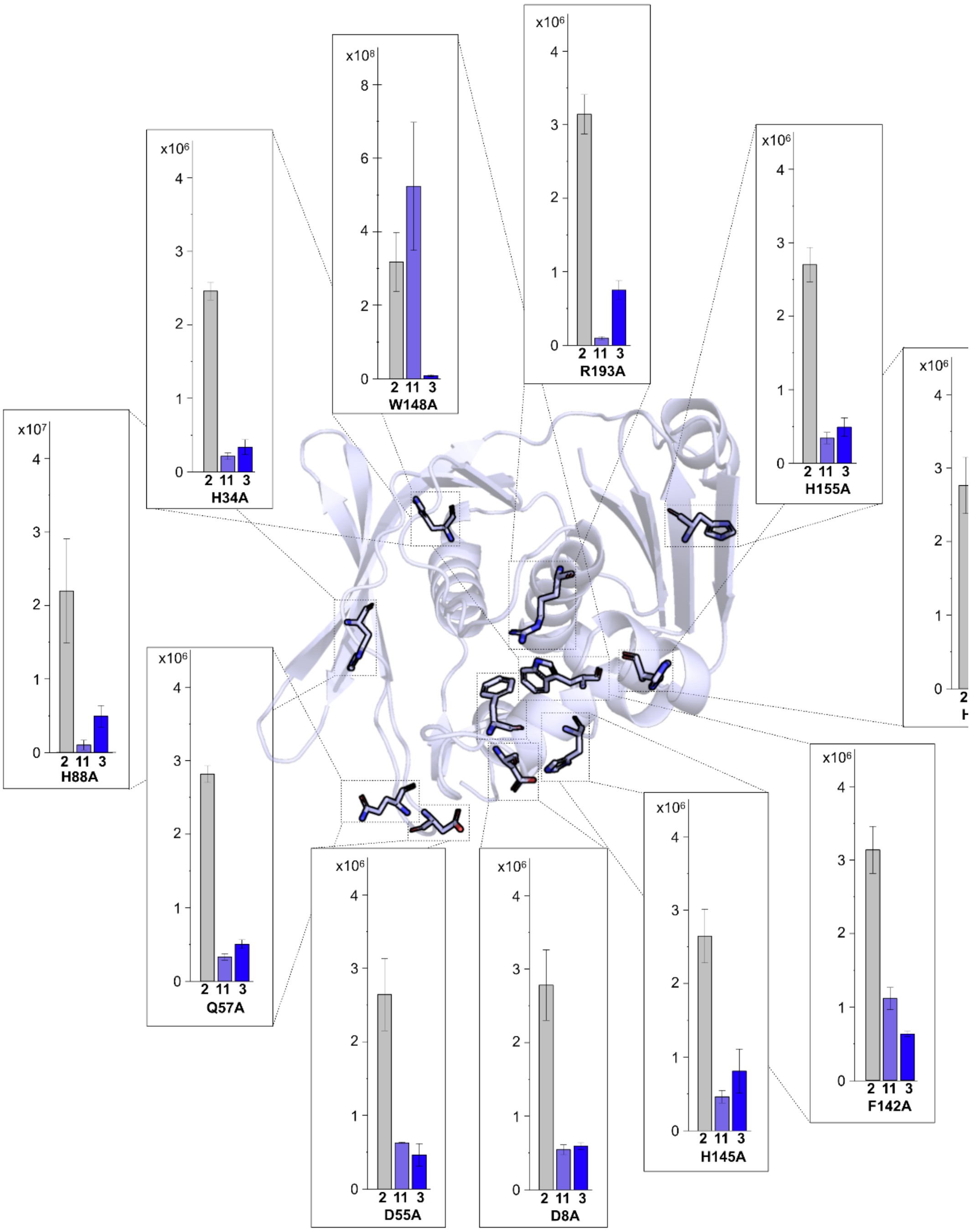


**Figure S8: Production titers of 2, 3, and 11 from point mutations within the DH domain of the NRPS/PKS-2 construct.** Production titers are based on the ion count of the MS area using HPLC-MS (n=3 ±SD). The AlphaFold[1] model of the *Micromonospora* sp. RV43 dehydratase is shown with the amino acid residues of interest indicated in sticks.

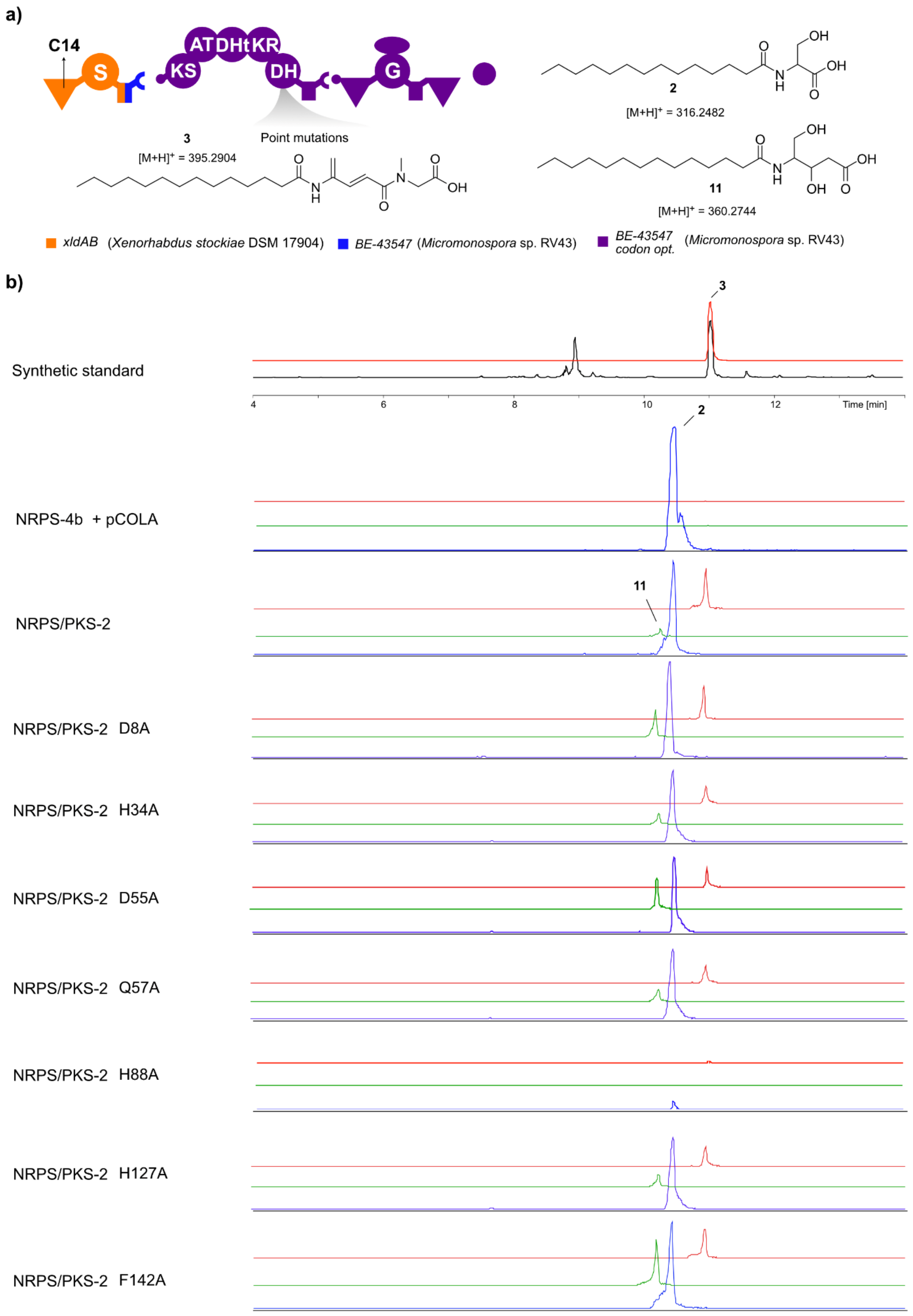

a)
C14
S
KS
AT
DH
t
KR
DH
G
Point mutations
3
[M+H]+ = 395.2904
2
[M+H]+ = 316.2482
11
[M+H]+ = 360.2744
xldAB (Xenorhabdus stockiae DSM 17904)
BE-43547 (Micromonospora sp. RV43)
BE-43547 codon opt. (Micromonospora sp. RV43)
b)
Synthetic standard
Time [min]
NRPS-4b + pCOLA
NRPS/PKS-2
NRPS/PKS-2 D8A
NRPS/PKS-2 H34A
NRPS/PKS-2 D55A
NRPS/PKS-2 Q57A
NRPS/PKS-2 H88A
NRPS/PKS-2 H127A
NRPS/PKS-2 F142A

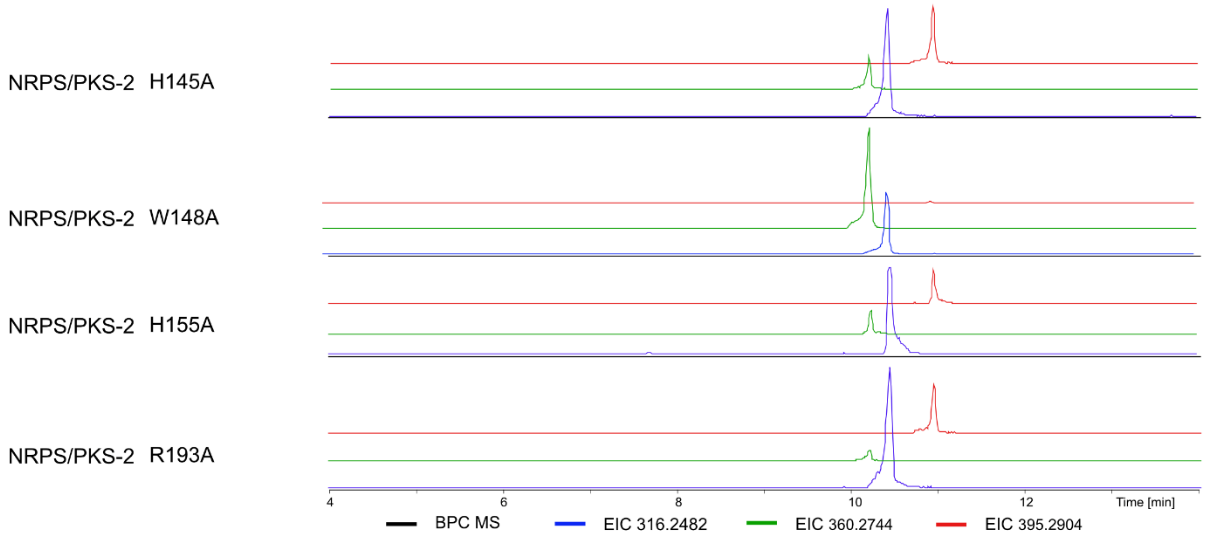


**Figure S9: Chromatograms and structures of 2, 3, and 11 derived from point mutations in the NRPS/PKS-2 DH domain.** a) Description of the NRPS/PKS-2 construct with indications of point mutations in the DH domain, along with expected products **2**, **3,** and **11** indicated. b) Base peak chromatogram (BPC) is shown for the synthetic standard along with the corresponding extracted ion chromatograms (EIC). The EICs are shown for **3** (*m/z* 395.2904, red), **11** (*m/z* 360.2744, green), and **2** (*m/z* 316.2904, blue) in all productions. pCOLA is an empty vector. All productions are done in XXP3 media with 2% XAD beads. HPLC-MS data refers to **Figure S8**.

**a)**

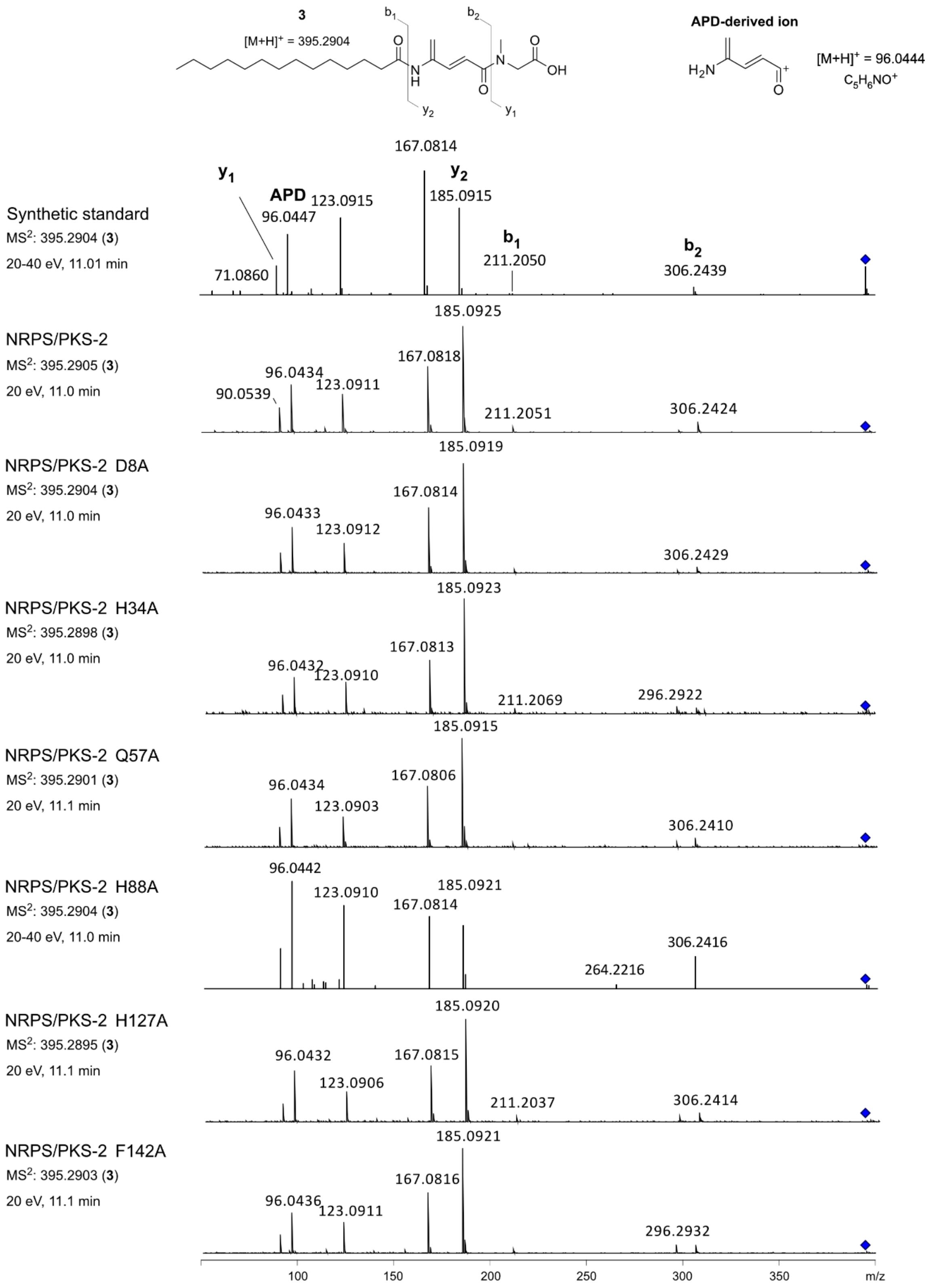

3
[M+H]+ = 395.2904
b1
b2
y2
y1
APD-derived ion
[M+H]+ = 96.0444
C5H6NO+
Synthetic standard
MS2: 395.2904 (3)
20-40 eV, 11.01 min
y1
APD
y2
b1
b2
71.0860
96.0447
123.0915
167.0814
185.0915
211.2050
306.2439
NRPS/PKS-2
MS2: 395.2905 (3)
20 eV, 11.0 min
90.0539
96.0434
123.0911
167.0818
185.0925
211.2051
306.2424
NRPS/PKS-2 D8A
MS2: 395.2904 (3)
20 eV, 11.0 min
96.0433
123.0912
167.0814
185.0919
306.2429
NRPS/PKS-2 H34A
MS2: 395.2898 (3)
20 eV, 11.0 min
96.0432
123.0910
167.0813
185.0923
211.2069
296.2922
NRPS/PKS-2 Q57A
MS2: 395.2901 (3)
20 eV, 11.1 min
96.0434
123.0903
167.0806
185.0915
306.2410
NRPS/PKS-2 H88A
MS2: 395.2904 (3)
20-40 eV, 11.0 min
96.0442
123.0910
167.0814
185.0921
264.2216
306.2416
NRPS/PKS-2 H127A
MS2: 395.2895 (3)
20 eV, 11.1 min
96.0432
123.0906
167.0815
185.0920
211.2037
306.2414
NRPS/PKS-2 F142A
MS2: 395.2903 (3)
20 eV, 11.1 min
96.0436
123.0911
167.0816
185.0921
296.2932
100
150
200
250
300
350
m/z

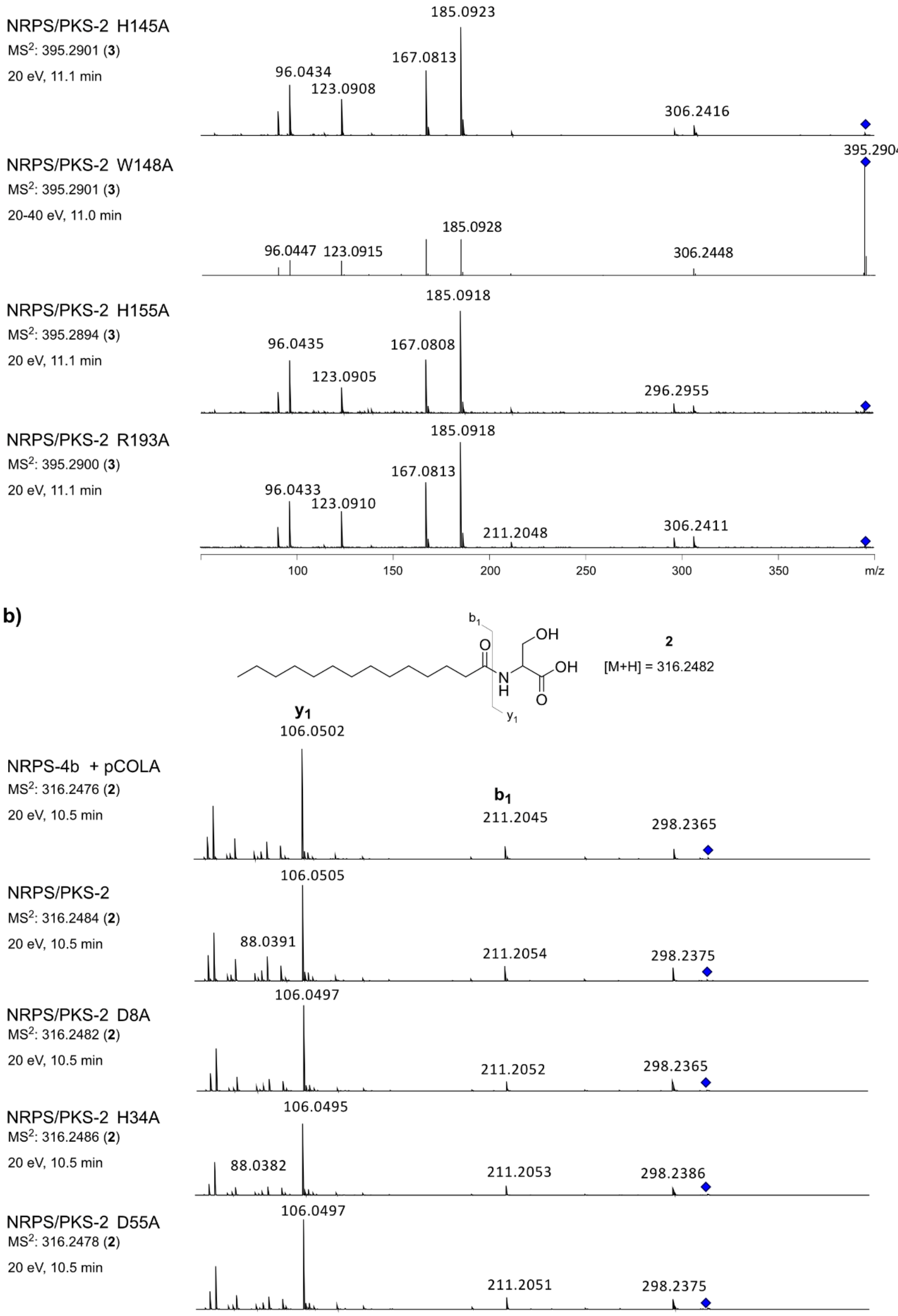

NRPS/PKS-2 H145A
MS2: 395.2901 (3)
20 eV, 11.1 min
185.0923
167.0813
96.0434
123.0908
306.2416
NRPS/PKS-2 W148A
MS2: 395.2901 (3)
20-40 eV, 11.0 min
395.2904
185.0928
96.0447
123.0915
306.2448
NRPS/PKS-2 H155A
MS2: 395.2894 (3)
20 eV, 11.1 min
185.0918
96.0435
167.0808
123.0905
296.2955
NRPS/PKS-2 R193A
MS2: 395.2900 (3)
20 eV, 11.1 min
185.0918
167.0813
96.0433
123.0910
211.2048
306.2411
100
150
200
250
300
350
m/z
b)
b1
OH
O
N
H
OH
O
y1
2
[M+H] = 316.2482
y1
106.0502
NRPS-4b + pCOLA
MS2: 316.2476 (2)
20 eV, 10.5 min
b1
211.2045
298.2365
NRPS/PKS-2
MS2: 316.2484 (2)
20 eV, 10.5 min
106.0505
88.0391
211.2054
298.2375
NRPS/PKS-2 D8A
MS2: 316.2482 (2)
20 eV, 10.5 min
106.0497
211.2052
298.2365
NRPS/PKS-2 H34A
MS2: 316.2486 (2)
20 eV, 10.5 min
106.0495
88.0382
211.2053
298.2386
NRPS/PKS-2 D55A
MS2: 316.2478 (2)
20 eV, 10.5 min
106.0497
211.2051
298.2375

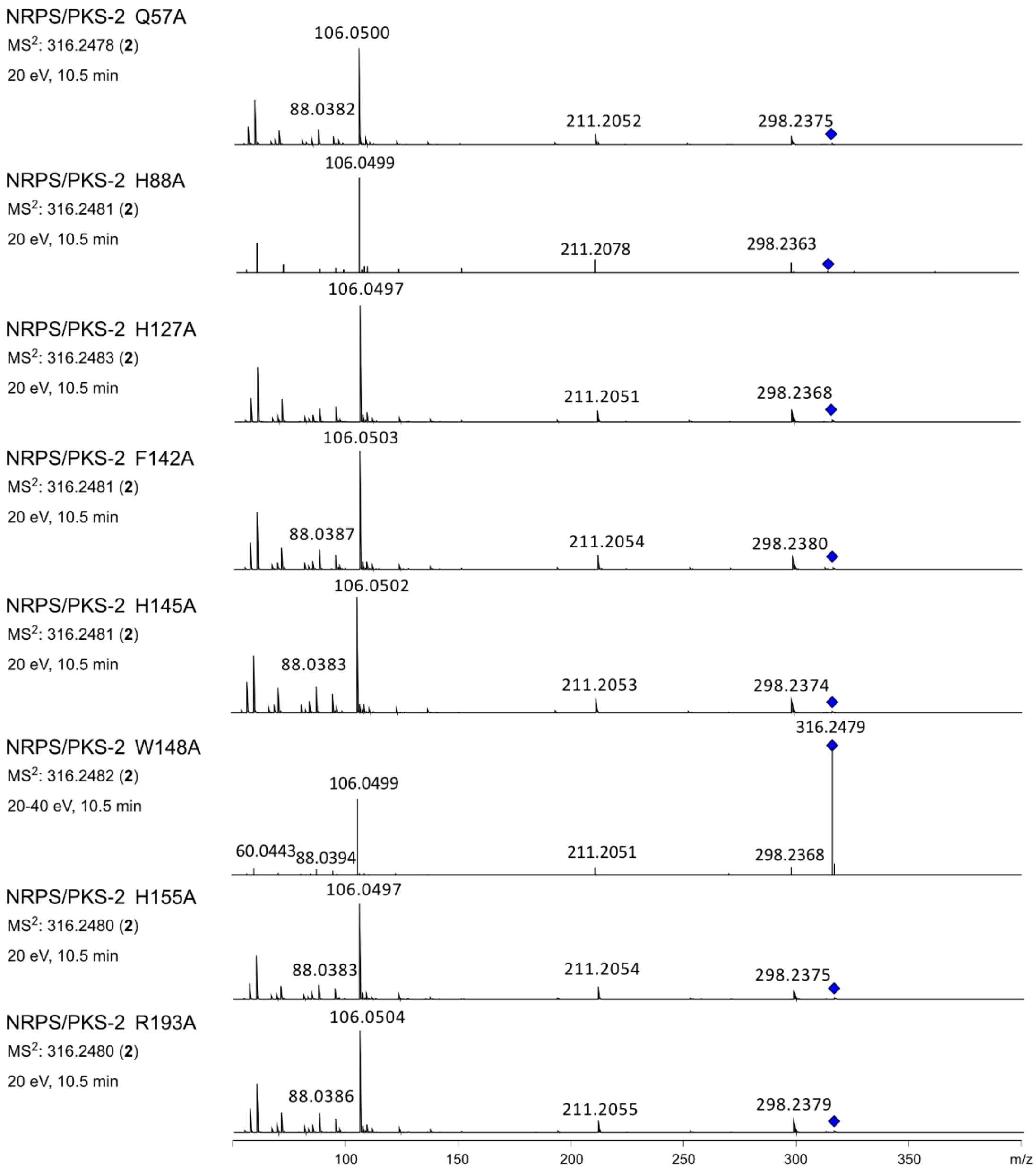
NRPS/PKS-2 Q57A
MS$^2$: 316.2478 (**2**)
20 eV, 10.5 min
106.0500
88.0382
211.2052
298.2375
NRPS/PKS-2 H88A
MS$^2$: 316.2481 (**2**)
20 eV, 10.5 min
106.0499
211.2078
298.2363
NRPS/PKS-2 H127A
MS$^2$: 316.2483 (**2**)
20 eV, 10.5 min
106.0497
211.2051
298.2368
NRPS/PKS-2 F142A
MS$^2$: 316.2481 (**2**)
20 eV, 10.5 min
106.0503
88.0387
211.2054
298.2380
NRPS/PKS-2 H145A
MS$^2$: 316.2481 (**2**)
20 eV, 10.5 min
106.0502
88.0383
211.2053
298.2374
NRPS/PKS-2 W148A
MS$^2$: 316.2482 (**2**)
20-40 eV, 10.5 min
316.2479
106.0499
60.0443
88.0394
211.2051
298.2368
NRPS/PKS-2 H155A
MS$^2$: 316.2480 (**2**)
20 eV, 10.5 min
106.0497
88.0383
211.2054
298.2375
NRPS/PKS-2 R193A
MS$^2$: 316.2480 (**2**)
20 eV, 10.5 min
106.0504
88.0386
211.2055
298.2379
100
150
200
250
300
350
m/z

**c)**

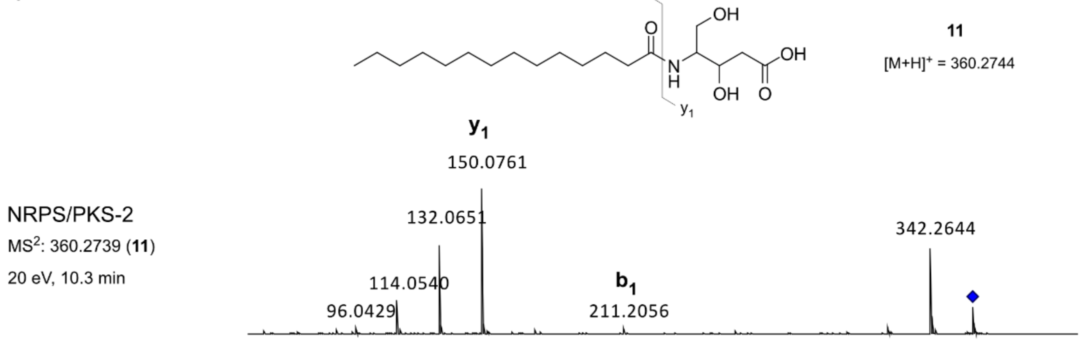
b1
y1
OH
O
N
H
OH
OH
O
11
[M+H]+ = 360.2744
y1
150.0761
NRPS/PKS-2
MS$^2$: 360.2739 (**11**)
20 eV, 10.3 min
132.0651
342.2644
114.0540
b1
96.0429
211.2056

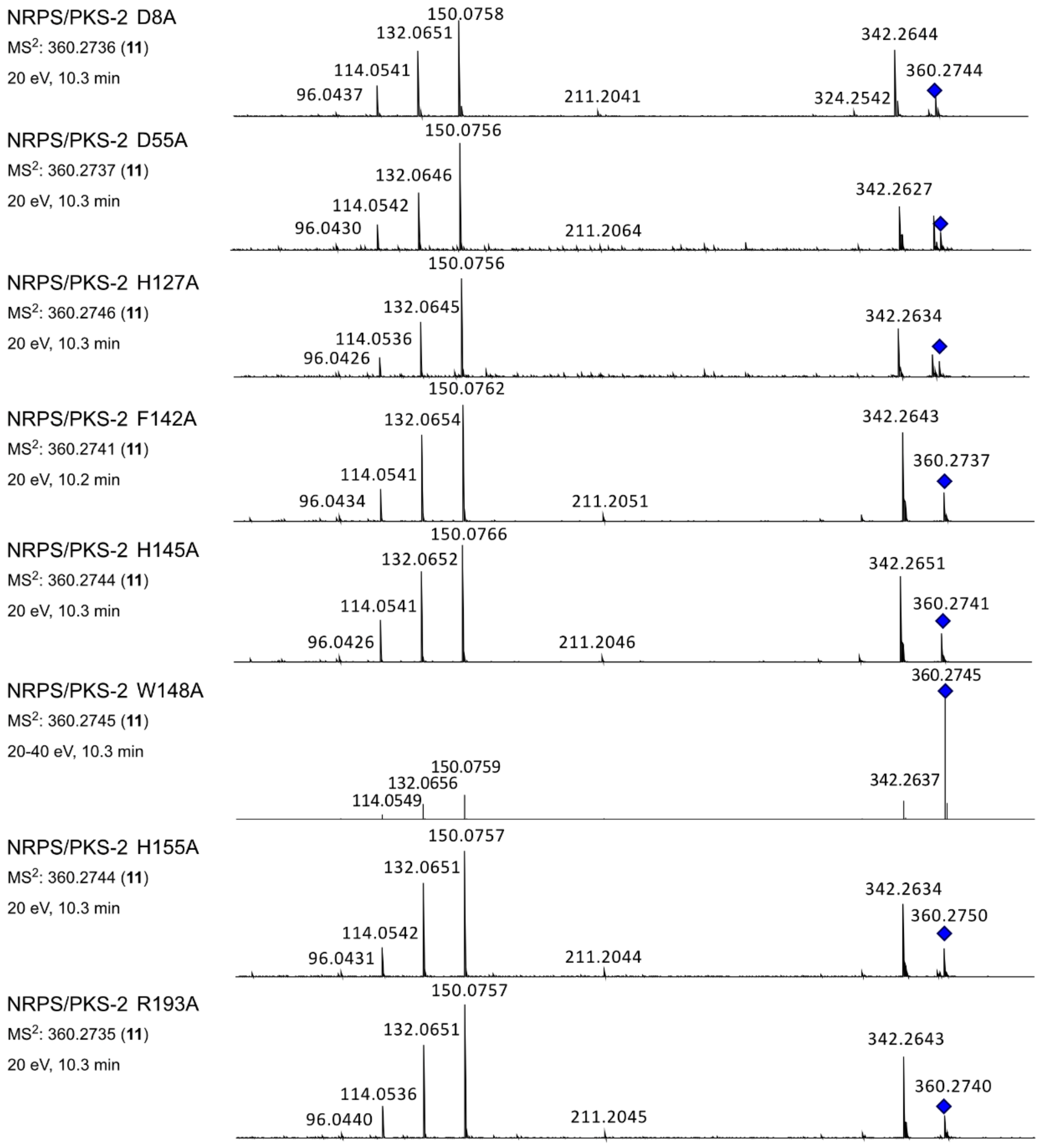


**Figure S10: MS² fragmentation spectra of 2, 3, and 11 derived from point mutations in the NRPS/PKS-2 DH domain.** a) MS² fragmentation of **3** from productions, except for D55A, as the signal was too low to be selected for fragmentation. b) MS² fragmentation of **2** from productions. c) MS² fragmentation of **11** from productions, except for H34A and Q57A, as the signals were too low to be selected for fragmentation. HPLC-MS/MS data refers to **Figure S8**.

```
1                                                                                                48
a) L E G L G A I W A A G A T L D W T G L C E A D R R R R V R L P S Y P F Q R R R Y W V E P E - R P E -
b) L A G L G A V W S A G A A V D W A A V Q S P E R H R P V R L P G Y A F Q R R R Y W V E P A G L P R -
c) - - - - - - - - - - - - - - - - - - - F A D E D R R L V R L P G Y A F E R Q R Y W I D P D S T A T A
d) - - G L G T I W A A G A T L D W S A V C E A D R R R R V R L P S Y P F R R R R Y W V E P D - R P E -
e) L A G L G D L W A A G A P V D W A A R T G G T - R R L L R L P T Y P F Q R Q R Y W I E P Q P G P V -
f) L A G L G E L W S A G V P V D W T A R D G D A T R R L M R L P T Y A F Q R Q R Y W I D P E T T P - -
g) L A G L G E L W A A G A P V D W A A R A G D T P R R L L R L P T Y P F Q R Q R Y W I E P R P G P V -
```

```
49                                                                                               87
a) - T S G P A S P P A - - - S S S T Y R P S W R R L Q - P - V R P R A E A - - - - - G P W L V L G A D
b) - A A E P A A T T A - - - D G W F Y T P G W Q R L A L P - V R S T A A A T G - - - D T W V I L G A G
c) D E P E P V L S A A A R L D D W F H V P T W R R L P A D G G H P R P D E D - - - - T V W A V L G T D
d) - T S G P A T P A G - - - S S P T Y R P S W R R L Q - P - G R P G V D D - - - - - G P W L L L G A E
e) - R P A P V T A T G P D - A Q W L H V P G W R R L T A P - A A G T T D A - - - - - A T W V V L G A E
f) - A P R R A V T T G A D - A E W F H V P G W R R L T T A - V A E A A G G D S G G V A V W V L L G A E
g) - R P A P V A A T G P D - A Q W F H V P G W R R L T A P - A A G T T D G - - - - - A T W V V L G A E
```

```
88                                                                                              133
a) L P E G R A L A E S L P S - - - - V V R V H A G A G F A Q G G P D A F T V D P A D R D D Y R R L L D
b) R A L G D A L A D R L A A A G D L V V R V T A G T S M E R T G E G S W T V D P A D R D H L S A L A S
c) L A L G G A L A R R L E D D G A T V V R V A A G D A L D R A G D R S W T L S P T S R E H L T D L L K
d) L P A G R A L A G S L P A - - - - V V R V R A G S G F A Q D G P D A F T V D P T D R D D H R R L L D
e) L P L G V A L A G R L A G D G A T V L R V S A G D E L R E T G E R A W S L D P A S R D Q H A A L L K
f) L P L G A E L A A R L A G G D A V V L R V S A G D E L R E T G D R A W E L D P T S R D Q Y A G L L K
g) L P L G V A L A G R L A D D G A T V L R V S A G D E L R E T G E R A W S L D P A S R D Q H A A L L K
```

```
                                       HxxxxxxxP
134                                                                                             169
a) A L - - - - - - - G D R P P A R V V H L W S L A V E P G - - - - - - - T D L D R A E A V G F H S L L
b) S L A T P P F A V R D G Q R F R F V H L W G A A E G P V D G A A P T H E R L D A A R R T G F D S L L
c) S L - - - - - D A D G A K A L R V V H L W S T A V A P A A R L D A - - A A L D A A R R A G F D S L L
d) A L - - - - - - - G D R R P T R V V H L W S L A V E P G - - - - - - - T D L D R A E A L G F H S V L
e) S L - - - - - E V D A G R T I R V V H L W S L T V E P G D G P D P - - D R I D S A R R V G F D S L L
f) S L - - - - - E T Y A P G T I R V V H L W S L T A E P A D G L D G - - G R I D R A R R V G F D S L L
g) S L - - - - - E V D A G R T I R V V H L W S L T V E P G D G P D P - - D R I D S A R R V G F D S L L
```

```
170                                                                                             219
a) A L A Q A L A D R A E H P A L T I E V L T R G V F S V T G D E C L Q P E N A T V I G P C T V V P Q E
b) A L A Q A L G A A R L P A P V V V D L L C R G V Y D V T G D E P L Q P E H A L L L G A A T V I P Q E
c) A L A Q A A G D V R P T A P L T V D V L G R G L Y D V T G E E A L Q P E N A G L L G A A T V V P Q E
d) A F A Q A L A D R P D Q L A L T L D V L T R G V F G V T G D E R L Q P E N A T L V G P C T V V P Q E
e) A L A Q A I G D V Q P A G P V E I D V L G R G L F A V T G D E D L Q P E N A P L L G A T T V I P Q E
f) A L A Q A I G D V Q P T G P V E L D V L S R G L F G V T G D E E L Q P E N A - - - - - - - - - - - - -
g) A L A Q A I G D V Q P A G P V E I D V L S R G L F A V T G E E S L Q P E N A P L L G A T T V I P Q E
```

```
220                                                                                             255
a) V P G V T C R L L D V T G T G - - - - - - - - - - - - - - V E S L A D A V R S A P D E P V L A R R G
b) T A E T A C R V L D I T G T D P D A P G Q D A - - - - - - V R A V L T A L T D T D A G A E L A L R G
c) V E G T L C R V L D V T G A D P H H P G E E T - - - - - - V D D L V A V L R R P V D D R E L A L R G
d) V P G V T C R L L D L T G T G - - - - - - - - - - - - - - V E S L A D A L R S A S D E P V L A R R G
e) V P D V T C R V L D L T G A D L S A A D P A P G R P A G V P T T V L A R L T A P A A D R E L A L R G
f) - - - - - - - - - - - - - - - - - - - - - - - - - - - - - - - - - - - - - - - - - - - - - - - - - -
g) V P D V T C R V L D L P G A D L P A A D P A P G - - - - - P T T V L A R L T A P T A D R E L A L R G
```

```
256                                            279
a) S H W W A R S F D A V D L D P - G P A T G R L R D
b) R H L W A R G F D R V D W I D - R T A P G D G D R
c) R H W W V R D F A A V A L - - - - - - - - - - - -
d) S H W W T R T F D A V D - - - - - - - - - - - - -
e) R H W W V R D F E A L P A D A T G A T G G R L R P
f) - - - - - - - - - - - - - - - - - - - - - - - - -
g) R H W W V R D F E A L P A D A A G G T G G R L R P
```

**Figure S11: Protein sequence alignment of DHt domains in APD-CLD producers, whose production has been confirmed during this study or previously in the laboratory.** A possible catalytic active motif is highlighted (HxxxxxxxP), although it is one amino acid residue shorter than canonical DH domains. Histidine residues in *Micromonospora* sp. RV43 DHt sequence changed into alanine residues are indicated by circles. a) *Micromonospora* sp. RV43, b) *Streptomyces* sp. MspMP-M5, c) *Streptomyces lilacinus* NRRL B-1968, d) *Salinispora arenicola* CNR 107, e) *Micromonospora purpueogenes* NRRL B-2672, f) *Micromonospora eburnea* DSM 44814, g) *Micromonospora chalcea* DSM 43026, h) *Micromonospora* sp. M42.

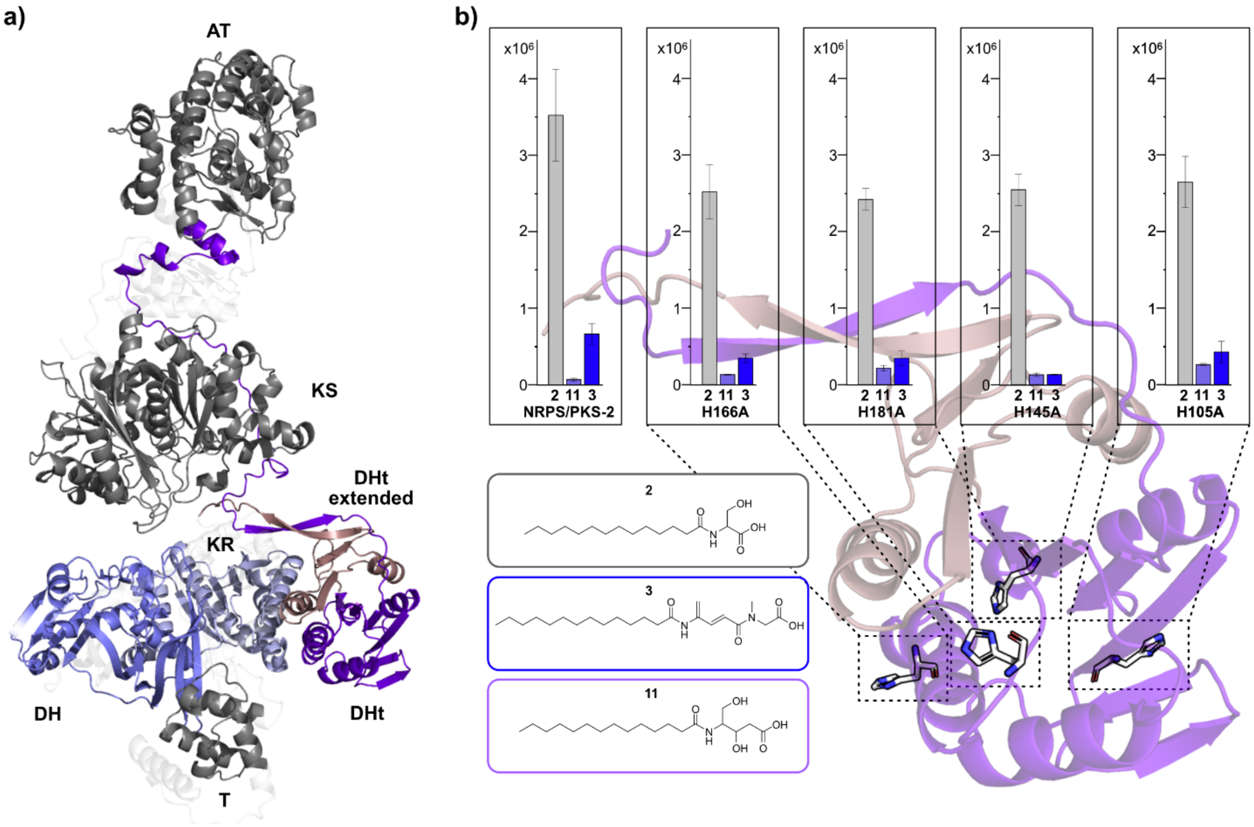


**Figure S12: AlphaFold model of PKS module 6 from Micromonospora sp. RV43 and the DHt point mutations**. a) AlphaFold[1] model of PKS module 6 from *Micromonospora* sp. RV43, with the different domains indicated. DH: blue, KR: light blue, DHt: purple, DHt extended N-terminal: light purple, KS, AT, and T domains: dark grey, connecting regions: light grey. b) AlphaFold model of the DHt domain and extended N-terminal region with amino acids of interest indicated in sticks. Production titers of **2**, **3**, and **11** are based on the ion count of the MS area using HPLC-MS **(**n=3 ±SD**).**

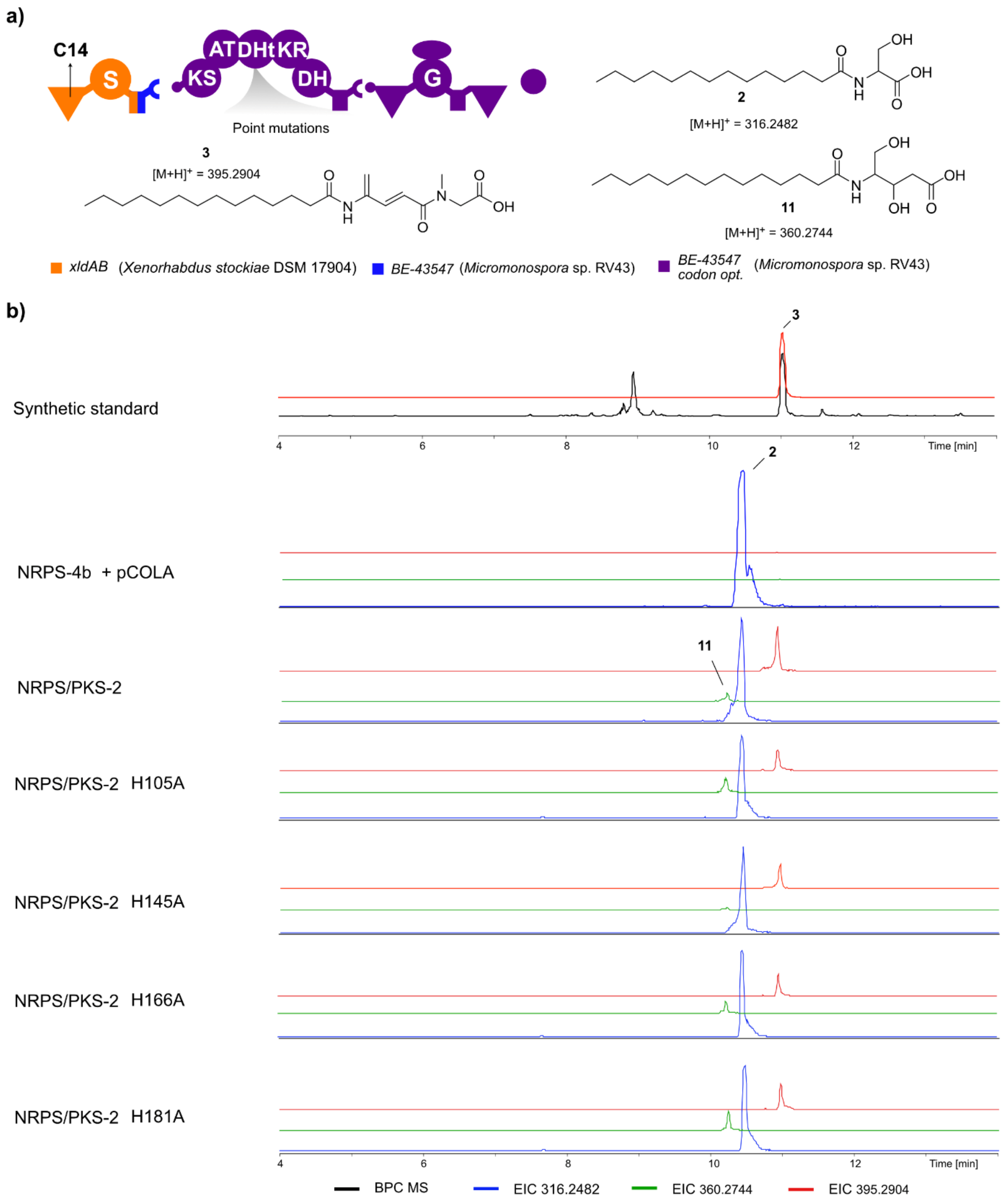


**Figure S13: Chromatograms and structures of 2, 3, and 11 derived from point mutations in the NRPS/PKS-2 DHt domain.** a) Description of NRPS/PKS-2 construct with indications of point mutations in the DHt domain, along with expected products **2**, **3,** and **11** indicated. b) Base peak chromatogram (BPC) is shown for the synthetic standards along with the corresponding extracted ion chromatograms (EIC). The EICs are shown for **3** (*m/z* 395.2904, red), **11** (*m/z* 360.2744, green), and **2** (*m/z* 316.2904, blue) in all productions. pCOLA is an empty vector. All productions are done in XXP3 media with 2% XAD beads. HPLC-MS data refers to **Figure S12**.

**a)**

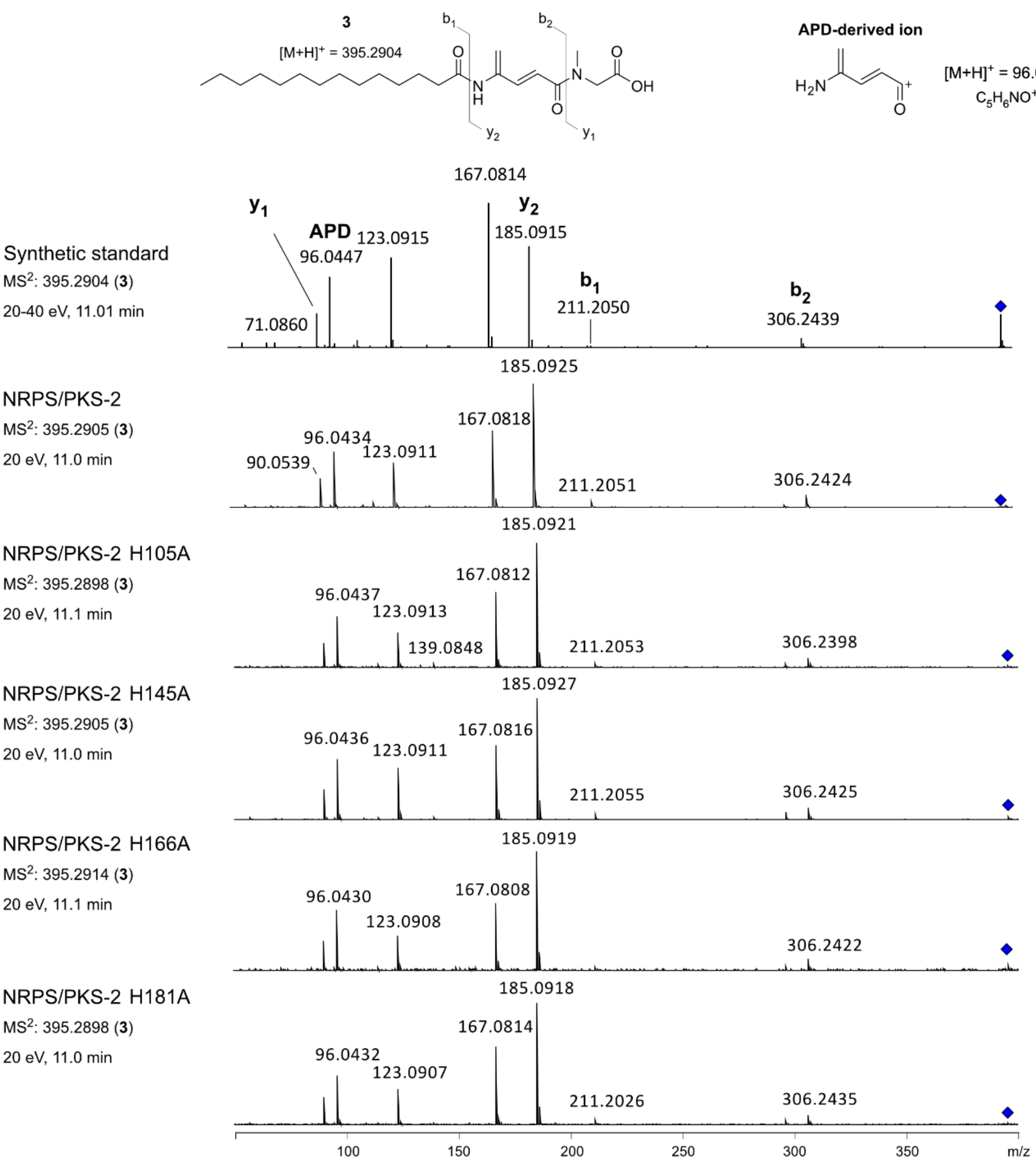

3
[M+H]+ = 395.2904
b1
b2
y2
y1
O
N
H
O
N
O
OH
APD-derived ion
H2N
O
[M+H]+ = 96.0444
C5H6NO+
Synthetic standard
MS2: 395.2904 (3)
20-40 eV, 11.01 min
y1
APD
96.0447
123.0915
167.0814
y2
185.0915
b1
211.2050
b2
306.2439
71.0860
NRPS/PKS-2
MS2: 395.2905 (3)
20 eV, 11.0 min
185.0925
167.0818
96.0434
123.0911
90.0539
211.2051
306.2424
NRPS/PKS-2 H105A
MS2: 395.2898 (3)
20 eV, 11.1 min
185.0921
167.0812
96.0437
123.0913
139.0848
211.2053
306.2398
NRPS/PKS-2 H145A
MS2: 395.2905 (3)
20 eV, 11.0 min
185.0927
167.0816
96.0436
123.0911
211.2055
306.2425
NRPS/PKS-2 H166A
MS2: 395.2914 (3)
20 eV, 11.1 min
185.0919
167.0808
96.0430
123.0908
306.2422
NRPS/PKS-2 H181A
MS2: 395.2898 (3)
20 eV, 11.0 min
185.0918
167.0814
96.0432
123.0907
211.2026
306.2435
100
150
200
250
300
350
m/z


**b)**

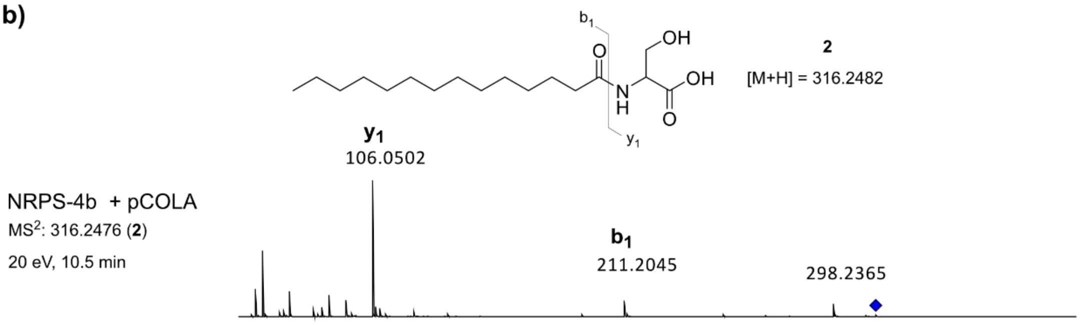

b1
OH
O
N
H
OH
O
y1
2
[M+H] = 316.2482
y1
106.0502
NRPS-4b + pCOLA
MS2: 316.2476 (2)
20 eV, 10.5 min
b1
211.2045
298.2365

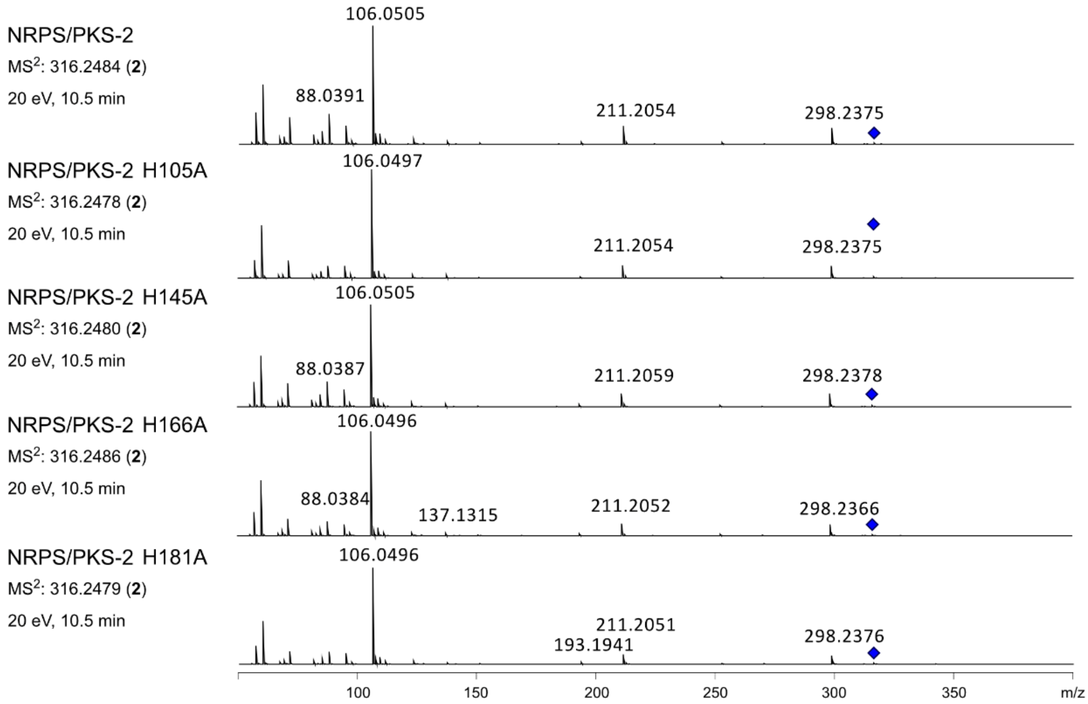


**c)**

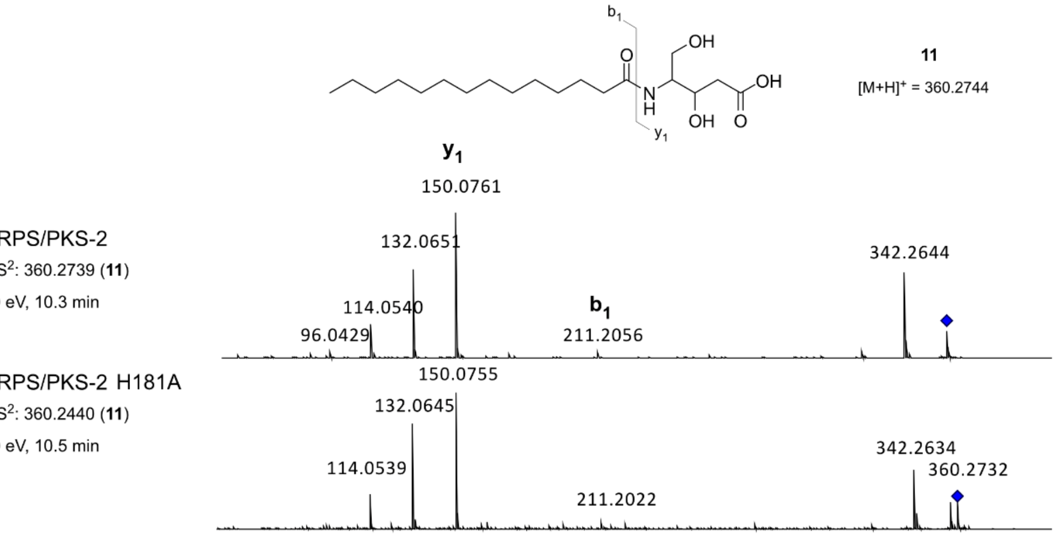


**Figure S14: MS² fragmentation spectra of 2, 3, and 11 derived from point mutations in the NRPS/PKS-2 DHt domain.** a) MS² fragmentation of **3** from productions. b) MS² fragmentation of **2** from productions. c) MS² fragmentation of **11** from productions, except for H105A, H145A, and H166A, as the signals were too low to be selected for fragmentation. HPLC-MS/MS data refers to **Figure S12**.

**Methods**

**NRPS engineering of the BE-43547 producer**

Plasmid construction

All plasmids constructed are found in **Table S2** (for NRPS engineering) and **Table S3** (for point mutations).

Linear DNA was amplified using Phusion High-fidelity DNA polymerase (NEB #M0530) and combined using NEBuilder HiFi DNA assembly (NEB #2621S). Primers and templates used for each plasmid are shown in **Table S4**. For a 50 µL reaction, the following general protocol was used unless stated otherwise, where GC buffer was added if using a high-GC template: 5x HF buffer (10 µL), 10 mM dNTPs (1 µL), Primer F (2.5 µL), Primer R (2.5 µL), DMSO (3 µL), Phusion DNA polymerase (0.5 µL), Template DNA (variable), GC buffer if needed (10 µL), nuclease-free water (to 50 µL). PCR products were loaded onto agarose gel (1% wt/vol, 30 min, 100-120 V) and gel-purified according to the manufacturer (NEB #T1120). The purified products were assembled with HiFi DNA assembly mix, according to the manufacturer, and transformed into chemically competent *E. coli* DH10B::*mtaA* or *E. coli* DH5-α as described below (incubated on selective LB agar (kanamycin: 50 µg/ml and/or chloramphenicol: 34 µg/ml), 37°C, 1 d). Overnight cultures (selective LB at the previous concentrations, 37°C, 16-18 h, 140 rpm) were used for plasmid isolation (GenJET plasmid miniprep #K0502) and verified first on agarose gel (1% wt/vol, 30 min, 100-120 V) before sequencing. Sequenced-confirmed plasmids were transformed into *E. coli* DH10B::*mtaA*, which, if needed, already contained a secondary plasmid, creating the dual-plasmid system (incubated in selective LB, 37°C, 16-18 h, 140 rpm). Glycerol stocks were prepared using ratio 1:1 of glycerol (50% wt/vol) and culture before stored at -80°C.

Transformation

Chemically competent *E. coli* DH10B::*mtaA*[6] or *E. coli* DH5-α cells were thawed on ice, mixed with 0.5-2.0 µl plasmid or HiFi assembled DNA mix. The cells were incubated on ice

(15-30 min) and subsequently heat-shocked (45 s, 42 °C). For recovery, cells were cooled down on ice for 1 min, 900 ml SOC medium was added, and the cells were incubated (1-2 h, 37 °C, 140 rpm). After recovery, cells were plated on selective LB-agar plates (kanamycin: 50 µg/ml and/or chloramphenicol: 34 µg/ml).

Heterologous expression

Overnight cultures (5 ml in 50 ml Falcon tubes) of *E. coli* DH10B::*mtaA* containing the appropriate plasmid(s) were incubated (selective LB at the previous concentrations, 37°C, 16-18 h, 140 rpm) and used as inoculum for production. Heterologous expressions were done in 24-deep well plates, where media (4 ml in XPP3), XAD 16-N adsorber resins (100 µl), L-arabinose (0.2%), inoculum (40 µl), and antibiotics (kanamycin: 25 µg/ml and/or chloramphenicol: 17 µg/ml), were mixed and the plates were incubated (22°C, 3 d, 200 rpm). Cultivation broth was removed before adding MeOH (4 ml) for extraction from the XAD beads (22°C, 1-2 h, 200 rpm). All liquid was evaporated (SpeedVac, 1-2 h, 0.1 bar), and the sample was resuspended in MeOH (200 µl). All samples were analyzed via HPLC-MS, either with timsTOF or QTOF. For QTOF, the LC-MS system consisted of UPLC-QTOF with an ACQUITY I-Class UPLC (Waters Cooperation, Milford, MA, USA) equipped with an ACQUITY UPLC HSS T3 column (2.1 mm x 100 mm, 1.8 µm, Waters) and coupled to a Bruker maXis Impact QTOF mass spectrometer (Bruker Daltonics, Bremen, Germany). All samples were analyzed with ESI+. The mobile phase consisted of milli-Q water with 0.1% formic acid (A) and a mixture of MeOH/ACN (1:1 v/v) with 0.1% formic acid (B). The flow rate was set to 0.4 mL/min at a column temperature of 50 °C with the following 21 min gradient: 0% B (0-2 min), 0-40% B (2-6 min), 40-60% B (6-6.5 min), 60-88% B (6.5-11 min), 88-100% (11-11.5min), 100% B (11.5-18 min), 100-0% B (18-19.5 min), 0% B (19.5-21 min). Collision energies were 20 eV and 30 eV. For timsTOF, the same UPLC was used but instead coupled to mass spectrometric detection on a Bruker timsMetabo (Bruker Daltonics, Bremen, Germany) operated in positive VIP-HESI (vacuum insulated probe heated ESI) mode with tims enabled. The acquisition was performed in PASEF scan mode with 2 PASEF ramps, a ramp time of 100 ms, and an accumulation time of 30 ms. The scan range was *m/z* 20-1400. Collision energies were 20 eV (CE1) and 40 eV (CE2). The capillary voltage in the VIP-HESI source was set to 4500 V, the nebulizer pressure was 2 bar, with a drying gas flow of 10 L/min at 220°C, while the sheath gas (N2) flow was 4 L/min at 400°C. For mass accuracy, the instrument was calibrated using a combination of ESI Tune Mix and sodium formate, providing well-defined cluster ions across the *m/z* ranges to ensure accurate alignment of mass.

**Synthesis of synthetic standards**

All reactions were conducted in a fume hood. Furthermore, all reactions were carried out in flame-dried glassware and under an atmosphere of argon unless otherwise stated. Cooling to 0 °C was done using an ice bath and cooling to -78°C was done using a dry ice acetone bath. Heating was done using a polyethyleneglycol 400 (PEG-400) bath. $CH_2Cl_2$ and THF were dried over aluminium oxide via an MBrain SPS-800 solvent purification system and either used directly or stored over preactivated molecular sieves (4 Å). DMF, DMSO and DCE were purchased anhydrous. The dryness of solvents was controlled via Karl Fischer titration. $Et_3N$ and $(iPr)_2NEt$ were dried by stirring for at least 30 minutes over $CaH_2$ followed by distillation onto preactivated molecular sieves (4 Å). Reagents were used as received from commercial suppliers unless otherwise stated (Sigma Aldrich, Merck, AK Scientific, Fluorochem and TCI). Molecular sieves were activated by drying in the oven at 120 °C for a minimum of 24 hours, before they were heated in the microwave at maximum power for 2 minutes followed by 20 minutes under high vacuum to remove the formed vapour. This was repeated 3-4 times. Concentration *in vacuo* was performed using a rotary evaporator with the water bath temperature at 35 °C, followed by further concentration using a high vacuum pump. TLC analysis was carried out on silica coated aluminium foil plates (Merck Kieselgel 60 F254). The TLC plates were visualized by UV irradiation and staining with $KMnO_4$ stain ($KMnO_4$(5.0 g), 5 % aq. NaOH (8.3 mL) and $K_2CO_3$ (33.3 g) in $H_2O$ (500 mL)) or vanillin stain (vanillin (30.0 g), EtOH

(500 mL), conc. $H_2SO_4$(5.0 mL)). Automated flash column chromatography (AFCC) was carried out with Interchim Puriflash 420 or 5.050 using prepacked silica gel columns (30 µm particle size). Preparative HPLC was carried out using Interchim PuriFlash 5.250p using Interchims C18-HQ Prep-LC column (5 µm particle size, 150×10.0 mm). Infrared spectra (IR) obtained on a PerkinElmer Spectrum TwoTM UATR. Mass spectra (HRMS) were recorded on a Bruker Daltonics MicrOTOF time-of-flight spectrometer with positive electrospray ionization, or negative ionization when stated. Nuclear magnetic resonance (NMR) spectra were acquired on a Varian Mercury 400 MHz spectrometer. $^{1}$H NMR, $^{13}$C NMR and $^{31}$P NMR data were collected at 400 MHz, 101 MHz and 162 MHz respectively. The Chemical shifts ($\delta$) are reported in ppm relative to solvent signals, as reported by Gotlieb *et al.*[3] $^{31}$P NMR is reported relative to neat $H_3PO_4$ (0.00 ppm). Data for NMR resonances are given as multiplicity (s (singlet), d (doublet), t (triplet), q (quartet), p (pentet), m (multiplet), br. (broad)) and coupling constants *J* in Hz.

For detailed synthetic procedures and characterization data, see **Synthetic procedures**.

**Bioactivity assays**

To test the bioactivity of **3** and **10**, we performed both the standard minimum inhibitory concentration determinations and spot tests.

For minimum inhibitory concentration determinations, a stock solution of **3** and **10** (DMSO, 12.8 mg/ml) was used to make a 2-fold dilution series of each with Mueller-Hinton Broth in 96-well plates. The final concentrations of compound spanned 256 µg/ml to 0.25 µg/ml. Here, the test strain *Staphylococcus aureus* DSM 20231 (5x$10^6$ CFU/mL) was added, and the plates were incubated either aerobically or anaerobically overnight (37°C). If grown anaerobically, the plate was transferred to a sealed box containing a Microbiology Anaerocult A bag, and incubated. The well with the lowest compound concentration that showed no visual growth was designated the minimal inhibitory concentration (MIC). It was found that compound **3** precipitated within the wells at high concentrations (256 µg/ml to 16 µg/ml) and that **10** precipitated in lower degree (256 µg/ml to 64 µg/ml).

For the spot tests, we used an overnight culture of *S. aureus* DSM 20231 (37°C, 140 rpm, 16-18 h) in MHB. If tested on MHB with 25 mM $NaHCO_3$, both the overnight culture and the media used for dilution contained the same concentration. The culture was either cast into MHA (± 25 mM $NaHCO_3$) or spread across the agar plate using a cotton swab. In the plates containing *S. aureus* cast into the agar, the final OD was 0.008. In the plates containing *S. aureus* spread across the agar plate, the culture was first diluted to OD 0.008, and then the diluted culture was the one spread across the agar plate. On the plates, the following was spotted (10 µl): **3** (100 µg/ml), **10** (100 µg/ml), cell extract from NRPS/PKS-2, NRPS/PKS-2 W148A, and NRPS-5b, along with BE-43547 (10 µg/ml), DMSO, and MeOH. Compounds **3**, **10**, and BE-43547 were dissolved in DMSO. The cell extracts were dissolved in MeOH. All plates were incubated either aerobically or anaerobically overnight (37°C). If grown anaero-bically, the plate was transferred to a sealed box containing a Microbiology Anaerocult A bag and incubated.

**Point mutations in the DH and DHt domain**

All mutants created in this study were generated by PCR, introducing point mutations in either the DH or DHt domain using the original plasmid (pMT011) as the template. The same approach using Phusion high-fidelity DNA polymerase for amplification and implementation of the point mutation was applied. Similarly, the construction of the plasmid with NEBuilder HiFi DNA assembly, transformation, and purification was done as described previously. The primers used to target specific amino acid residues are found in **Table S4**. Heterologous ex-pression was done using the 24-deep well format as described previously. The amino acids selected for mutations were based on two criteria: 1) It was a histidine residue. 2) Or if the amino acid was located spatially favorably within or near the active site, and/or could have a supporting function to the catalytic active amino acids.

#### Tables

**Table S1: Media components used throughout the study.**

| Medium | Name | Amount | Ingredient |
|---|---|---|---|
| LB | Lysogeny broth (1 L) | 25 g | Premixed LB |
| | | 15 g | Agar if needed |
| MHA | Mueller-Hinton agar (1 L) | 38 g | Premixed MHA |
| MHB | Mueller-Hinton broth (1 L) | 21 g | Premixed MHB |
| SOC | Super Optimal Broth (1 L) | 20 g<br>5 g<br>0.5 g<br>0.186<br>Autoclave sterilisation then add | Tryptone<br>Yeast extract<br>NaCl<br>KCl<br>pH adjusted to 7 |
| | | | 20 mM sterile glucose<br>10 mM $MgCl_2$ |
| XPP3[4] | XPP3 (1 L) | 10 g<br>20 ml<br>20 ml<br>20 g<br>Add $dH_2O$<br>Autoclaved | Glycerin<br>M9 salt A<br>M9 salt B<br>Tryptone Type P |
| | | | Before use, add from sterile stocks<br>0.2% vitamin solution<br>0.1% trace metal solution<br>0.04% L-arabinose |
| | Salt A (1 L) | 350 g<br>100 g | $K_2HPO_4$<br>$KH_2PO_4$ |
| | Salt B (1 L) | 29.4 g<br>50 g<br>5 g | Sodium citrate<br>$(NH_4)_2SO_4$<br>$MgSO_4$ |
| | Vitamin solution (1 L) | 10 mg<br>6 mg<br>200 mg<br>1 g<br>1.2 g<br>2.3 g<br>12 g<br>20 mg | Folic acid<br>Biotin<br>p-aminobenzoic acid<br>Thiamine-HCl<br>Panthothenic acid<br>Nicotinic acid<br>Pyridoxine HCl<br>Vitamine B12<br>Sterile filtration |

| | | | |
|---|---|---|---|
| | Trace element solution (1 L) | 40 mg<br>200 mg<br>10 mg<br>10 mg<br>10 mg<br>10 mg | $ZnCl_2$<br>$FeCl_3 \bullet 6H_2O$<br>$CuCl_2 \bullet 2H_2O$<br>$MnCl_2 \bullet 4H_2O$<br>$Na_2B_4O_7 \bullet 10H_2O$<br>$(NH_4)_6Mo_7O_{24} \bullet 4H_2O$ |

**Table S2: Overview of plasmids used for NRPS engineering.**

| NRPS | Plasmids | Genotype | Reference |
|---|---|---|---|
| | pCK_1134-a<br>pCK_1099-c* | ori p15A, *cm*$^{R}$, *araC- PBAD, xfpS* C1A1$^{Leu}$T1$^{I}$ modules 2-3 fitAB<br>ori ColA, *kan*$^{R}$, *araC- PBAD*, modules *4-6 fitAB* | [5] |
| | pCPU1 | ori p15A, *cm*$^{R}$, *araC- PBAD, xldAB* C1A1$^{ser}$T1CEA2 *xtpS* T2$^{I}$ modules 2-3 | Un-published |
| | pMT078 | ori ColA, kan$^{R}$, *araC- PBAD* | This study |
| -1 | pMT057 | ori p15A, *cm*$^{R}$, *araC- PBAD, xfpS* C1A1$^{Leu}$T1$^{IV}$ *BE-43547* T5$^{IV}$ | This study |
| -2 | pMT036 | ori p15A, *cm*$^{R}$, *araC- PBAD, xfpS* C1A1$^{Leu}$T1$^{I}$ *BE-43547* T5$^{I}$C2A2$^{ser}$T6 | This study |
| -3a | pMT046 | ori p15A, *cm*$^{R}$, *araC- PBAD, xfpS* C1A1$^{Leu}$T1$^{I}$ codon optimized *BE-43547* T5$^{I}$C2A2$^{ser}$T6 | This study |
| -3b | pMT045 | ori p15A, *cm*$^{R}$, *araC- PBAD, xfpS* C1A1$^{Leu}$T1$^{IV}$ codon optimized *BE-43547* T5$^{IV}$C2A2$^{ser}$T6 | This study |
| -4a | pMT035 | ori p15A, *cm*$^{R}$, *araC- PBAD, xldAB* C1A1$^{Ser}$T1$^{I}$ *BE-43547* T5$^{I}$ | This study |
| -4b | pMT024 | ori p15A, *cm*$^{R}$, *araC- PBAD, xldAB* C1A1$^{Ser}$T1$^{IV}$ *BE-43547* T5$^{IV}$ | This study |
| | pMT011 | ori ColA, *kan*$^{R}$, *araC- PBAD*, codon optimized *BE-43547* module 6-7 and TE | This study |
| /PKS-1 | pMT035<br>pMT011 | | This study |
| /PKS-2 | pMT024<br>pMT011 | | This study |
| /PKS-3 | pMT057<br>pMT011 | | This study |
| -4b + pCOLA | pMT024<br>pMT078 | ori p15A, *cm*$^{R}$, *araC- PBAD, xldAB* C1A1$^{Ser}$T1$^{IV}$ *BE-43547* T5$^{IV}$<br>ori ColA, *kan*$^{R}$, *araC- PBAD* | This study |

**Table S3: Overview of plasmids used for point mutation studies.**

| Plasmids | Function | Reference |
|---|---|---|
| pMT058 | ori ColA, *kan*$^R$, *araC- PBAD*, codon opti-mized *BE-43547* module 6-7 | This study |
| pMT059 | pMT011 mutation in DHt, H105A | This study |
| pMT060 | pMT011 mutation in DHt, H145A | This study |
| pMT061 | pMT011 mutation in DHt, H166A | This study |
| pMT062 | pMT011 mutation in DHt, H181A | This study |
| pMT063 | pMT011 mutation in DH, H10A | This study |
| pMT064 | pMT011 mutation in DH, H15A | This study |
| pMT065 | pMT011 mutation in DH, H34A | This study |
| pMT066 | pMT011 mutation in DH, D55A | This study |
| pMT067 | pMT011 mutation in DH, H88A | This study |
| pMT069 | pMT011 mutation in DH, H127A | This study |
| pMT070 | pMT011 mutation in DH, H145A | This study |
| pMT071 | pMT011 mutation in DH, H155A | This study |
| pMT072 | pMT011 mutation in DH, H181A | This study |
| pMT073 | pMT011 mutation in DH, D186A | This study |
| pMT074 | pMT011 mutation in DH, D8A | This study |
| pMT075 | pMT011 mutation in DH, Q57A | This study |
| pMT076 | pMT011 mutation in DH, F142A | This study |
| pMT077 | pMT011 mutation in DH, R193AA | This study |
| pMT088 | pMT011 mutation in DH, W148A | This study |

**Table S4: Primers used for plasmid constructions and sequencing.**

| Plasmid | Primer | Sequence (5' → 3') | Template |
|---|---|---|---|
| | T7term | TGCTAGTTATTGCTCAGCGG | Sequencing primer of NRPS plasmids |
| | pBAD-for | ATGCCATAGCATTTTTATCC | Sequencing primer of NRPS plasmids |
| | oMT_seq1_for | TGGGCCACCCATTGTTACAG | Sequencing primer for DH mutants |
| | oMT_seq4_rev | TGCTAGTTATTGCTCAGCGG | Sequencing primer of NRPS plasmids |
| | oMT_seq5_for | ATGCCATAGCATTTTTATCC | Sequencing primer NRPS plasmids |
| pMT078 | oMT174R | TGGCAG-CAGCCTAGGTTAATGGAATTCCTCCTGTTAGCCCAAAAAAACG | pCK_1099-c* |
| | oMT175F | ATTAACCTAGGCTGCTGCCACCGC | |
| pMT011 | oMT002R | CATGGAATTCCTCCTGTTAGCCCAAAAAAACGG | pMT078 |
| | oMT067F | TAAATTAACCTAGGCTGCTGCCACCG | |

| | | | |
|---|---|---|---|
| | oMT003F | TTTTTTTGGGCTAACAGGAG-GAATTCCATGTCCGACAGCACAGATGAT-TTAAGTCAGGTCG | Synthetic fragment 1 from BE-43547 BGC |
| | oMT004R | AAAAGTCTCAACAGTAGGAGTGGAAC-GCAGG | |
| | oMT005F | CTGCGTTCCACTCCTACTGTTGA-GACTTTTC | Synthetic fragment 2 from BE-43547 BGC |
| | oMT006R | GGTGCGATATAAACGGTCACCAAGTCCG | |
| | oMT007F | ATCGGACTTGGTGACCGTTTATATCG-CACC | Synthetic fragment 3 from BE-43547 BGC |
| | oMT008R | TCAGCGGTGGCAG-CAGCCTAGGTTAATTTAACGCAC-CTGGTCACAGACTGGAGAC | |
| pMT024 | oCPU5F | TGACAATTAATCATCGGCTCG-TATAATGTGTGG | pCK_1134-a |
| | oCPU6R | CATGGAATTCCTCCTGTTAGCCCAAAAAAACG | |
| | oMT045F | AGGACACTCACTGCTGATGGCGG | gDNA *Micromonospora* sp. RV43 |
| | oMT046R | CACATTATACGAGCCGATGATTAATT-GTCACTCCCCTGGGGATTCGGTACG | |
| | oCPU1F | TTTTTTTGGGCTAACAGGAG-GAATTCCATGAATACACAGCGTAAC-CACAATCCATCATCC | pCPU1 |
| | oMT044R | GCAGTTCCGCCATCAGCAGTGAG-TGTCCTCCCAG-TTCAAAAAAGTGGTCATGGCGAC | |
| pMT035 | oCPU5F | TGACAATTAATCATCGGCTCG-TATAATGTGTGG | pCK_1134-a |
| | oCPU6R | CATGGAATTCCTCCTGTTAGCCCAAAAAAACG | |
| | oMT073F | TACGAGGCGCCGCGCGACGAG | gDNA *Micromonospora* sp. RV43 |
| | oMT046R | CACATTATACGAGCCGATGATTAATT-GTCACTCCCCTGGGGATTCGGTACG | |
| | oMT069F | CCGTTTTTTTGGGCTAACAGGAG-GAATTC | pCPU1 |
| | oMT072R | CTCGTCGCGCGGCGCCTCGTAATCAC-GGGTGAGCACGGAGGATAAG | |
| pMT036 | oMT069F | CCGTTTTTTTGGGCTAACAGGAG-GAATTC | pCK_1134-a |
| | oMT102R | GTAGGGAGTCTCCGGGCCG-GAGCGTCCG-TAAACATAGCGGCTCTGTTTAAAATCTGGCTC | |
| | oMT101F | TACGGACGCTCCGGCCCGGAGAC | gDNA *Micromonospora* sp. RV43 |
| | oMT046R | CACATTATACGAGCCGATGATTAATT-GTCACTCCCCTGGGGATTCGGTACG | |
| | oCPU5F | TGACAATTAATCATCGGCTCG-TATAATGTGTGG | pCK_1134-a |
| | oCPUC6R | CATGGAATTCCTCCTGTTAGCCCAAAAAAACG | |
| pMT045 | oMT069F | CCGTTTTTTTGGGCTAACAGGAG-GAATTC | pCK_1134-a |

| | | | |
|---|---|---|---|
| | oMT094R | GCAACGCGTGTCGCAA-GCAGACTGTGCCCGCCGATACGGAA-GAAGTTATCTTCAACG | |
| | oMT082F | CGGGCACAGTCTGCTTGCGAC | Synthetic fragment 4 from BE-43547 BGC |
| | oMT086R | CACATTATACGAGCCGATGATTAATT-GTCATTCGCCTGGCGATTCAGTAC-GAATTTC | |
| | oCPU5F | TGACAATTAATCATCGGCTCG-TATAATGTGTGG | pCK_1134-a |
| | oCPUC6R | CATGGAATTCCTCCTGTTAGCCCAAAAAAACG | |
| pMT046 | oMT069F | CCGTTTTTTTGGGCTAACAGGAG-GAATTC | pCK_1134-a |
| | oMT095R | GTTCTAATGGATCACGAGGAGCACGA-TAAACATAGCGGCTCTGTTTAAAATCTGGCTC | |
| | oMT077F | TATCGTGCTCCTCGTGATCCATTAG | Synthetic fragment 4 from BE-43547 BGC |
| | oMT086R | TGTTCTAATGGATCACGAGGAGCACGA-TAGCTGTCTTTGTTTCCCCATATCGG | |
| | oCPU5F | TGACAATTAATCATCGGCTCG-TATAATGTGTGG | pCK_1134-a |
| | oCPUC6R | CATGGAATTCCTCCTGTTAGCCCAAAAAAACG | |
| pMT057 | oMT045F | AGGACACTCACTGCTGATGGCGG | gDNA *Micromonospora* sp. RV43 |
| | oMT046R | CACATTATACGAGCCGATGATTAATT-GTCACTCCCCTGGGGATTCGGTACG | |
| | oMT052F | TTTTTTTGGGCTAACAGGAG-GAATTCCATGGA-TAACATTCTGGCCTCGCCATTTACAC | pMT045 |
| | oMT132R | GTTCCGCCATCAGCAGTGAG-TGTCCTCCGATACGGAA-GAAGTTATCTTCAACGC | |
| | oCPU5F | TGACAATTAATCATCGGCTCG-TATAATGTGTGG | pCK_1134-a |
| | oCPUC6R | CATGGAATTCCTCCTGTTAGCCCAAAAAAACG | |
| pMT058 | oMT005F | CTGCGTTCCACTCCTACTGTTGA-GACTTTTC | pMT011 |
| | oMT004R | AAAAGTCTCAACAGTAGGAGTGGAAC-GCAGG | |
| | oMT133R | CGGTGGCAGCAGCCTAGGTTAATTTAA-GCCGGTTGACCGGCCCCGTC | pMT011 |
| | oMT067F | TAAATTAACCTAGGCTGCTGCCACCG | |
| pMT059 | oMT137F | TTCCGTCAGTAGTTCGTG-TAGCTGCGGGCGCAGGGTTCG-CACAAGG | pMT011 |
| | oMT135F | GGATCGCAGTGGTGAGTAACCATGC | |
| | oMT134R | GCATGGTTACTCACCACTGCGATCC | pMT011 |
| | oMT136R | GCTACACGAACTACTGACGGAAGC | |
| pMT060 | oMT138R | GCCACAACGCGTGCCGGGGGACG | pMT011 |
| | oMT139F | GTCCCCCGGCACGCGTTGTGGCCTT-GTGGTCGTTAGCGGTAGAGCCC | |
| pMT061 | oMT140R | GCAAAACCTACTGCTTCCGCGCGG | pMT011 |

| | | | |
|---|---|---|---|
| | oMT141F | GCGCGGAAGCAGTAGGTTTTGCCAG-TCTGTTGGCATTGGCTCAAGC | |
| pMT062 | oMT142R | GCCTCGGCACGATCGGCAAGAGC | pMT011 |
| | oMT143F | CTCTT-GCCGATCGTGCCGAGGCTCCCGCGTT-GACTATCGAAGTTC | |
| pMT063 | oMT144R | GCTTCATCCACAATCCAAGAATCGGCC | pMT011 |
| | oMT145F | GATTCTTGGATTGTGGATGAAGCTCG-CATGCAGGGTCACGGATTGG | |
| pMT064 | oMT146R | GCACCCTGCATGCGATGTTCATCC | pMT011 |
| | oMT147F | GATGAACATCGCATGCAGGGTGCCG-GATTGGTGCCCGGAACAACG | |
| pMT065 | oMT148R | GCACGTGCGACAGCAGCGCGAACC | pMT011 |
| | oMT149F | CGCGCTGCTGTCGCAC-GTGCCGCCGATGGGCGCGAGATTG | |
| pMT066 | oMT150R | GCTGGAACGATTACGGGGGACGTG | pMT011 |
| | oMT151F | GTCCCCCGTAATCGTTCCAGCCGACCAG-GAACGCGAGATGC | |
| pMT067 | oMT152R | GCTTCCTGGCGGCCGGCAGCG | pMT011 |
| | oMT153F | GCTGCCGGCCGCCAGGAA-GCCTGTGCCGGAACAGTAGTATTGC | |
| pMT069 | oMT156R | GCACGCAGAGCTGCCTCGCCC | pMT011 |
| | oMT157F | GGCGAGGCAGCTCTGCGTGCCCGTCTGC GTTTGGATTTTGC | |
| pMT070 | oMT158R | GCACGCAGAGCTGCCTCGCCC | pMT011 |
| | oMT159F | GGCGAGGCAGCTCTGCGTGCCCGTCTGC GTTTGGATTTTGC | |
| pMT071 | oMT160R | GCCACAGCAAAGCGGATCAAACC | pMT011 |
| | oMT161F | GTTTGATCCGCTTT-GCTGTGGCTGGGCGTT-GGCGCTCTTTAAGC | |
| pMT072 | oMT162R | GCTACGCGGCTTAAAGAGCGCC | pMT011 |
| | oMT163F | GCGCTCTTTAAGCCGCGTAGCCGTGG-GAACGACAGGAATGGTGG | |
| pMT073 | oMT164R | GCTAAAAGA-TAGGTGTCTAAATCTCCGGC | pMT011 |
| | oMT165F | GATTTAGACAC-CTATCTTTTAGCTCCCGCCTTACTGGAC-GTAGTTGG | |
| pMT074 | oMT166R | GCCACAATCCAAGAATCGGCCG | pMT011 |
| | oMT167F | GCCGATTCTT-GGATTGTGGCTGAACATCG-CATGCAGGGTCACG | |
| pMT075 | oMT168R | GCGTCGTCTGGAACGATTACGGGG | pMT011 |
| | oMT169F | GTAATCGTTCCAGACGACGCGGAAC-GCGAGATGCTGACGACC | |
| pMT076 | oMT170R | GCGCGGATCAAACCTCCGTCGGC | pMT011 |
| | oMT171F | GACGGAGGTTTGATCCGCGCTGCTGTG-CATGGGCGTTGGC | |
| pMT077 | oMT172R | GCGCTCGCGCCGCCAACTACGTCC | pMT011 |
| | oMT173F | GTAGTTGGCGGCGCGAGCGCTGTG-TATGCGGCCGAAGGGTATTATTTACC | |

| | | | |
|---|---|---|---|
| pMT088 | oMT182F | TTTGCTGTG-CATGGGCGTGCGCGCTCTTTAA-GCCGCGTACACG | pMT011 |
| | oMT183R | GCACGCCCATGCACAGCAAAGC | |

**Table S5: Overview of microorganisms used.**

| **Strain** | **Source** |
|---|---|
| *E. coli* DH5-α | In-house |
| *E. coli* DH10B::*mtaA*[6] | In-house |
| *Micromonospora* sp. RV43[7] | Kindly obtained from Prof. Dr. Ute Hentschel |
| *Staphylococcus aureus* DSM 20231 | DSMZ |
| *Xenorhabdus bovienii* SS-2004[8] | In-house |
| *Xenorhabdus stockiae* DSM 17904 | DSMZ |

#### Synthetic procedures

##### Methyl N-(tert-butoxycarbonyl)-N-methylglycinate (4)

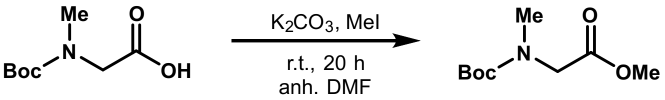


To a flame-dried flask containing compound *N*-Boc-sarcosine (5002 mg, 26.44 mmol, 1.0 equiv.) and $K_2CO_3$ (6211 mg, 44.94 mmol, 1.7 equiv.) in anh. DMF (35 mL, 0.75 M) was added MeI (2.30 mL, 37.0 mmol, 1.4 equiv.). The reaction mixture was stirred at r.t. for 20 h. After full consumption of *N*-Boc-sarcosine as indicated by TLC (90:8:2 $CH_2Cl_2$/EtOAc/ Formic Acid) the reaction mixture was diluted using water (40 mL) and $CH_2Cl_2$ (40 mL). The aqueous layer was extracted using $CH_2Cl_2$ (2 × 40 mL). The organic phases were combined and washed using aq. 1M LiCl (40 mL). The organic phase was collected, dried over anh. $Na_2SO_4$ and concentrated *in vacuo*. The crude mixture was purified using AFCC (100% heptane to 3:1 heptane/EtOAc) to afford compound **4** (4297 mg, 21.14 mmol, 80%) as a yellow oil.

$R_f$ (4:1 heptane/EtOAc, $KMnO_4$ stain) 0.26

$\tilde{\nu}_{max}$ (ATR) 2976, 1752, 1694, 1390, 1366, 1207, 1146, 881, 777

HRMS (ESI) Calc. for $C_9H_{17}NNaO_4^+$ 226.1050; found 226.1052

The compound was found to exist as a 1:1 mixture of rotamers in $CDCl_3$ at room temperature.

$^1$H NMR (400 MHz, Chloroform-d) $\delta_H$ (ppm) 3.98 and 3.90 (s, 2H), 3.74 and 3.73 (s, 3H), 2.93 and 2.91 (s, 3 H), 1.47 and 1.42 (s, 9H).

$^{13}$C NMR (101 MHz, Chloroform-d) $\delta_C$ (ppm) 170.6, 156.2 and 155.6, 80.3, 52.1, 51.1 and 50.3, 35.7, 28.5 and 28.4.

Data is in accordance with literature.[9,10]

**Methyl N-(2-bromoacetyl)-N-methylglycinate (6)**

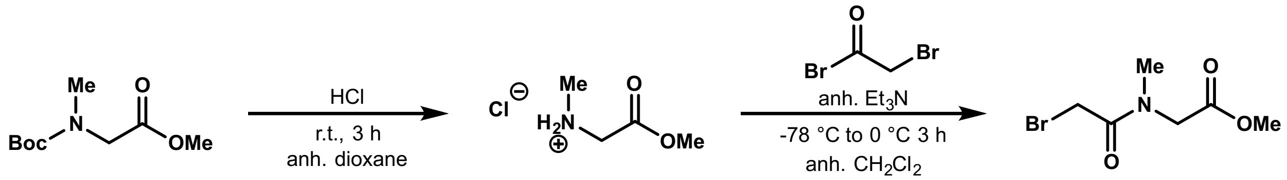


A flame-dried flask containing compound **4** (1016 mg, 4.999 mmol, 1.0 equiv.) equipped with a magnetic stirring bar, was azeotroped with anh. PhMe (2 × 30 mL). The flask was placed under high vacuum for 30 min and backfilled with argon. A solution of 4 M HCl in 1,4-dioxane (25 mL, 0.2 M) was added and the reaction mixture was stirred at r.t. for 3 h. After full consumption of compound **4** as indicated by TLC (100% EtOAc) the mixture was concentrated *in vacuo*. The crude mixture was then co-evaporated with $Et_2O$ two times and placed under high vacuum for 30 min. The hydrochloride salt was then re-dissolved in anh. $CH_2Cl_2$ (30 mL, 0.16 M) and was added anh. $Et_3N$ (2.10 mL, 15.1 mmol, 3.0 equiv). The reaction mixture was cooled to -78 °C and a solution of bromoacetyl bromide (653 µL, 7.51 mmol, 1.5 equiv.) in anh. $CH_2Cl_2$ (10 mL, 0.75 M) was added dropwise over a period of 5 min. The reaction mixture was stirred at -78 °C for 30 min and was allowed to warm to 0 °C over 3 hours. After full consumption of the hydrochloride salt as indicated by TLC (100% EtOAc) the reaction mixture was diluted using $CH_2Cl_2$ (100 mL) and washed using 1 M aq. HCl (50 mL). The aqueous phase was back-extracted using $CH_2Cl_2$ (50 mL). The combined organic phases were washed using brine (75 mL), and the aqueous phase was back-extracted using $CH_2Cl_2$ (50 mL). The organic phases were combined, dried over anh. $Na_2SO_4$ and concentrated *in vacuo*. The crude mixture was purified using AFCC (4:1 heptane/EtOAc to 2:3 heptane/EtOAc) to afford compound **6** (1085 mg, 4.843 mmol, 97 %) as an orange oil.

$R_f$ (1:1 heptane/EtOAc, UV, $KMnO_4$ stain) 0.26

$\tilde{\nu}_{max}$ (ATR) 2953, 1743, 1650, 1400, 1208, 1090

HRMS (ESI) Calc'd m/z for $C_6H_{10}BrNNaO_3^+$ 245.9737 $[M+Na]^+$; found 245.9740

The compound was found to exist as a 4:1 mixture of rotamers in $CDCl_3$ at room temperature. The NMR-data for each rotamer is given separately below.

**Major rotamer:**

$^{1}H$ NMR (400 MHz, Chloroform-d) $\delta_H$ (ppm) 4.14 (s, 2H), 3.92 (s, 2H), 3.75 (s, 3H), 3.18 (s, 3H).

$^{13}C$ NMR (101 MHz, Chloroform-d) $\delta_C$ (ppm) 169.3, 167.5, 52.4, 49.8, 37.5, 25.8.

**Minor rotamer:**

$^{1}H$ NMR (400 MHz, Chloroform-d) $\delta_H$ (ppm) 4.16 (s, 2H), 3.80 (s, 2H), 3.80 (s, 3H), 3.01 (s, 3H).

$^{13}C$ NMR (101 MHz, Chloroform-d) $\delta_C$ (ppm) 169.1, 167.2, 52.8, 52.2, 35.6, 25.8.

Data is in accordance with literature.[11,12]

**Methyl N-(2-(diethoxyphosphoryl)acetyl)-N-methylglycinate (7)**

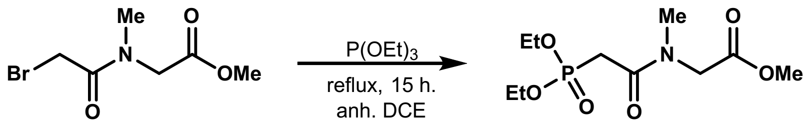


A flame-dried flask containing compound **6** (1354 mg, 6.043 mmol, 1.0 equiv.) was azeotroped using anh. PhMe (2 × 30 mL). The flask was placed under high vacuum for 30 min and backfilled with argon. Anh. DCE (6 mL, 0.8 M) was added, followed by $P(OEt)_3$ (2.09 mL, 12.1 mmol, 2.0 equiv.) The flask was fitted with a reflux condenser, heated to reflux (ca. 90 °C) and stirred for 15 h. After full consumption of compound **6** as indicated by TLC (100% EtOAc), the reaction mixture was cooled to r.t. and concentrated *in vacuo*. The crude oil was triturated using pentane (3 × 4 mL) and the resulting crude purified using AFCC (100% $CH_2Cl_2$ to 9:1 $CH_2Cl_2$/MeOH) to afford compound **7** (1485 mg, 5.280 mmol, 87%) as a yellow oil.

$R_f$ (9:1 $CH_2Cl_2$/MeOH, UV, $KMnO_4$ stain) 0.49

$\tilde{\nu}_{max}$ (ATR) 2984, 2952, 1747, 1648, 1249, 1209, 1048, 1019, 966

HRMS (ESI) Calc'd m/z for $C_{10}H_{20}NNaO_6P^+$ 304.0921 $[M+Na]^+$; found 304.0922

$^{31}$P NMR (162 MHz, Chloroform-d) $\delta_P$ (ppm) 21.20

The compound was found to exist as a 3:1 mixture of rotamers in $CDCl_3$ at room temperature. The NMR-data for each rotamer is given separately below.

**Major rotamer:**

$^{1}$H NMR (400 MHz, Chloroform-d) $\delta_H$ (ppm) 4.30 – 4.10 (m, 6H), 3.72 (s, 3H), 3.19 (s, 3H), 3.09 – 2.94 (m, 2H), 1.34 (t, J = 7.0 Hz, 6H).

$^{13}$C NMR (101 MHz, Chloroform-d) $\delta_C$ (ppm) 169.5, 165.8 (d, J = 5.5 Hz), 62.8 (d, J = 6.4 Hz), 52.3, 49.7, 37.9, 32.9 (d, J = 37.7 Hz), 16.5 (d, J = 6.3 Hz).

**Minor rotamer:**

$^{1}$H NMR (400 MHz, Chloroform-d) $\delta_H$ (ppm) 4.30 – 4.10 (m, 6H), 3.77 (s, 3H), 3.14 (s, 3H), 3.09 – 2.94 (m, 2H), 1.33 (t, J = 7.0 Hz, 6H).

$^{13}$C NMR (101 MHz, Chloroform-d) $\delta_C$ (ppm) 169.7, 165.4 (d, J = 5.5 Hz), 62.9 (d, J = 6.4 Hz), 52.6, 52.4, 35.5, 34.3 (d, J = 37.7 Hz), 16.4 (d, J = 6.3 Hz).

Data is in accordance with literature.[12]

**N-(1,3-dihydroxypropan-2-yl)tetradecanamide (9)**

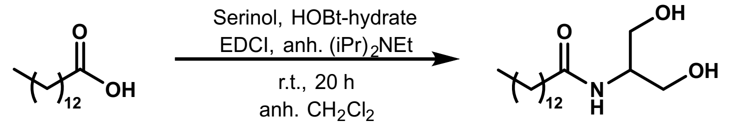


To a flame-dried flask containing myristic acid (3007.4 mg, 13.169 mmol, 1.0 equiv.) in anh. $CH_2Cl_2$ (70 mL, 0.2 M) was added serinol (1260 mg, 13.83 mmol, 1.05 equiv.). The mixture was cooled to 0 °C and sequentially added HOBt-hydrate (2420 mg, 15.80 mmol, 1.2 equiv.), EDCI (3029 mg, 15.80 mmol, 1.2 equiv.) and anh. $(iPr)_2NEt$ (2.75 mL, 15.8 mmol, 1.2 equiv.). The mixture was allowed to heat to r.t. and stirred for 20 h. After full consumption of myristic acid as indicated by TLC (90:8:2, $CH_2Cl_2$/EtOAc/Formic acid) the white precipitate was filtered and washed using ice cold $CH_2Cl_2$. The solid was collected and recrystallized from EtOH (10 mL). The resulting precipitate was filtered and washed using ice cold EtOH to afford compound **9** (3564 mg, 11.82 mmol, 90 %) as a white solid.

R$_f$ (9:1 $CH_2Cl_2$/MeOH, $KMnO_4$ stain) 0.46

$\tilde{\nu}_{max}$ (ATR) 3297, 2955, 2917, 2849, 1639, 1543, 1071,1057, 973, 678

HRMS (ESI) Calc'd m/z for $C_{17}H_{36}NO_3^+$ 302.2690; found 302.2693

$^1$H NMR (400 MHz, DMSO-d$_6$) $\delta_H$ (ppm) 7.44 (br. d, J = 8.1 Hz, 1H), 4.56 (br. s, 2H), 3.68 (dp, J = 8.0, 5.7 Hz, 1H), 3.37 (d, J = 5.7 Hz, 4H), 2.05 (t, J = 7.4 Hz, 2H), 1.45 (p, J = 7.0 Hz, 2H), 1.23 (s, 20H), 0.85 (t, J = 6.9 Hz, 3H).

$^{13}$C NMR (101 MHz, DMSO-d$_6$) $\delta_C$ (ppm) 172.1, 60.2, 52.7, 35.4, 31.3, 29.1, 29.1, 29.1, 29.0, 28.9, 28.8, 28.7, 25.4, 22.1, 14.0.

**N-(3-oxoprop-1-en-2-yl)tetradecanamide (8)**

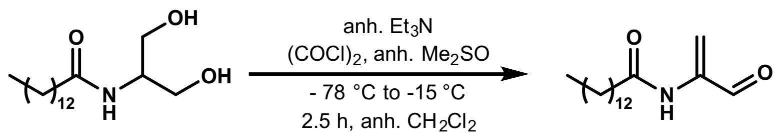


To a flame-dried flask containing $(COCl)_2$ (224 µL, 2.65 mmol, 2.0 equiv.) in anh. $CH_2Cl_2$ (3.0 mL, 0.9 M) was cooled to -78 °C and added a solution of anh. $Me_2SO$/$CH_2Cl_2$ (1:1, 1.9 mL) dropwise. The mixture was stirred at -78 °C for 10 min. Then compound **9** (399.7 mg, 1.326 mmol, 1.0 equiv.) dissolved in anh. $Me_2SO$/$CH_2Cl_2$ (2:1, 5.3 mL, 0.25 M) was added dropwise and stirred at -78 °C for 20 min. Then anh. $Et_3N$ (924 µL, 6.63 mmol, 5.0 equiv.) was added. The reaction mixture was stirred at -78 °C for another 5 min. and was then allowed to gradually (ca. 120 min) warm to -15 °C. At this point full consumption of compound **9** as indicated by TLC (9:1 $CH_2Cl_2$/MeOH). The reaction mixture was diluted using water (30 mL) and $CH_2Cl_2$ (30 mL) and the organic phase was separated. The aqueous phase was extracted with $CH_2Cl_2$ (2 × 30 mL). The combined organic phases were washed with 10% citric acid (20 mL) and brine (30 mL). The organic phase was collected, dried over anh. $Na_2SO_4$ and concentrated in vacuo. The crude mixture was purified by silica plug (1:1 heptane/EtOAc) to afford compound **8** (176.6 mg, 0.6275 mmol, 47 %) as a white solid.

R$_f$ (4:1 heptane/EtOAc, UV, $KMnO_4$ stain) 0.39

$\tilde{\nu}_{max}$ (ATR) 3675, 3386, 2953, 2916, 2848, 1704, 1682, 1674, 1524, 902

HRMS (ESI) Calc. for $C_{23}H_{41}N_2O_4^+$ 409.3061; found 409.3064

$^{1}$H NMR (400 MHz, Chloroform-d) $\delta_H$ (ppm) 9.15 (s, 1H), 7.64 (br. s, 1H), 7.16 (s, 1H), 5.56 (d, J = 1.3 Hz, 1H), 2.33 (t, J = 7.8 Hz, 2H), 1.73 – 1.61 (m, 2H), 1.37 – 1.23 (m, 20H), 0.88 (t, J = 7.0 Hz, 3H).

$^{13}$C NMR (101 MHz, Chloroform-d) $\delta_C$ (ppm) 189.3, 172.3, 139.8, 118.2, 37.7, 32.1, 29.8, 29.8, 29.7, 29.6, 29.5, 29.5, 29.3, 25.4, 22.8, 14.3.

**Methyl (E)-N-methyl-N-(4-tetradecanamidopenta-2,4-dienoyl)glycinate (10)**

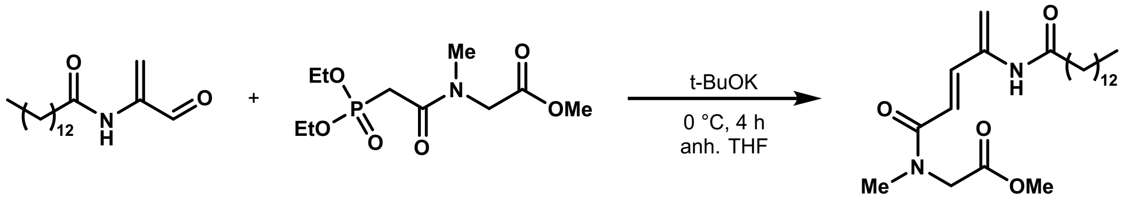


To a flame-dried flask containing compound **7** (55 mg, 0.195 mmol, 1.1 equiv.) in anh. THF (2.0 mL, 0.1 M) was cooled to -78 °C and added t-BuOK (1 M in anh. THF, 195 µL, 0.195 mmol, 1.1 equiv.) dropwise. The reaction mixture was stirred at -78 °C for 10 min and then added a solution of compound **8** (50 mg, 0.178 mmol, 1.0 equiv.) in anh. THF (900 µL, 0.2 M) dropwise. The reaction mixture was then allowed to warm to 0 °C and stirred for 4 h. After full consumption of compound **8** as indicated by TLC (2:1 heptane/EtOAc) the reaction mixture was diluted using EtOAc (15 mL) and washed using water (15 mL). The aqueous phase was extracted using EtOAc (2 × 15 mL). The organic phases were combined, dried over anh. $Na_2SO_4$ and concentrated *in vacuo*. The crude mixture was purified using AFCC (1:1 heptane/EtOAc to 100% EtOAc) to afford compound **10** (64.6 mg, 0.158 mmol, 89 %) as a white solid.

$R_f$ (100% EtOAc, UV, $KMnO_4$ stain) 0.55

$\tilde{\nu}_{max}$ (ATR) 3273, 2955, 2916, 2849, 1731, 1651, 1528, 1233, 1127, 975

HRMS (ESI) Calc. for $C_{23}H_{41}N_2O_4^+$ 409.3061; found 409.3064

The compound was found to exist as a 4:1 mixture of rotamers in $CD_2Cl_2$ at room temperature. The NMR-data for each rotamer is given separately below.

**Major rotamer:**

<u>$^{1}$H NMR</u> (400 MHz, Dichloromethane-d$_2$) $\delta_H$ (ppm) 7.20 (br. s, 1H), 7.07 (d, J = 15.3 Hz, 1H), 6.47 (d, J = 15.4 Hz, 1H), 5.95 (s, 1H), 5.21 (s, 1H), 4.15 (s, 2H), 3.71 (s, 3H), 3.13 (s, 3H), 2.30 (t, J = 7.6 Hz, 2H), 1.65 (p, J = 7.1 Hz, 2H), 1.37 – 1.24 (m, 20H), 0.87(t, J = 7.0 Hz, 3H).

<u>$^{13}$C NMR</u> (101 MHz, Dichloromethane-d$_2$) $\delta_C$ (ppm) 172.4, 170.1, 166.7, 141.2, 137.8, 116.6, 112.0, 52.4, 50.1, 37.7, 37.0, 32.3, 30.1, 30.1, 30.0, 30.0, 29.9, 29.8, 29.7, 29.7, 25.9, 23.1, 14.3.

**Minor rotamer:**

<u>$^{1}$H NMR</u> (400 MHz, Dichloromethane-d$_2$) $\delta_H$ (ppm) 7.14(br. s, 1H), 7.01 (d, J = 15.3 Hz, 1H), 6.28 (d, J = 15.4 Hz, 1H), 5.91 (s, 1H), 5.17 (s, 1H), 4.15 (s, 2H), 3.74 (s, 3H), 2.99 (s, 3H), 2.26 (t, J = 7.6 Hz, 2H), 1.65 (p, J = 7.1 Hz, 2H), 1.37 – 1.24 (m, 20H), 0.87 (t, J = 7.0 Hz, 3H).

<u>$^{13}$C NMR</u> (101 MHz, Dichloromethane-d$_2$) $\delta_C$ (ppm) 172.4, 169.9, 167.0, 140.7, 137.8, 116.9, 111.6, 52.8, 51.8, 37.7, 35.4, 32.3, 30.1, 30.1, 30.0, 30.0, 29.9, 29.8, 29.7, 29.7, 25.9, 23.1, 14.3.

**(E)-N-methyl-N-(4-tetradecanamidopenta-2,4-dienoyl)glycine (3)**

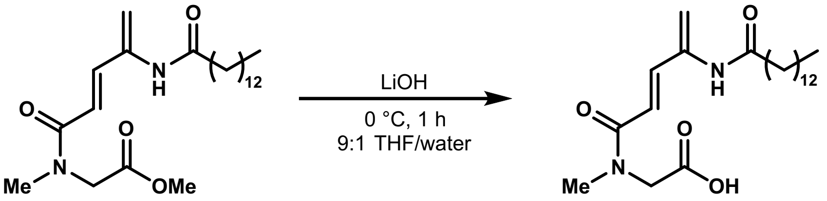


To a flask containing compound **10** (11.4 mg, 0.0278 mmol, 1.0 equiv.) in THF (1.0 mL, 0.028 M) was cooled to 0 °C and added 0.5 M aq. LiOH (111 µL, 0.0556 mmol, 2.0 equiv.). The reaction mixture was stirred at 0 °C for 1 h. After full consumption of compound **10** as indicated by TLC (95:4:1 EtOAc/MeOH/Formic acid) the reaction mixture was then diluted using 0.1 M aq. HCl (5.0 mL). The aqueous phase was then extracted using EtOAc (3 × 5.0 mL). The organic phases were combined and washed using brine (5.0 mL). The organic phase was collected, dried over anh. $Na_2SO_4$ and concentrated *in vacuo*. The crude mixture was purified using reverse phase AFCC (4:1 $H_2O$/MeCN to 100% MeCN with 0.1% formic acid additive) to afford compound **3** (6.5 mg, 0.017 mmol, 59 %) as a white solid.

$R_f$ (95:4:1 EtOAc/MeOH/Formic acid, UV, Vanillin stain) 0.51

HRMS (ESI) Calc'd m/z for $C_{22}H_{37}N_2O_4^-$ 393.2758 [M-H]$^-$; found 393.2751

The compound was found to exist as a 3:2 mixture of rotamers in DMSO-d$_6$ at room temperature. The NMR-data for each rotamer is given separately below. Not enough signal was achieved to characterise the compound using $^{13}$C NMR

**Major rotamer:**

$^1$H NMR (400 MHz, DMSO-d$_6$) $\delta_H$ (ppm) 13.01 (br. s, 1H), 8.99 (s, 1H), 6.91 (d, *J* = 15.7 Hz, 1H), 6.81 (d, *J* = 15.4 Hz, 1H), 5.91 (s, 1H), 5.20 (s, 1H), 4.05 (s, 2H), 3.13 (s, 3H), 2.31 (t, *J* = 7.8 Hz, 2H), 1.58 – 1.48 (m, 2H), 1.29 – 1.19 (m, 20H), 0.85 (t, *J* = 6.6 Hz, 3H).

**Minor rotamer:**

$^1$H NMR (400 MHz, DMSO-d$_6$) $\delta_H$ (ppm) 13.01 (br. s, 1H), 8.91 (s, 1H), 6.87 (d, *J* = 15.7 Hz, 1H), 6.67 (d, *J* = 15.4 Hz, 1H), 5.92 (s, 1H), 5.15 (s, 1H), 4.22 (s, 2H), 2.87 (s, 3H), 2.29 (t, *J* = 7.8 Hz, 2H), 1.58 – 1.48 (m, 2H), 1.29 – 1.19 (m, 20H), 0.85 (t, *J* = 6.6 Hz, 3H).

### Copies of NMR and IR spectra

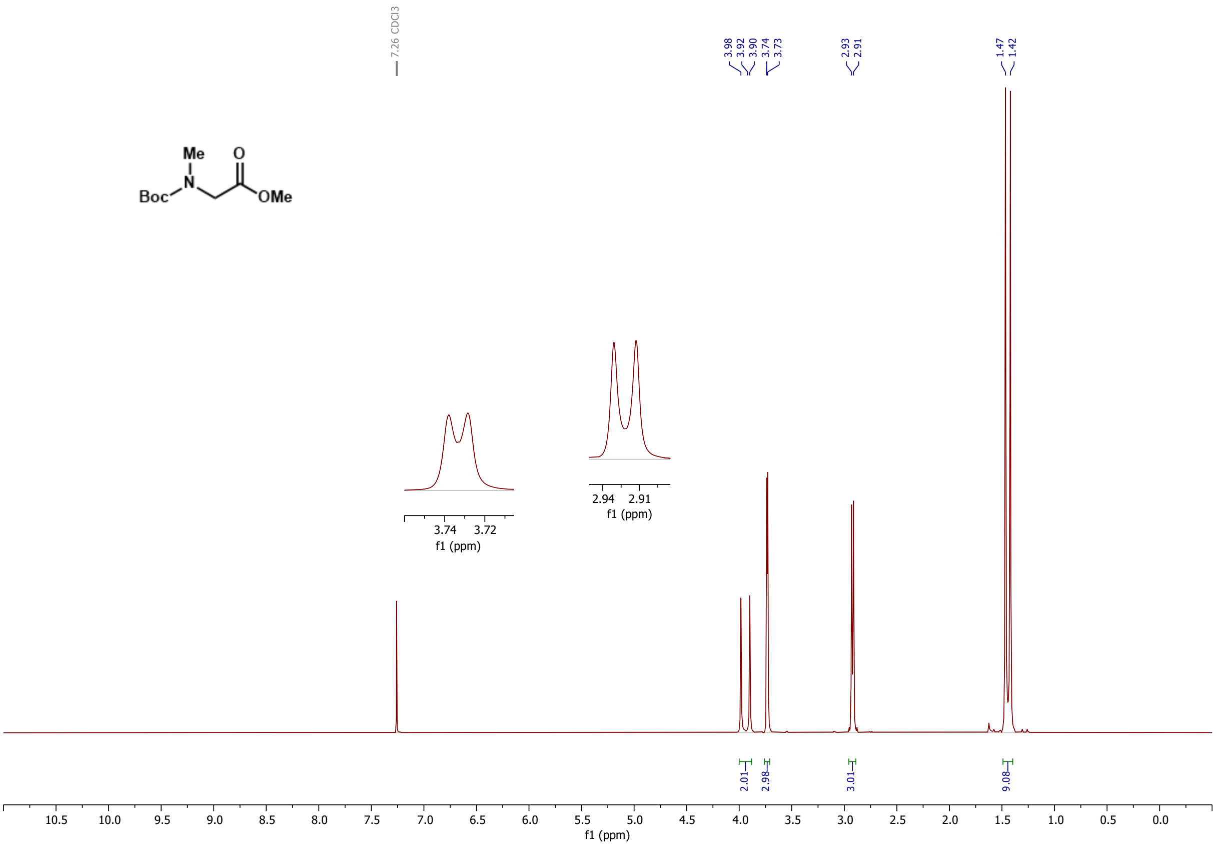


***Figure S15***: *$^{1}H$ NMR of compound* ***4*** *in $CDCl_3$*

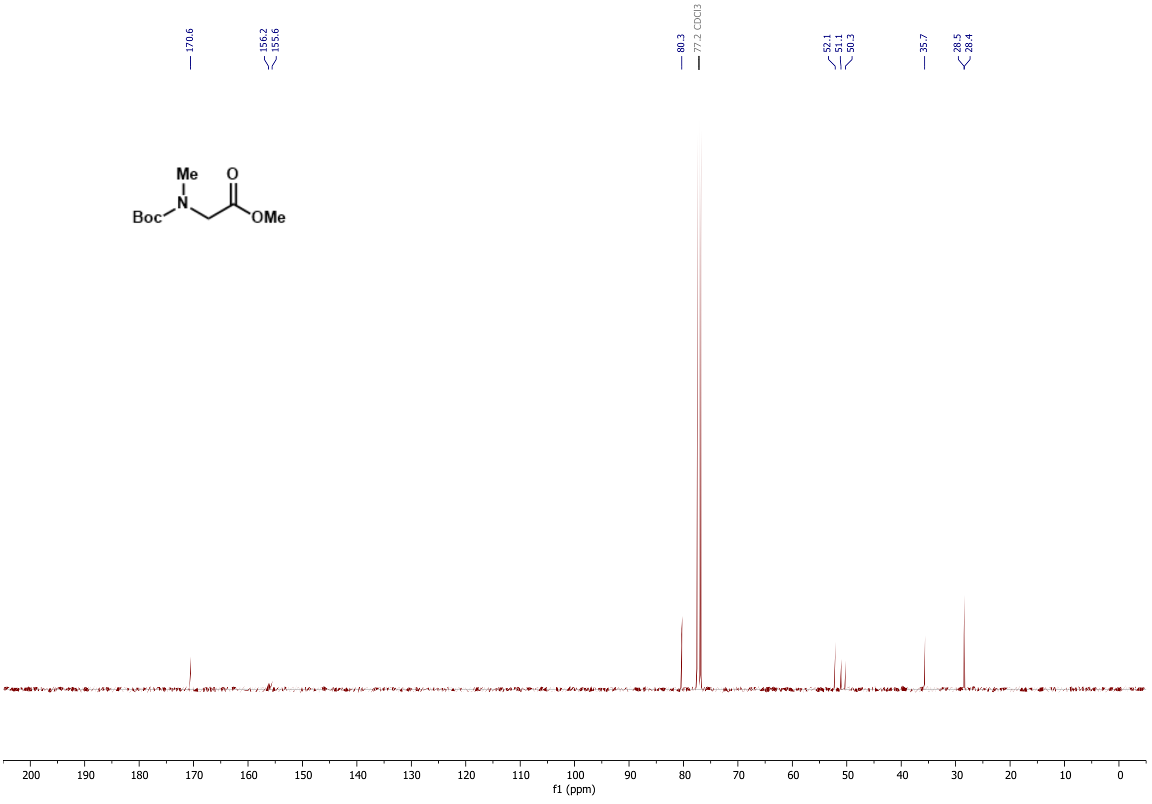

***Figure SS16**: $^{13}C$ NMR of compound **4** in $CDCl_3$*

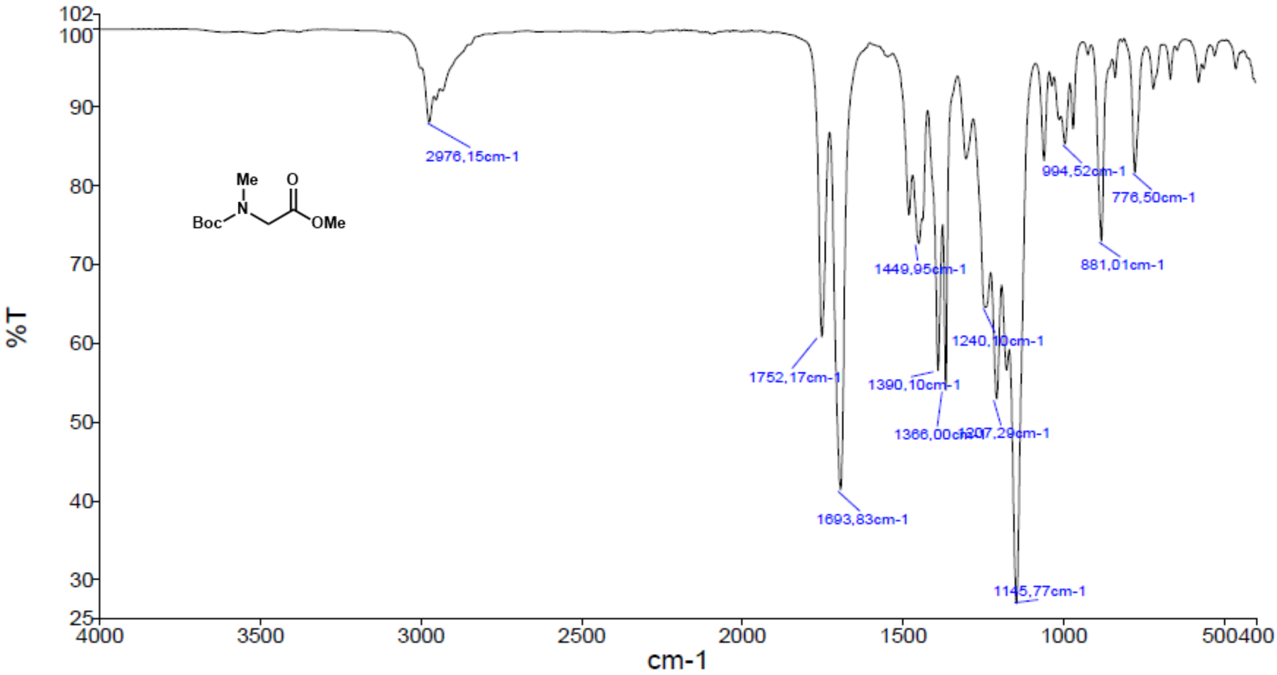


***Figure S17:** IR of compound **4***

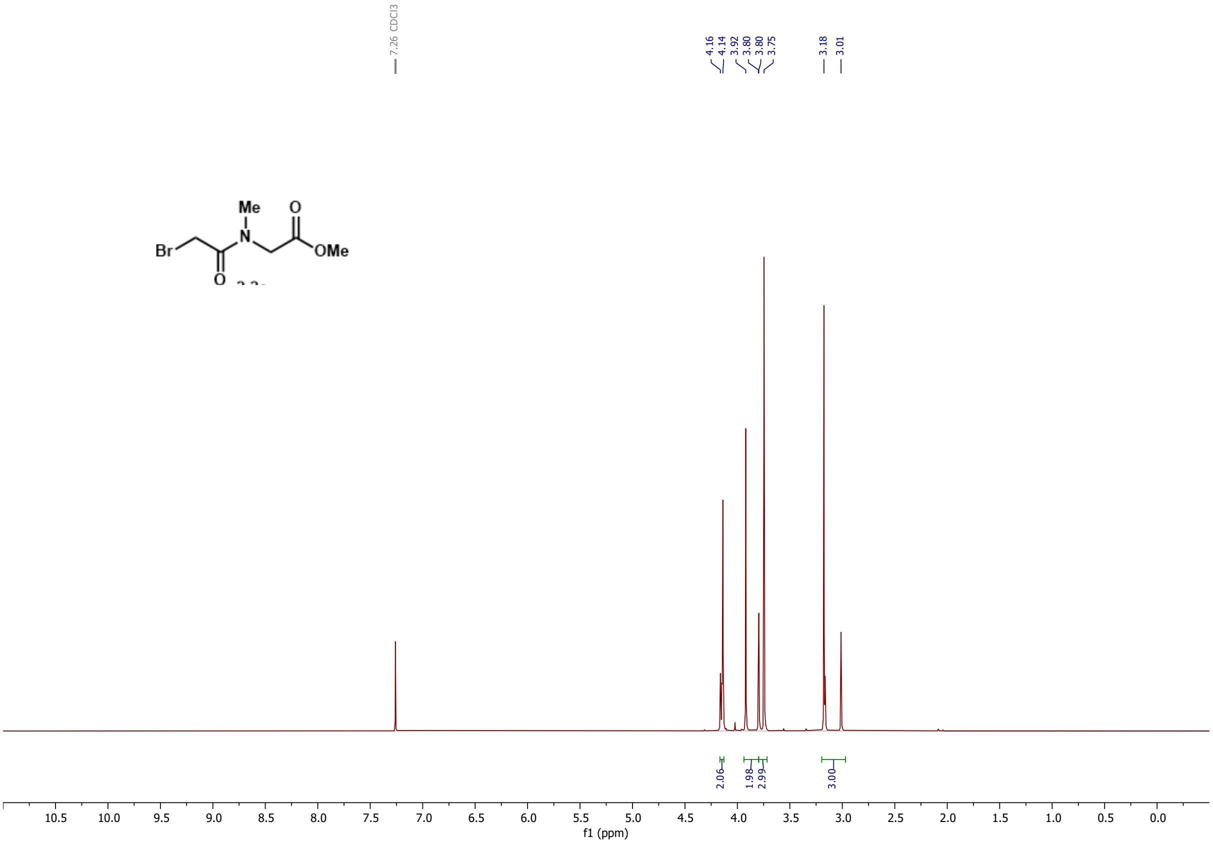


***Figure S18**: $^{1}H$ NMR of compound **6** in $CDCl_3$*

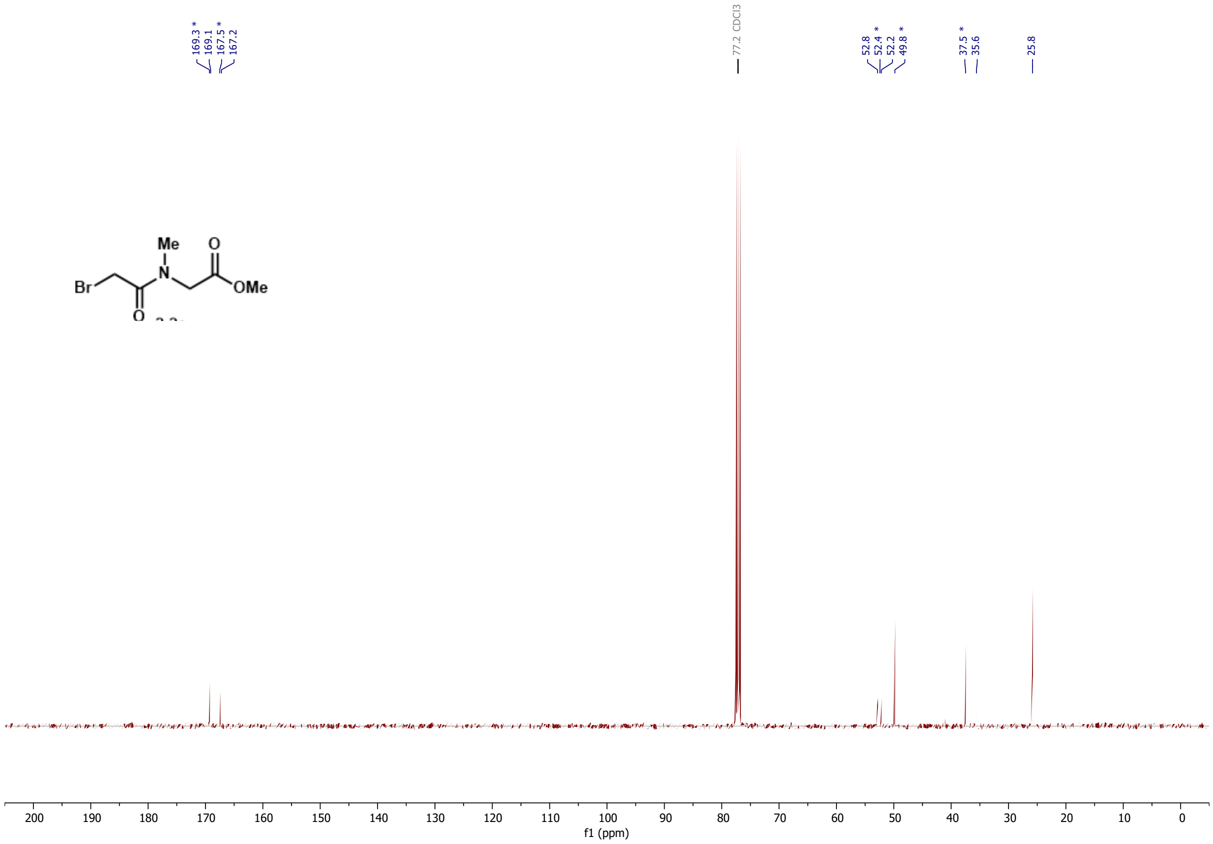


***Figure S19***: *$^{13}$C NMR of compound **6** in $CDCl_3$*

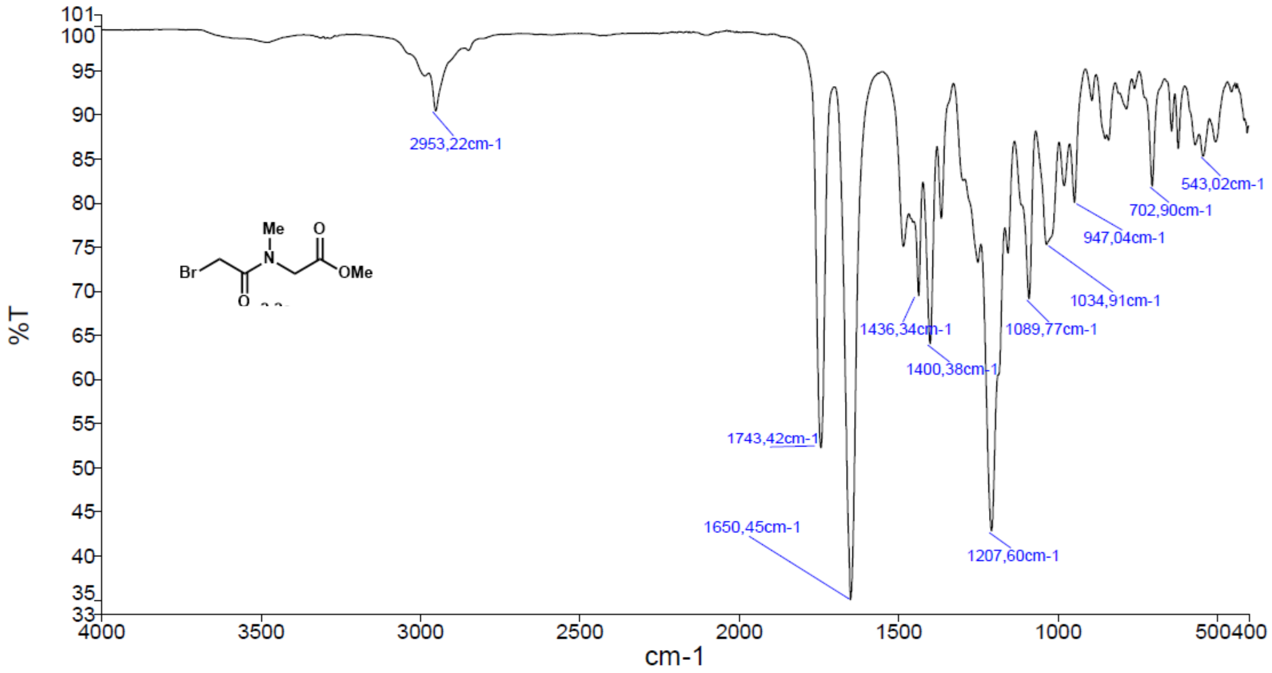


***Figure S20***: *IR of compound **6***

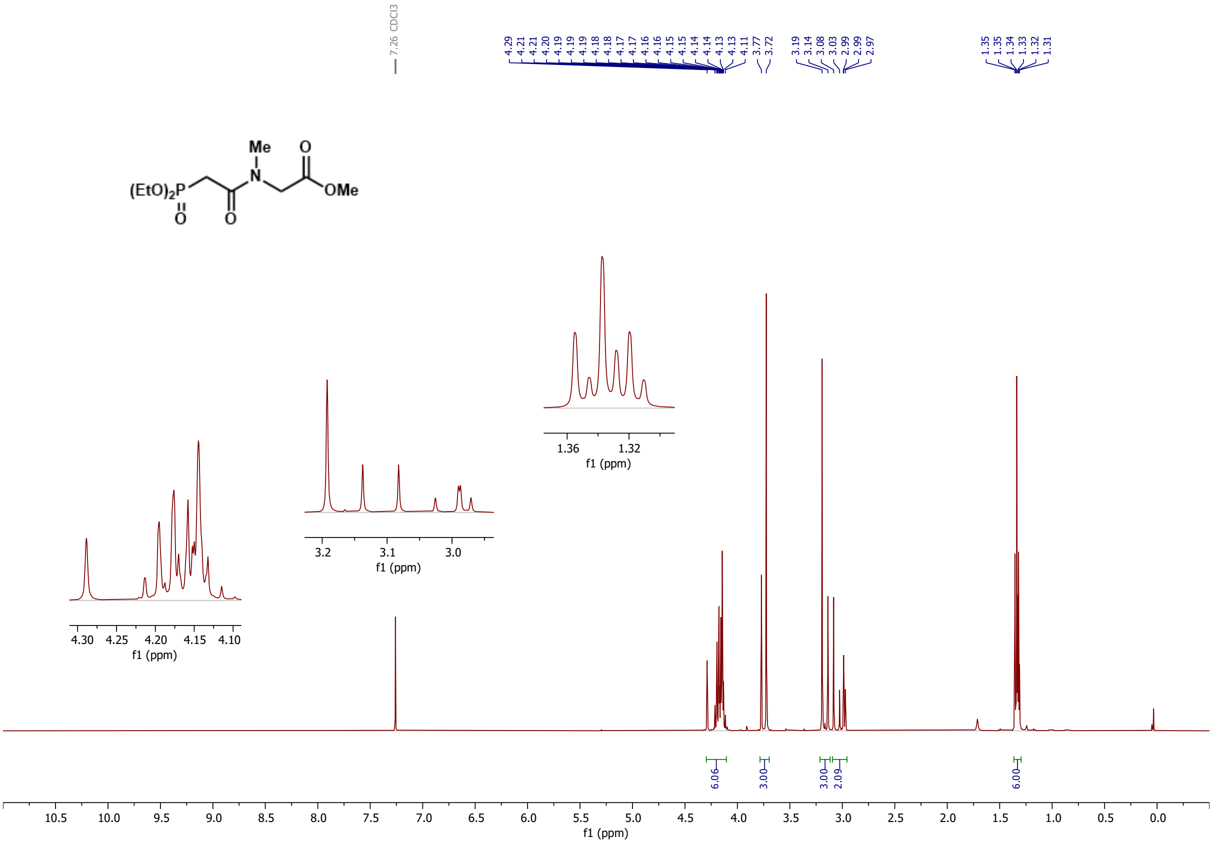

***Figure S21***: *[1]H NMR of compound **7** in $CDCl_3$*

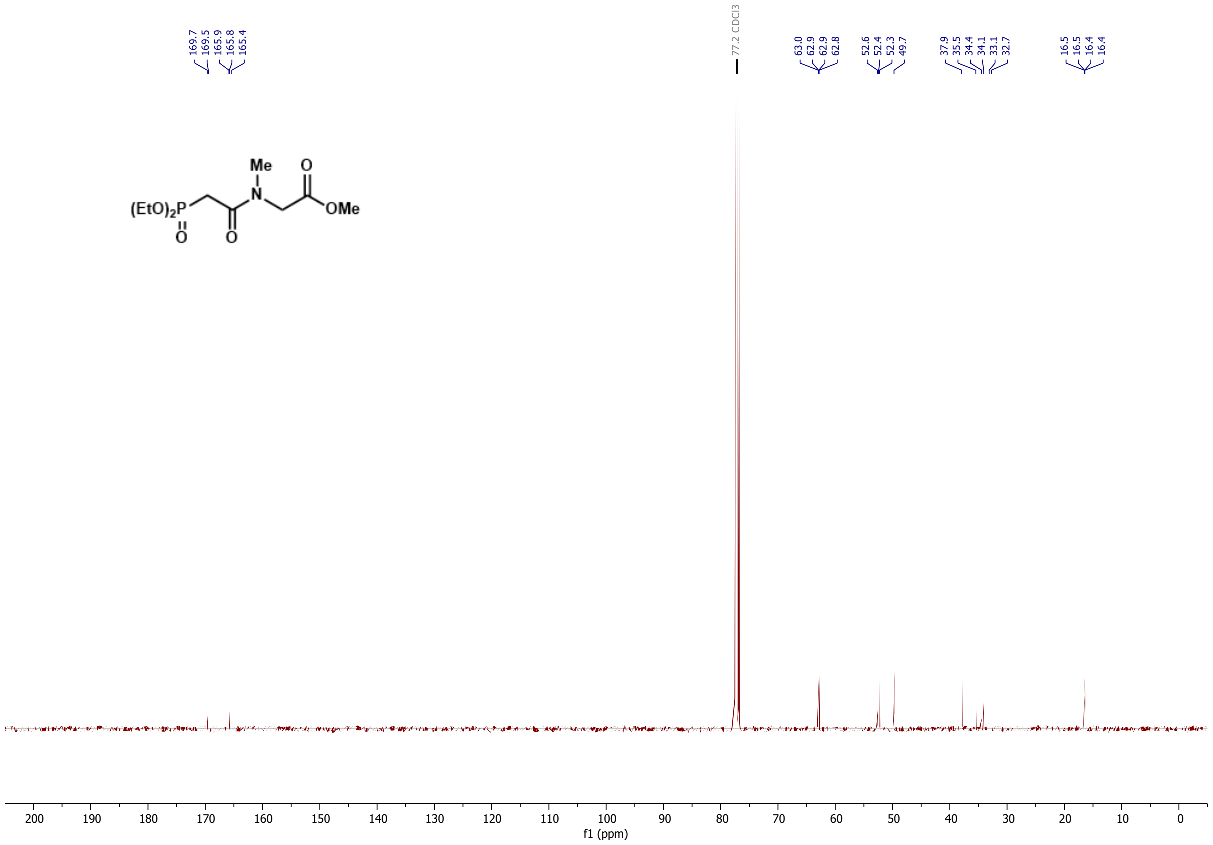

***Figure S22***: *[13]C NMR of compound **7** in $CDCl_3$*

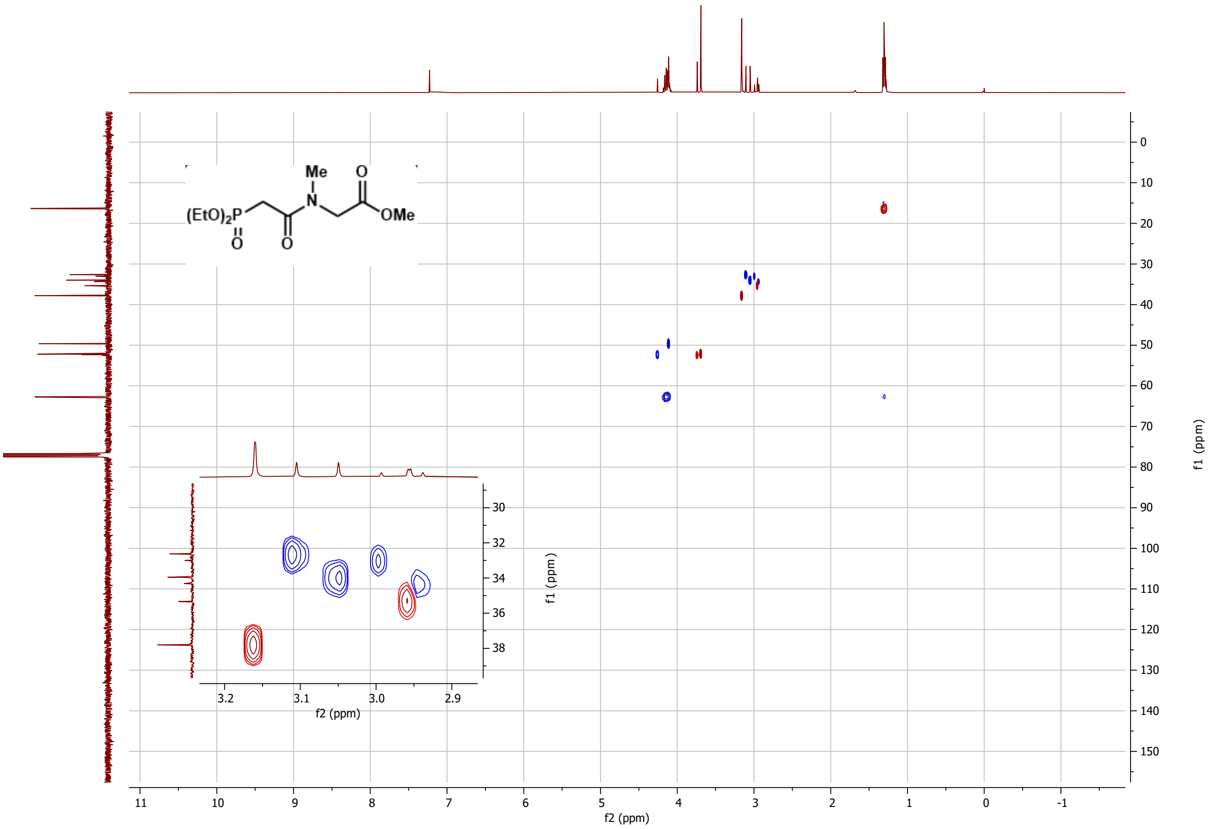


***Figure S23***: *[1]H-[13]C HSQC of compound **7** in $CDCl_3$*

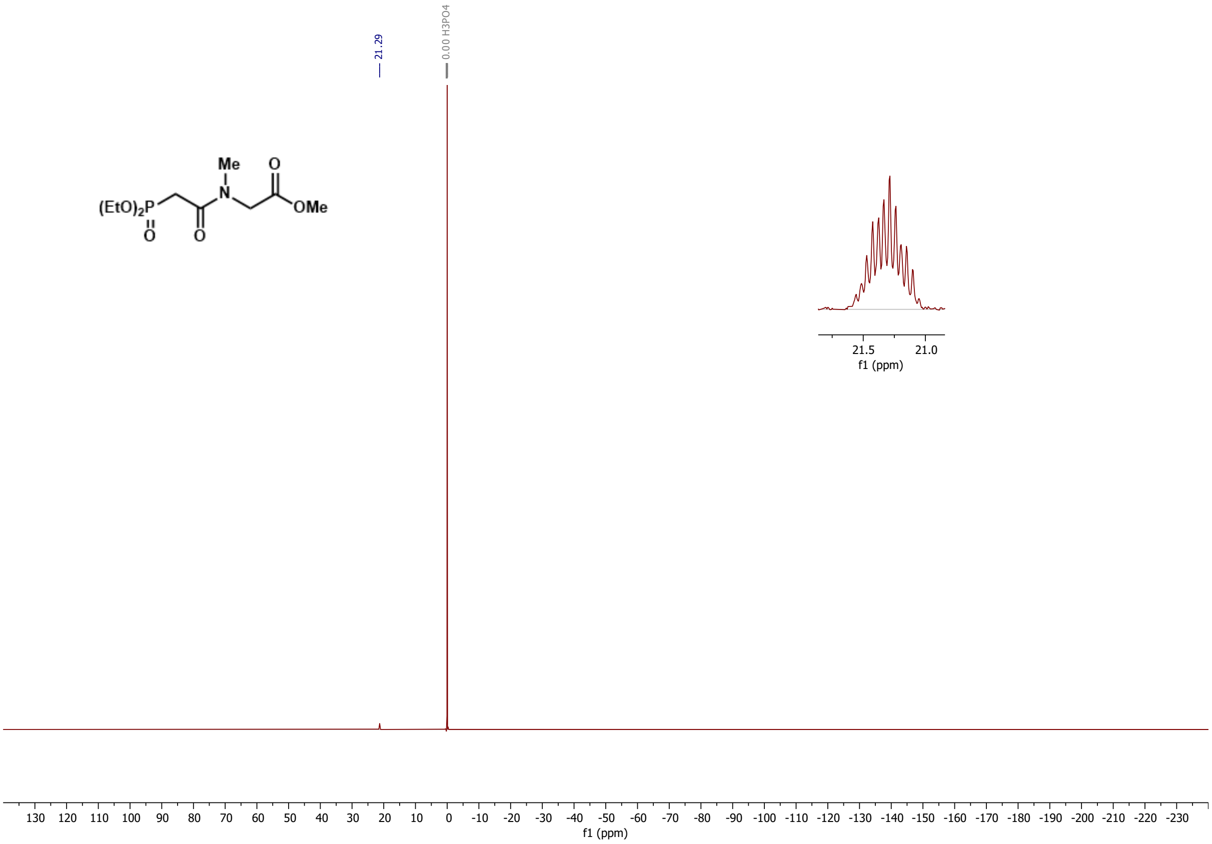


***Figure S24***: *[31]P NMR of compound **7** in $CDCl_3$*

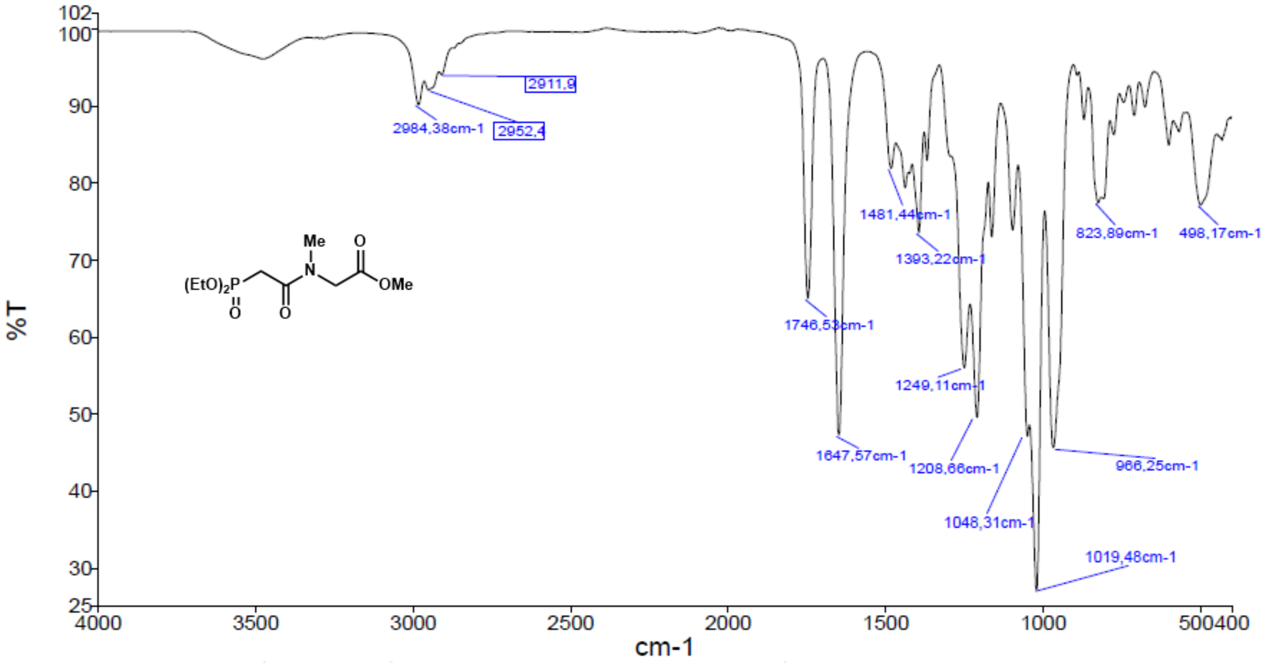


***Figure S25****: IR of compound* ***7***

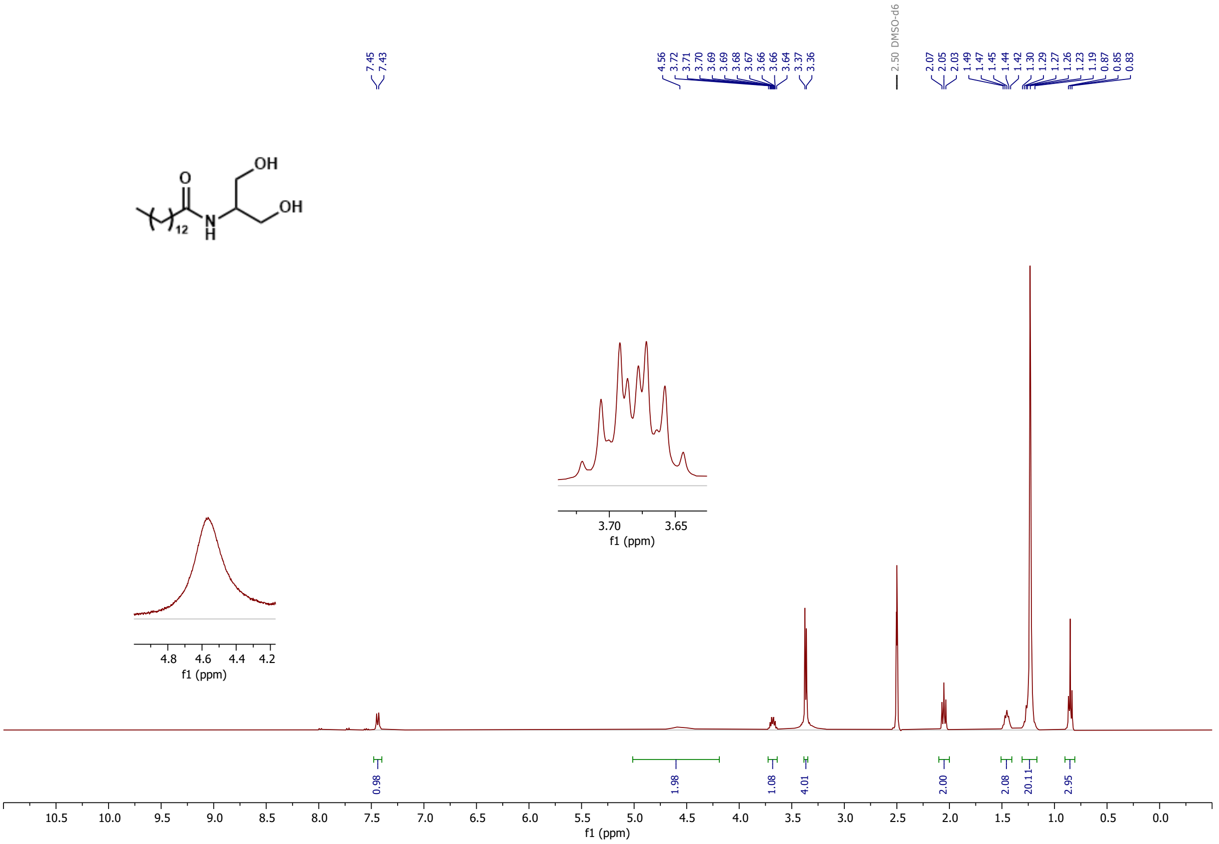


***Figure S26****: ¹H NMR of compound* ***9*** *in DMSO-d₆*

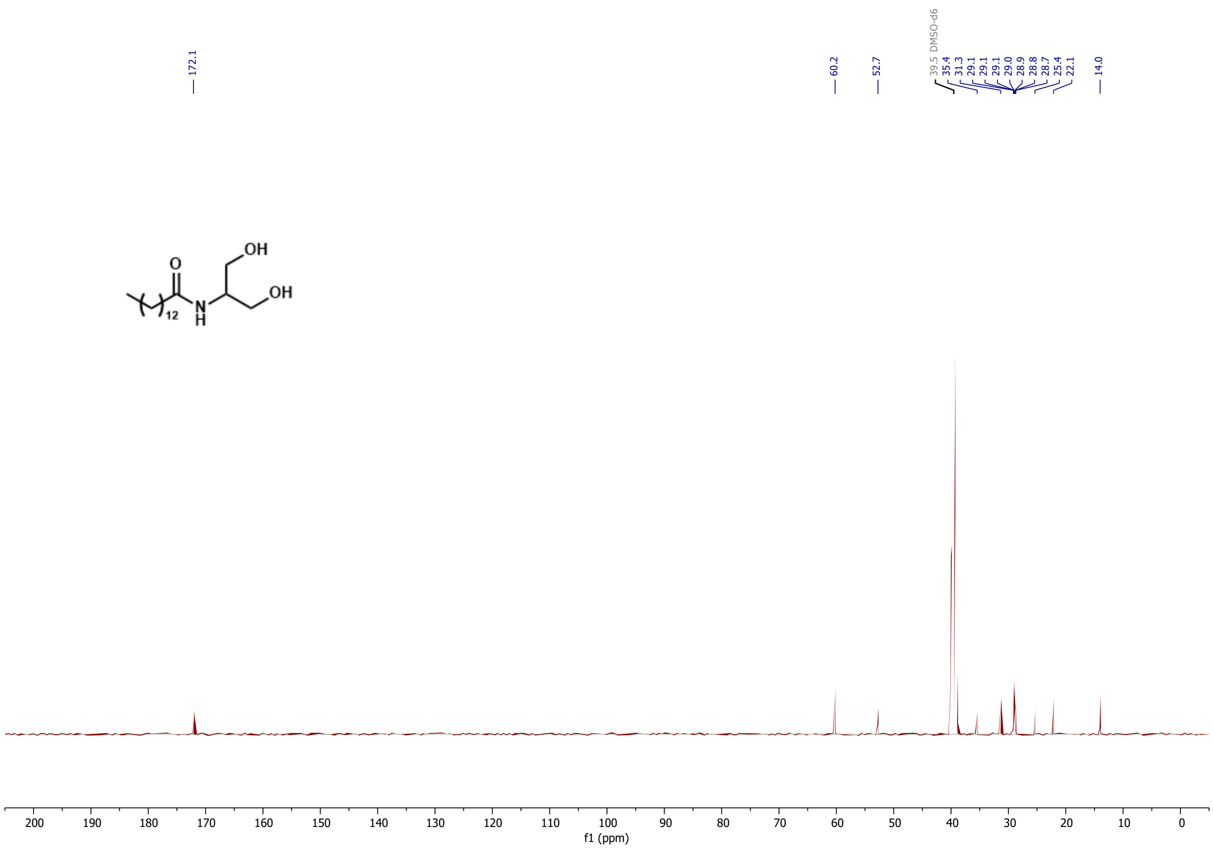


***Figure S27***: *$^{13}$C NMR of compound **9** in DMSO-$d_6$*

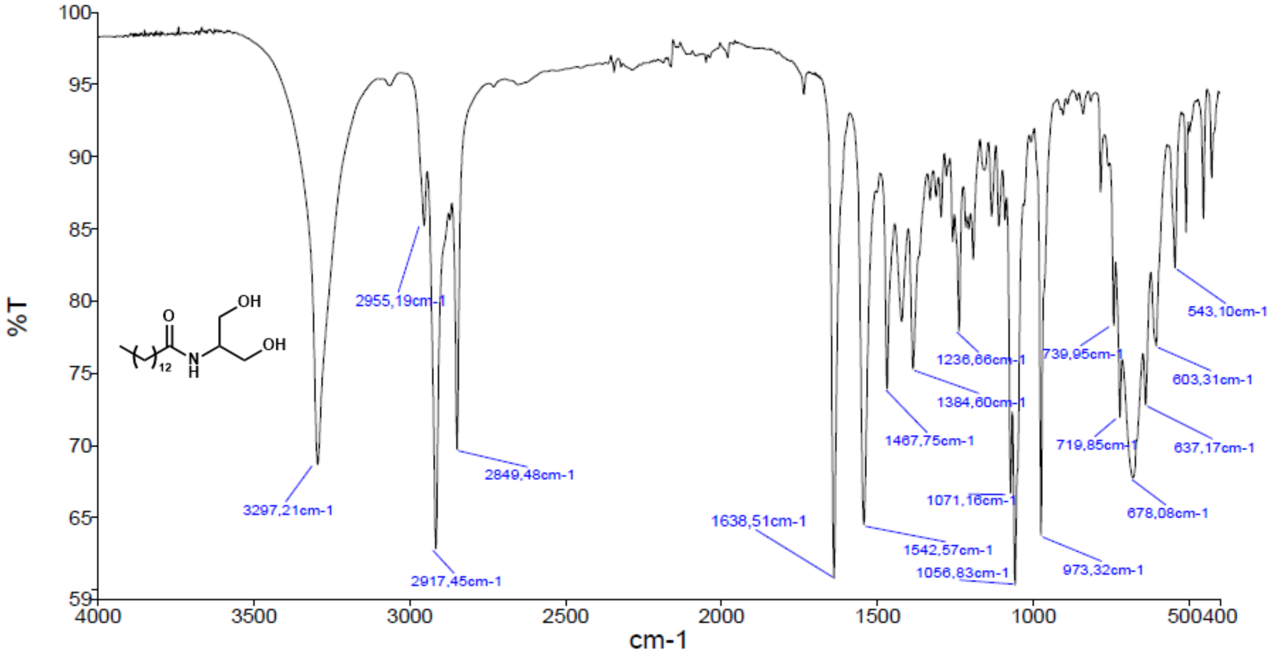


***Figure S28***: *IR of compound **9***

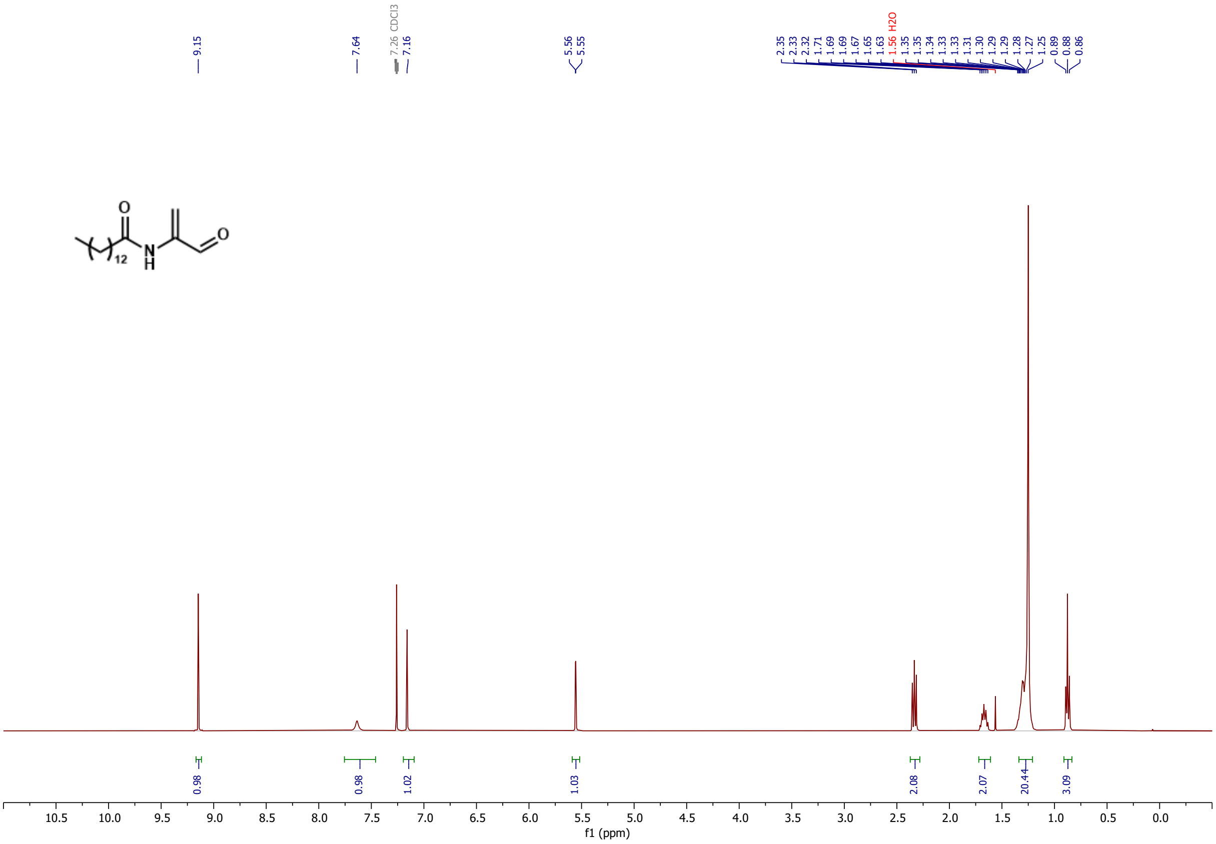

***Figure S29***: *[1]H NMR of compound **8** in $CDCl_3$*

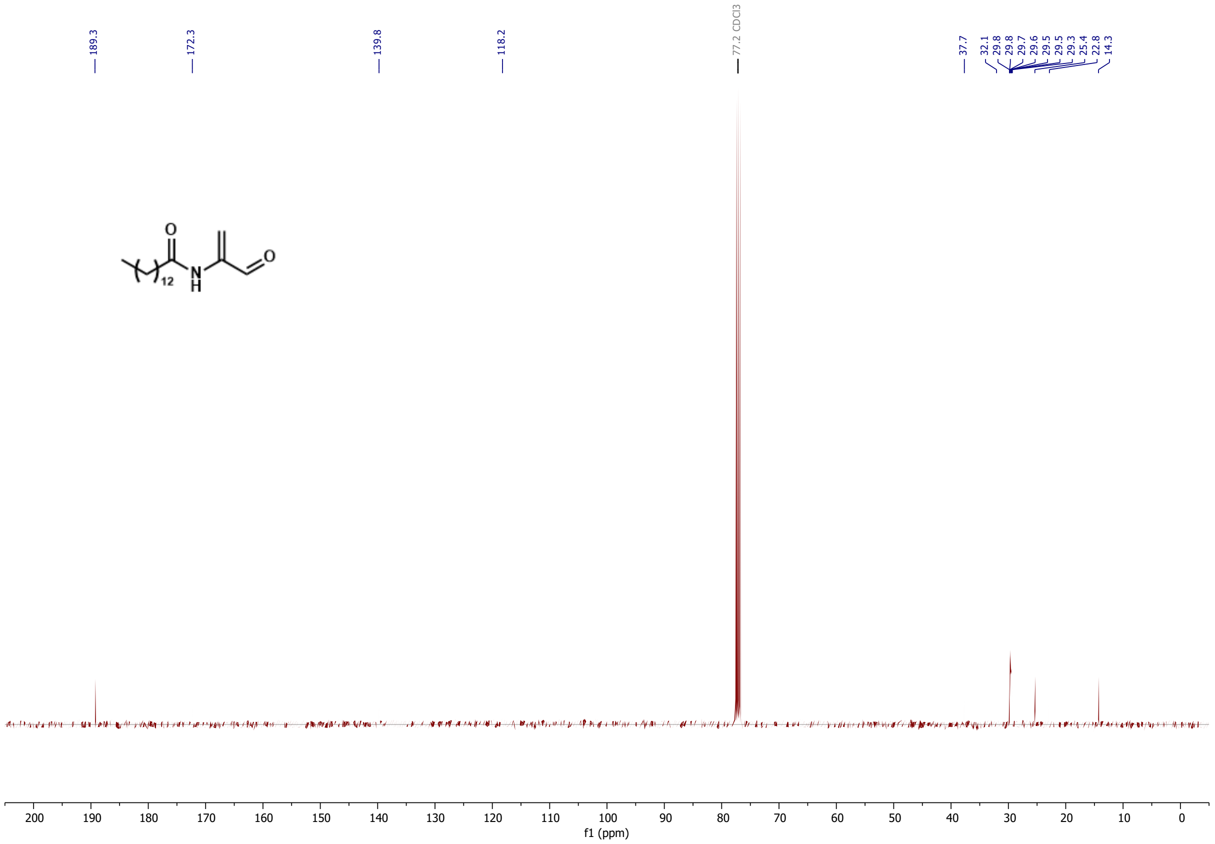

***Figure S30***: *[13]C NMR of compound **8** in $CDCl_3$*

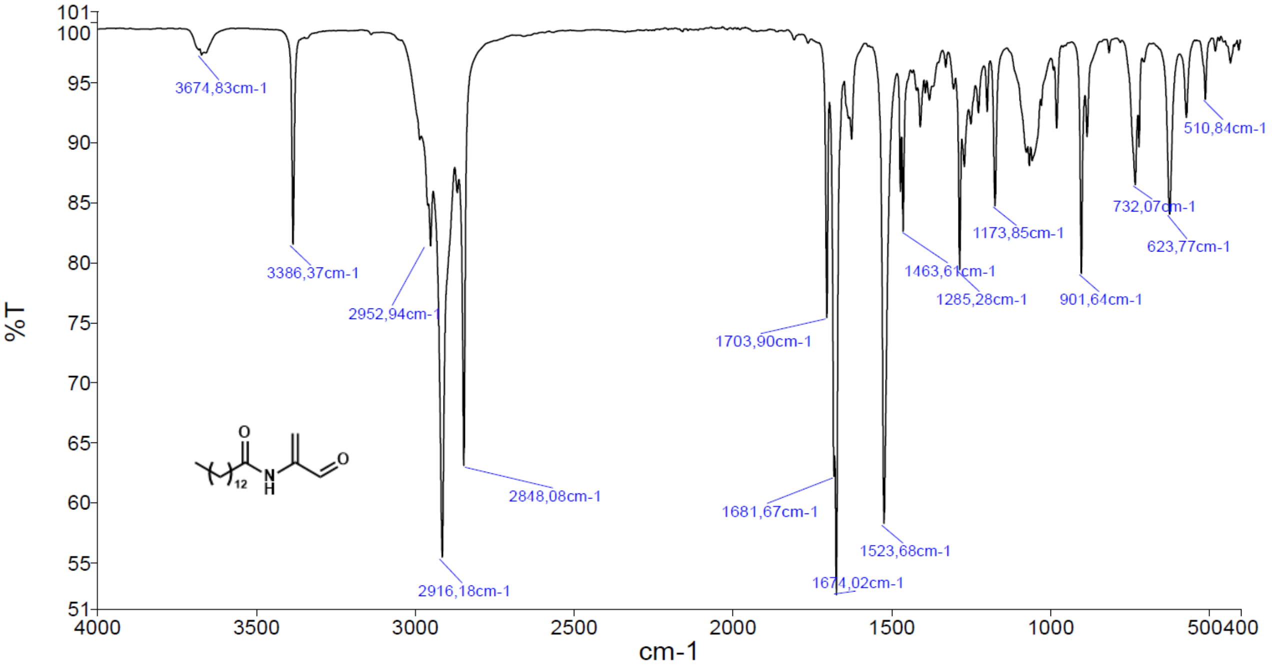


***Figure S31**: IR of compound **8***

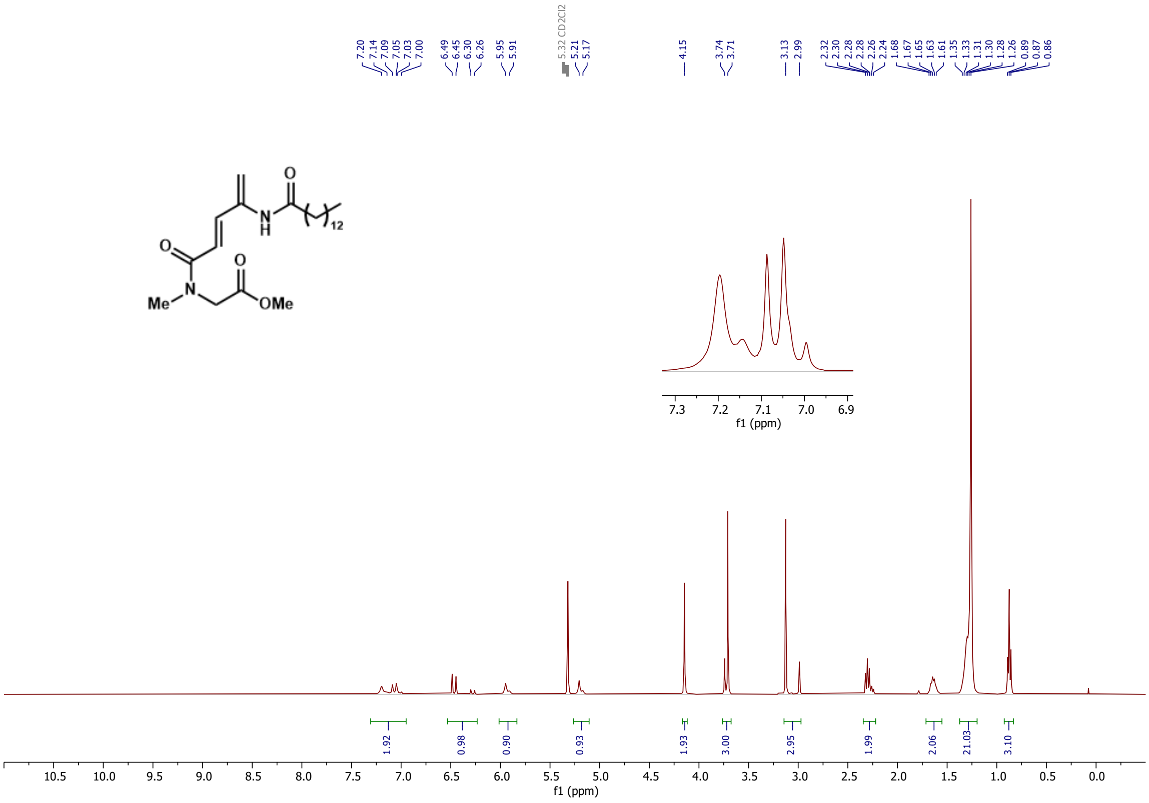


***Figure S32**: $^{1}H$ NMR of compound **10** in $CD_2Cl_2$*

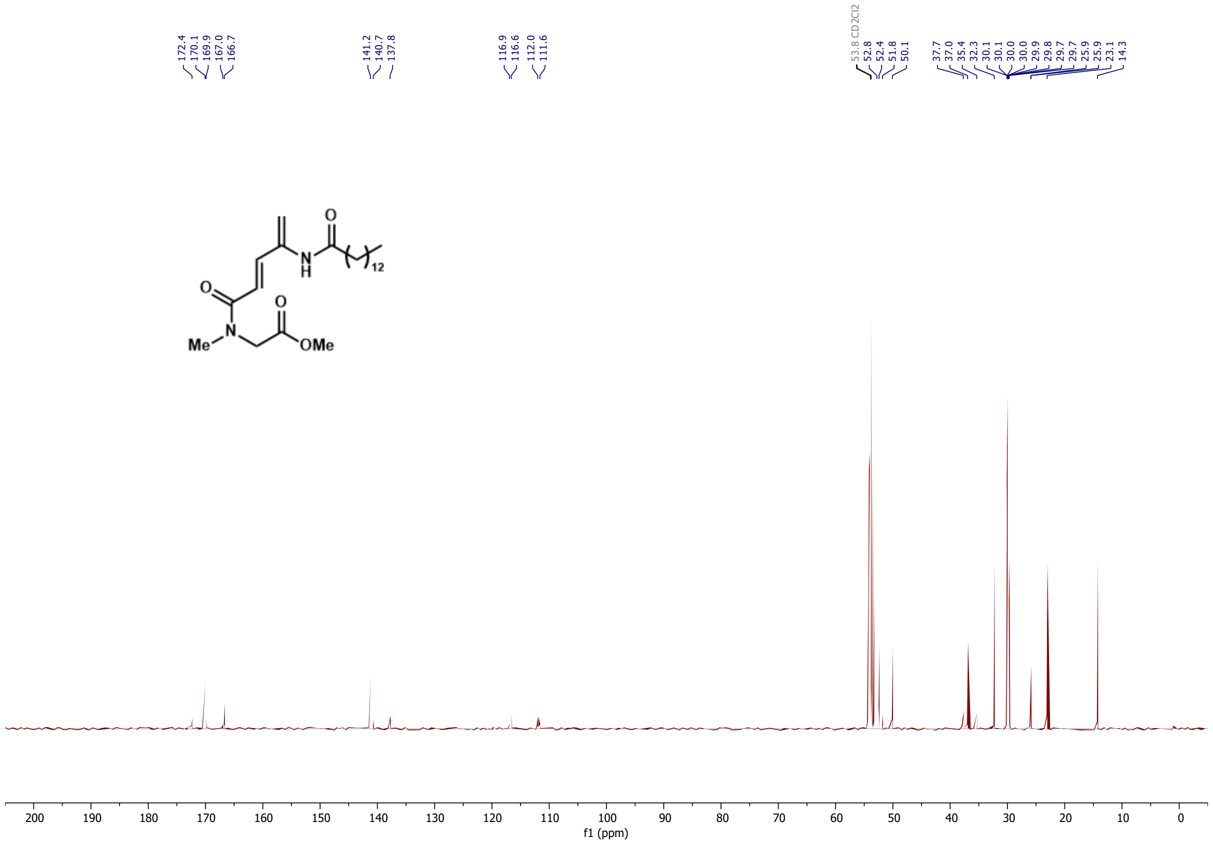

***Figure S33***: *[13]C NMR of compound* ***10*** *in* $CD_2Cl_2$

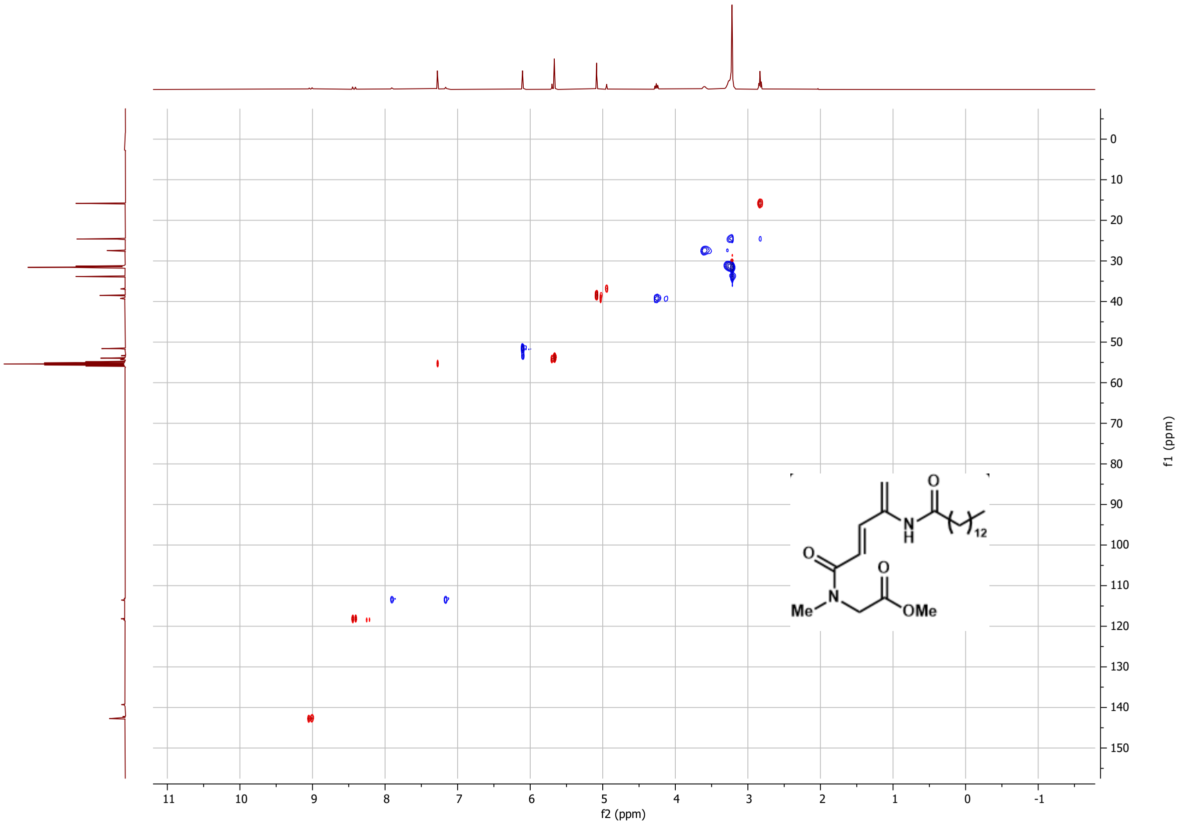

***Figure S34***: *[1]H-[13]C HSQC of compound* ***10*** *in* $CD_2Cl_2$

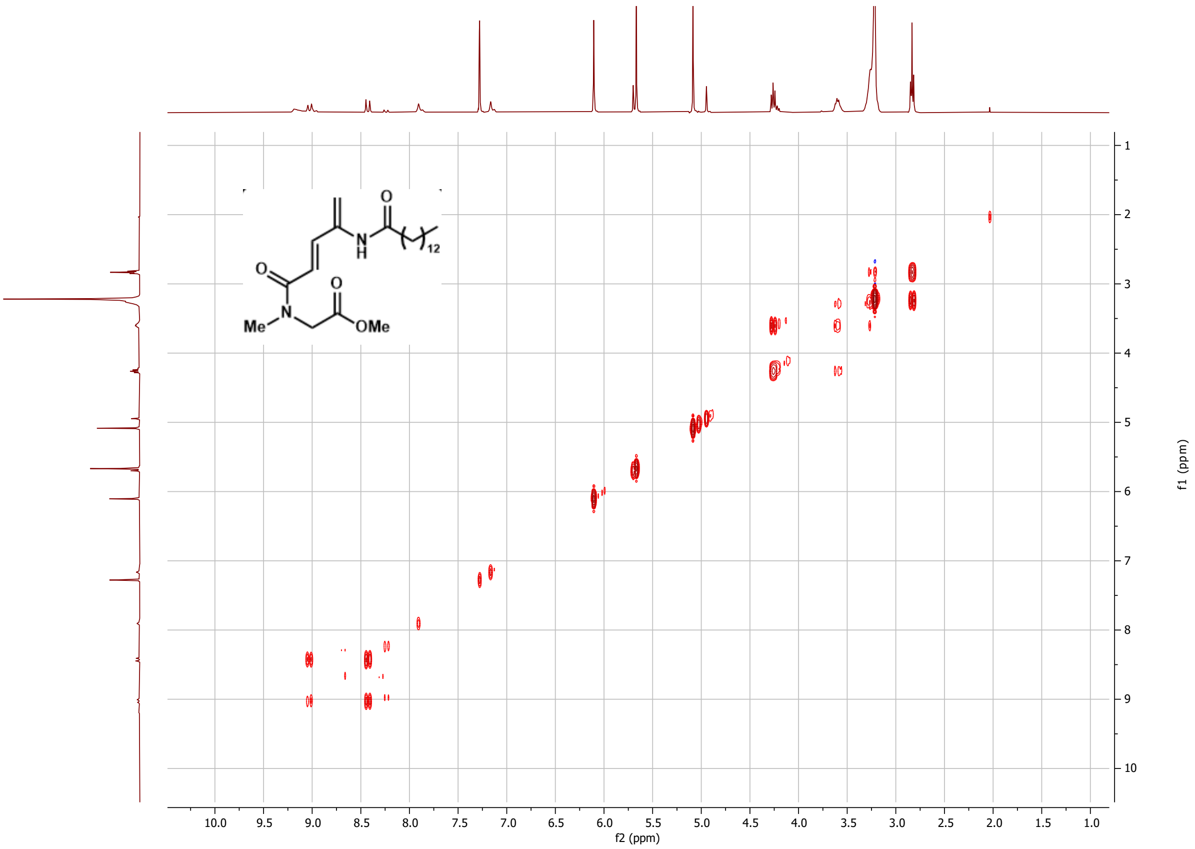


***Figure S35***: *$^1H$-$^1H$ COSY of compound **10** in $CD_2Cl_2$*

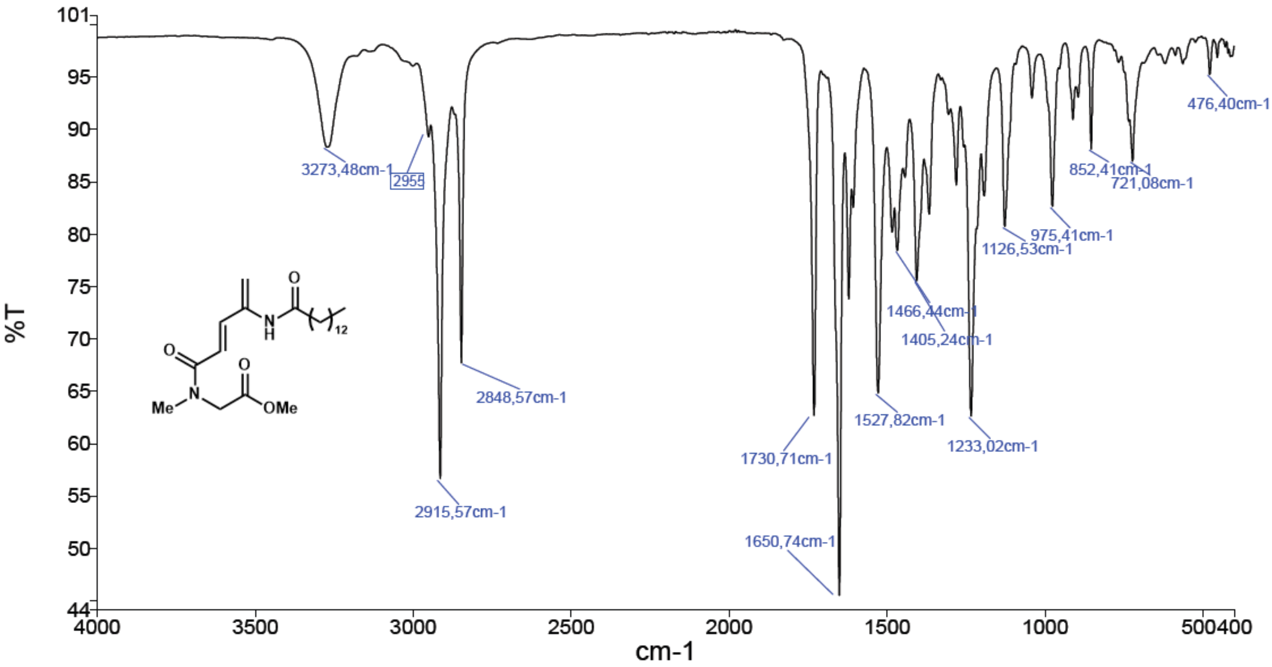


***Figure S36***: *IR of compound **10***

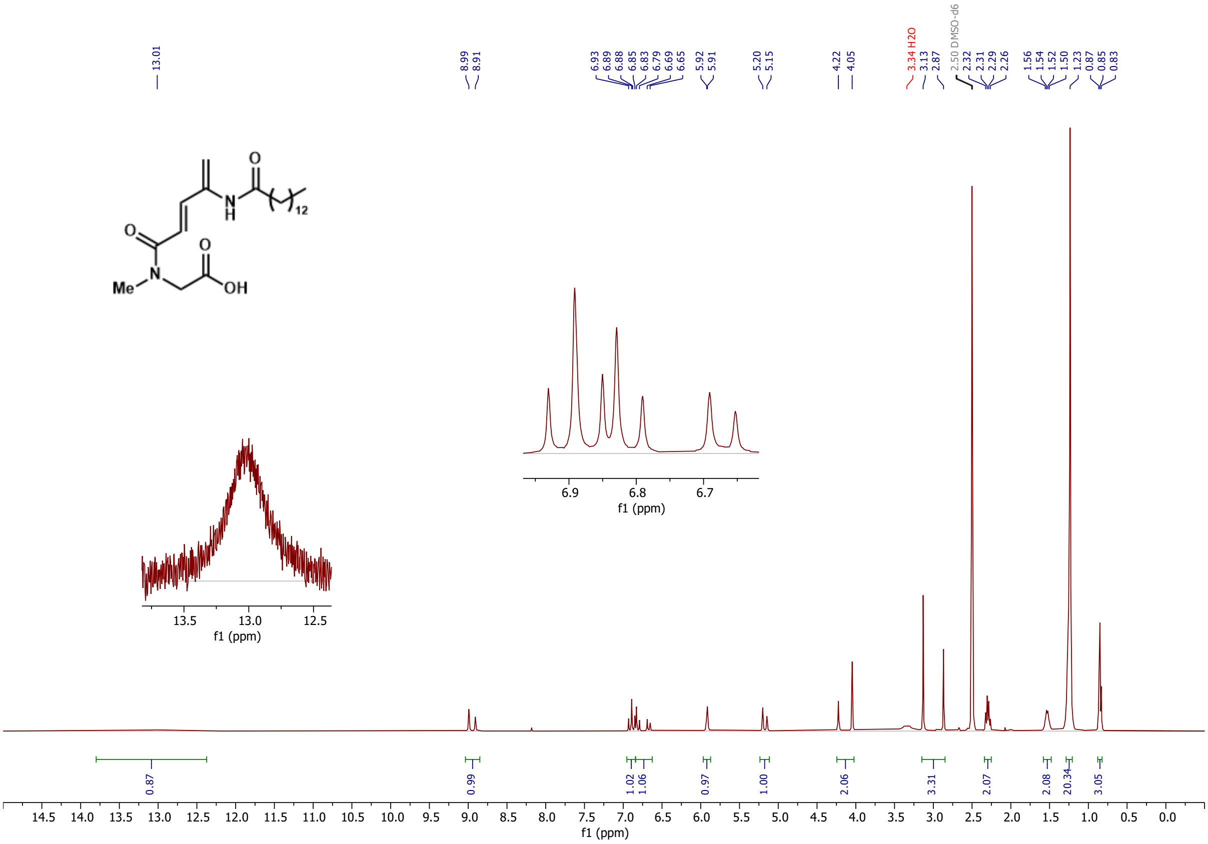

***Figure S37***: *¹H NMR of compound **3** in DMSO-d₆*

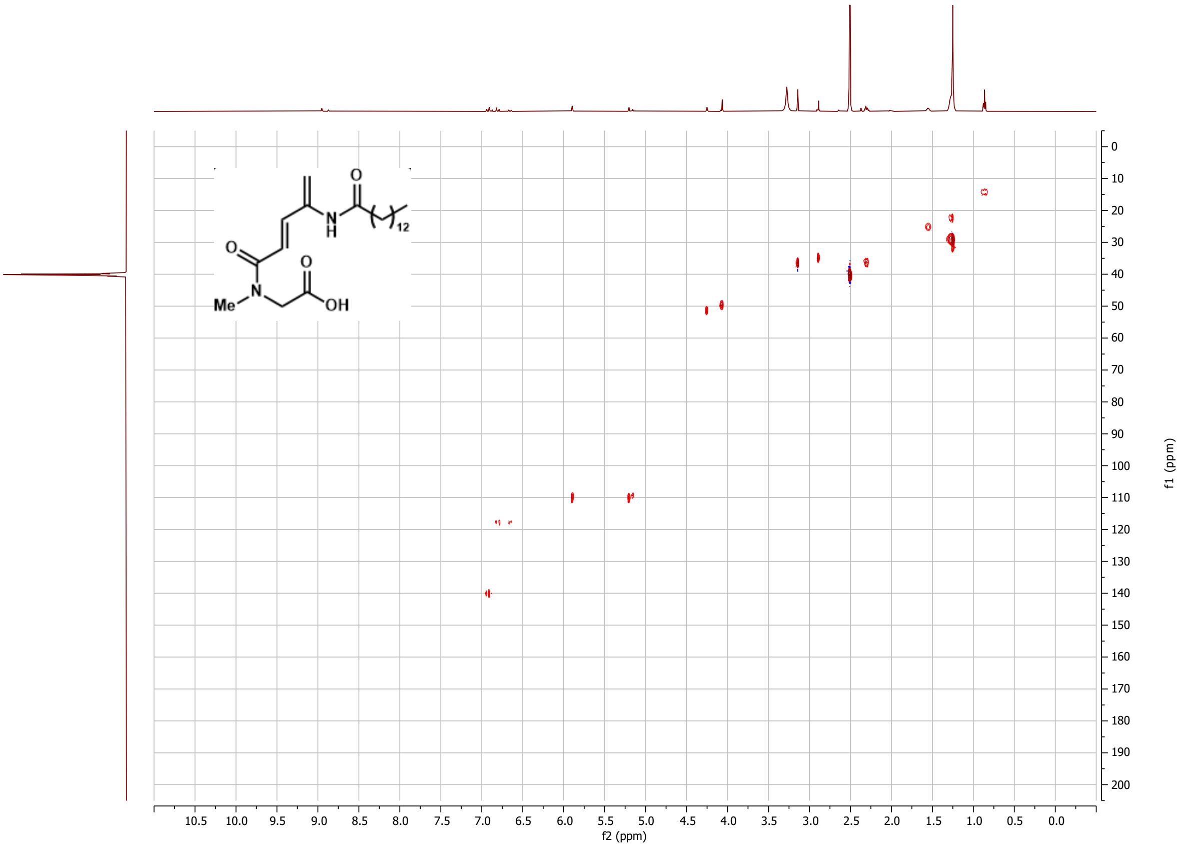

***Figure S38***: *¹H-¹³C HSQC of compound **3** in DMSO-d₆ (500 MHz)*

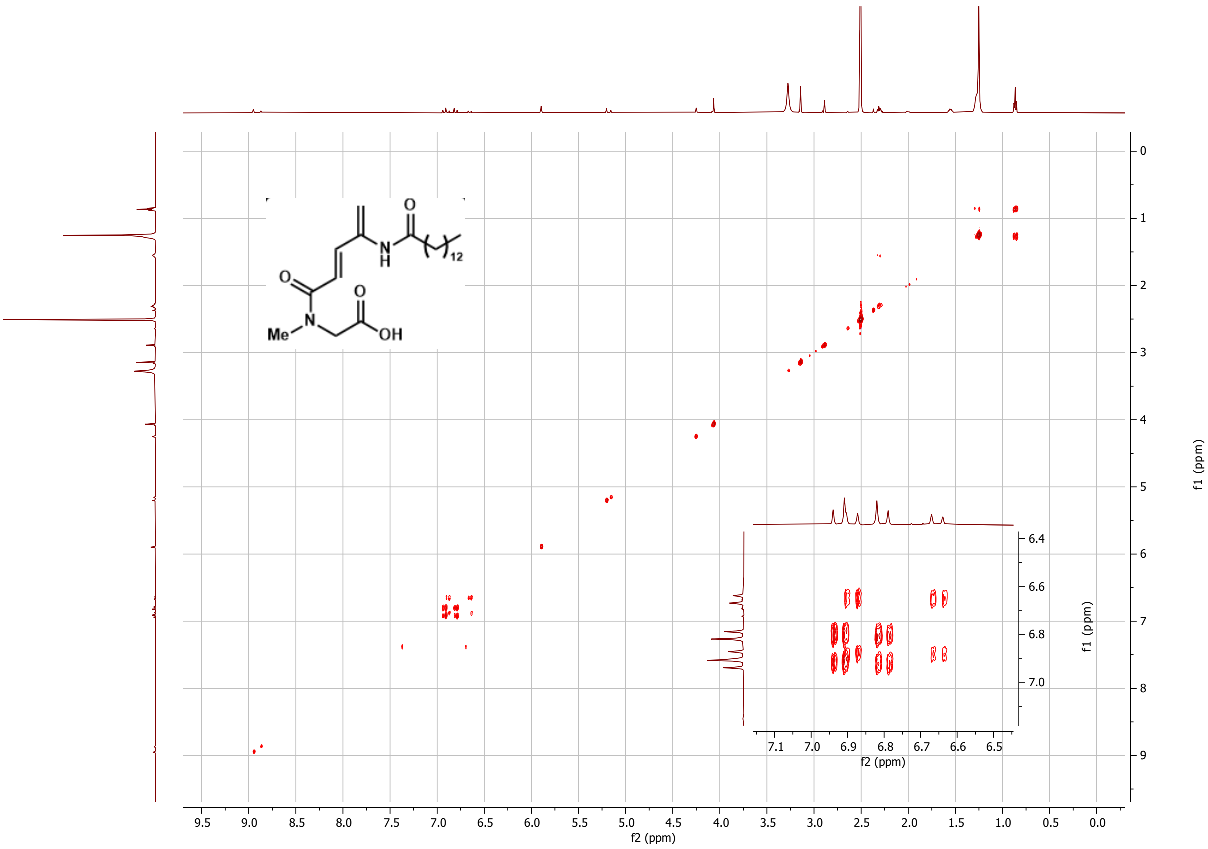


***Figure S39***: *$^1$H-$^1$H COSY of compound **3** in DMSO-$d_6$ (500 MHz)*
